# Evolutionary origins of sexual dimorphism : Lessons from female-limited mimicry in butterflies

**DOI:** 10.1101/2021.07.09.451774

**Authors:** Ludovic Maisonneuve, Charline Smadi, Violaine Llaurens

## Abstract

The striking female-limited mimicry observed in some butterfly species is a text-book example of sexually-dimorphic trait submitted to intense natural selection. Two main evolutionary hypotheses, based on natural and sexual selection respectively, have been proposed. Predation pressure favouring mimicry toward defended species could be higher in females because of their slower flight, and thus overcome developmental constraints favouring the ancestral trait that limits the evolution of mimicry in males but not in females. Alternatively, the evolution of mimicry in males could be limited by females preference for non-mimetic males. However, the evolutionary origin of female preference for non-mimetic males remains unclear. Here, we hypothesise that costly sexual interactions between individuals from distinct sympatric species might intensify because of mimicry, therefore promoting female preference for non-mimetic trait. Using a mathematical model, we compare the evolution of female-limited mimicry when assuming either alternative selective hypotheses. We show that the patterns of divergence of male and female trait from the ancestral traits can differ between these selection regimes. We specifically highlight that divergence in females trait is not a signature of the effect of natural selection. Our results also evidence why female-limited mimicry is more frequently observed in Batesian mimics.

## Introduction

The evolutionary forces involved in the emergence of sexual dimorphism in different animal species are still debated. As highlighted by Wallace [1865], divergent natural selection could drive the evolution of strikingly different phenotypes in males and females, because they may occupy different ecological niches. Sexual selection exerted by females is also a powerful force leading to the emergence of elaborated traits in males only, therefore leading to sexual dimorphism [Darwin, 1871]. The relative contributions of natural and sexual selections to the evolution of sexually dimorphic traits has generated important controversies. The evolution of sexual dimorphism in wing colour patterns in butterflies has been central to this debate because wing colour patterns are under strong natural selection by predators and are also involved in mate choice and species recognition [Turner, 1978]. Quantifying phenotypic divergence in males and females from the ancestral trait may allow one to identify the main evolutionary factors involved in the evolution of sexual dimorphism. Using a phylogenetic approach on European butterflies, van der Bijl et al. [2020] recently showed that the wing colour pattern dimorphism is mainly driven by the divergence of male phenotype from the ancestral trait, in line with the sexual selection hypothesis. In contrast to this general trend, sexual dimorphism where females exhibit a derived colour pattern is frequently observed in butterfly species involved in Batesian mimicry [Kunte, 2008]. In these palatable species, the evolution of colour patterns looking similar to the phenotype displayed in chemically-defended species living in sympatry is strongly promoted: because predators associate conspicuous colouration to defences, individuals displaying mimetic colouration in palatable species have a reduced predation risk [Bates, 1981, Ruxton et al., 2019]. Despite predation affecting individuals from both sexes, mimicry is sometimes surprisingly limited to females [Ford, 1975, Kunte, 2008, Long et al., 2014, Nishikawa et al., 2015], therefore begging the question of the evolutionary forces preventing the evolution of mimicry in males (*i.e*. female-limited mimicry, named FLM hereafter).

Because butterfly males and females generally differ in their behaviour, the strength of predation pressure might differ among sexes [Ohsaki, 1995, 2005]: for instance, females usually spend a lot of time ovipositing on specific host-plants, and thus have a more predictable behaviour for predators. Moreover, flight speed is generally higher in males than females: females are heavier because they carry eggs [Gilchrist, 1990], and males have higher relative thorax mass [Karlsson and Wickman, 1990] and muscle mass [Marden and Chai, 1991], resulting in increased flight power [Chai and Srygley, 1990]. Predation pressures are thus expected to be stronger in females. In line with this expectation Su et al. [2015] show that in sexually monomorphic mimetic butterflies females are more prefect mimics than males, suggesting also that some constraints limits perfect mimicry in males. Wing pattern evolution is also shaped by developmental constraints [Van Belleghem et al., 2020] that may impede divergence from the ancestral trait [Maisonneuve et al., 2021]. Phylogenetic analyses show that FLM derived from sexually monomorphic non-mimetic ancestors [Kunte, 2009, Timmermans et al., 2017] suggesting that mimicry in FLM species is associated with a costly displacement from an ancestral non-mimetic phenotype. In the female-limited polymorphic butterfly *Papilio polytes*, where both mimetic and non-mimetic females co-exist, the mimetic allele reduces the pre-adult survival rate [Komata et al., 2020, Katoh et al., 2020] (but see [Komata et al., 2018] in the FLM butterfly *Papilio memnon*), highlighting cost associated with mimicry. Such trade-off between developmental constraints favouring the ancestral trait and selection promoting mimicry might differ between sexes: if predation is lower in males, the constraints limiting mimicry may overcome the benefit from mimicry in males, whereas in females the higher predation pressure may promote mimicry. In line with this idea, in mimetic Asian pitvipers, where males suffer for a greater predation pressure, females are rarely mimetic, strengthening the role of sexually contrasted predation in promoting sex-limited mimicry [Sanders et al., 2006]. Nevertheless, evidence for the limited predation in males as compared to females is controversial in butterflies [Wourms and Wasserman, 1985] therefore questioning whether contrasted predation in males and females is actually the main driver of FLM.

Other constraints triggered by sexual selection might limit mimicry in males. In the female-limited Batesian mimic *Papilio polyxenes asterius*, experimental alteration of male colour pattern into female colour pattern leads to lower success during male-male encounters and increased difficulty in establishing a territory, therefore reducing mating opportunities [Lederhouse and Scriber, 1996]. Furthermore, in the female-limited Batesian mimic *Papilio glaucus*, females prefer control painted non-mimetic males over painted mimetic males [Krebs and West, 1988] (but see [Low and Monteiro, 2018] in the FLM butterfly *Papilio polytes*). Wing colour patterns in mimetic butterflies may therefore modulate male reproductive success, by influencing both male-male competition and mating success with females. In particular, females preference for ancestral trait may generate sexual selection limiting male mimicry [Belt, 1874., Turner, 1978]. Nevertheless, because mimetic colouration is under strong positive selection, females are predicted to prefer mimetic males because it leads to adapted mimetic offspring, favouring mimetic colouration in males, as observed in species involved in Müllerian mimicry, *i.e*. when co-mimetic species are all chemically-defended [Jiggins et al., 2001, Naisbit et al., 2001, Kronforst et al., 2006, Merrill et al., 2014]. It is thus unclear what does limit the evolution of females preference towards mimetic colouration in males from mimetic species.

Females preference for mimetic males may be disadvantageous because this behaviour may lead to mating interactions with unpalatable ’model’ species. Therefore reproductive interference, *i.e*. costly interactions between different species during mate acquisition (see [Gröning and Hochkirch, 2008] for a definition), may impair the evolution of females preference towards mimetic colour patterns displayed by other sympatric species. The evolution of mimetic colouration in males may indeed increase costs linked to reproductive interference in females, and therefore promote the evolution of preference for non-mimetic traits in males. Such reproductive interference has been observed between species sharing similar aposematic traits (in *Heliconius* and *Mechanitis* species [Estrada and Jiggins, 2008]). The rate of erroneous mating may be limited by the difference in male pheromones between mimetic species (see Darragh et al. [2017], González-Rojas et al. [2020] for empirical examples in *Heliconius* butterflies). However, females may still suffer from cost associated to reproductive interference, even if they refuse mating with heterospecific males: females may allow courting by heterospecific males displaying their preferred cue, resulting in increased investment in mate searching (see signal jamming in [Gröning and Hochkirch, 2008]). Pheromones may not limit this increase of investment in mate searching, because they act as short-distance cue that may be perceived only during the courtship [Mérot et al., 2015]. Females deceived by the colour pattern then need to deploy substantial efforts to avoid the heterospecific mating.

Theoretical studies highlight that the reproductive interference between sympatric species influence the evolution of traits used as mating cues. Reproductive interference indeed promotes the evolution of females preference towards traits differing from the phenotype displayed in other sympatric species, because it reduces the number of costly sexual interactions [McPeek and Gavrilets, 2006, Yamaguchi and Iwasa, 2013, Maisonneuve et al., 2021]. However these studies do not consider the independent evolution male and female traits. Under weak constraint on sex differentiation, reproductive interference may impede divergence of male trait, while natural selection may promote the evolution of female trait, leading to sexual dimorphism. For instance, in two of the three fruit fly species of the genus *Blepharoneura* that court on the same host plant, a morphometric analysis reveals sexual dimorphism in wing shape where males, but not females, from the two different species differ in wing shape Marsteller et al. [2009]. In the mexican spadefoot toads *Spea multiplicata*, the level of sexual size dimorphism increases with the proportion of species from the same genus *Spea bombifrons* living in sympatry [Pfennig and Pfennig, 2005] suggesting a link between species interactions and sexual dimorphism. In species exhibiting FLM, reproductive interference may thus inhibit natural selection in males, while females become mimetic. Theoretical studies show that reproductive interference can totally impair the evolution of mimicry [Boussens-Dumon and Llaurens, 2021] or lead to imperfect mimicry [Maisonneuve et al., 2021] therefore suggesting that reproductive interference might indeed be a relevant ecological interaction preventing mimicry in males. In the model investigating the effect of reproductive interference on mimicry described in Boussens-Dumon and Llaurens [2021], colour-patten based assortative mating was assumed, preventing the study of the evolution of disassortative preferences in females. Therefore understanding the impact of reproductive interference on the evolution of FLM requires to specifically explore the evolution of female preference, and to assume a genetic architecture enabling mating cues to evolve in different directions in males and females.

Interestingly, the two main hypotheses usually explaining FLM, *i.e*. (1) sexually contrasted predation and (2) sexual selection on males, are both equally relevant for palatable, as well as unpalatable mimetic species. Indeed, sympatric unpalatable species frequently display a common mimetic trait [Sherratt, 2008], suggesting a strong selection promoting mimicry. However, FLM is considered to be widespread in palatable species but rare in unpalatable ones [Mallet and Joron, 1999] (but see [Nishida, 2017]). This suggests that the evolution of sexual dimorphism in mimetic species might depend on the level of defences.

Here, we investigate how (1) reproductive interference and (2) sexually contrasted predation may promote the evolution of FLM, using a mathematical model. Firstly we pinpoint the specific evolutionary outcomes associated with the emergence of FLM driven by reproductive interference or sexually contrasted predation, therefore providing relevant predictions for comparisons with empirical data. Secondly, we study the impact of unpalatability levels on the emergence of sexual dimorphism, to test whether FLM may be restricted to palatable species. Our model describes the evolution of quantitative traits, following the framework established by Lande and Arnold [1985] in a *focal* species, living in sympatry with a defended *model* species exhibiting a fixed warning trait. We specifically study the evolution of (1) the quantitative traits displayed in males *t_m_* and females *t_f_* involved in mimetic interactions, (2) the preference of females for the different values of males trait *p_f_*. We assume that individuals in the *focal* species gain protection against predators from the similarity of their warning trait towards the trait displayed by the unpalatable *model* species. However, trait similarity between species generates fitness costs of reproductive interference paid by females from the *focal* species [McPeek and Gavrilets, 2006, Yamaguchi and Iwasa, 2013]. We assume that a mating between individuals from the *focal* and the *model* species never produce any viable hybrid. We also consider constraints limiting mimicry promoting the ancestral trait value in the *focal* species, by assuming selection promoting the ancestral trait value *t_a_*. Using a weak selection approximation [Barton and Turelli, 1991, Kirkpatrick et al., 2002], we obtain equations describing the evolution of the mean trait and preference values. We then use numerical analyses to investigate (1) the role of reproductive interference in FLM and (2) the effect of the level of unpalatability in the *focal* species on the emergence of FLM.

## Model

We consider a single *focal* species living in sympatry with a defended species displaying a fixed warning trait (referred to as the *model* species hereafter). Within the *model* species, all individuals display the same warning trait. We investigate the evolution of the warning trait expressed in the *focal* species, influenced by both (1) predators behaviour promoting mimicry towards the *model* species and (2) mate choice exerted by females on the trait expressed by males. We assume that female is the choosy sex, implying an asymmetry in the selection pressure exerted on male and female traits, potentially favouring the emergence of a sexual dimorphism. We thus study the traits *t_m_* and *t_f_* expressed in males and females respectively, as well as the mate preference expressed by females towards males displaying trait value *p_f_*. In contrast, both males and females of the *model* species display traits close to the mean value 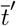, assumed to be fixed. Individuals of the *focal* species then benefit from increased survival when they display a trait similar to the trait expressed in the *model* species 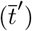, because of the learning behaviour of predators. This resemblance towards the *model* species then induces costs for individuals from the *focal* species, caused by reproductive interference. These reproductive interference costs depend on the discrimination capacities and mate preferences of females and on the phenotypic distances between (1) the traits displayed by males from the *focal* species and (2) the traits expressed in males from the *model* species.

We assume that the traits and preference in the *focal* species are quantitative traits, with an autosomal, polygenic basis with additive effects [Iwasa et al., 1991]. We assume that the distribution of additive effects at each locus is a multivariate Gaussian [Lande and Arnold, 1985]. We consider discrete and non-overlapping generations. Within each generation, natural selection acting on survival and sexual selection acting on reproductive success occur. Natural selection acting on an individual depends on the trait *t* expressed. We note 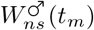 and 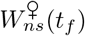 (defined after in equations (6) and (7)) the fitness components due to natural selection acting on a male of trait *t_m_* and a female of trait *t_f_* respectively. To compute the fitness component due to reproduction, we then note *W_r_*(*t_m_*, *p_f_*) (defined after in equation (21)) the contribution of a mating between a male with trait *t_m_* and a female with preference *p_f_* to the next generation. This quantity depends on (1) female mating preference, (2) male trait and (3) reproductive interference with the *model* species. The fitness of a mated pair of a male with trait *t_m_* and a female with trait *t_f_* and preference *p_f_* is given by:

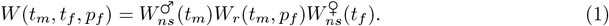

Using the Price’s theorem [Rice, 2004], we can approximate the change in the mean values of traits 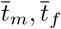 and preference 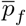 in the *focal* species after the natural and sexual selection respectively by:

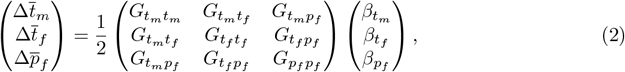

where for *i* ∈ {*t_m_*, *t_f_*, *p_f_*}, *G_ii_* is the genetic variance of *i* and for *i, j* ∈ {*t_m_*, *t_f_*, *p_f_*} with *i* ≠ *j G_ij_*, is the genetic covariance between *i* and *j* and with

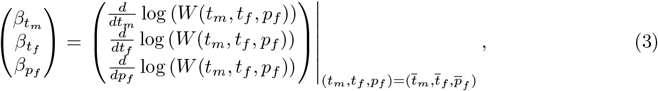

being the selection vector describing the effect of natural and sexual selections on mean traits and preference (see Appendix 1).

We assume weak natural and sexual selections [Iwasa et al., 1991, Pomiankowski and Iwasa, 1993], *i.e*. that the difference of fitness between different individuals is at maximum of order *ε*, with *ε* being small. Under this hypothesis genetic correlations generated by selection and non random mating quickly reach equilibrium [Nagylaki, 1993] and can thus be approximated by their equilibrium values. Weak selection hypothesis also implies that the variance of traits and preference is low [Iwasa et al., 1991].

Following [Iwasa et al., 1991], we assume that for *i* ∈ {*t_m_*, *t_f_*, *p_f_*}, *G_ii_* is a positive constant maintained by an equilibrium between selection and recurrent mutations. We assume *G_t_m_t_f__* to be constant: because neither selection nor nonrandom mating generate association between *t_m_* and *t_f_* this quantity depends only on the genetic architecture coding for traits expressed in males and females. For example *G_t_m_t_f__* = 0 would describe a situation where *t_m_* and *t_f_* are controlled by different sets of loci. Non-null value of *G_t_m_t_f__* would mean that *t_m_* and *t_f_* have (at least partially) a common genetic basis.

We assume that traits *t_m_* and *t_f_* have different genetic bases than preference *p_f_*. Thus only nonrandom mating generates genetic association between *t_m_* and *p_f_*. Under weak selection hypothesis *G_t_m_p_f__* is assumed to be at equilibrium. This quantity is given by (see Appendix 2):

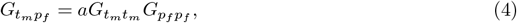

where *a* quantifies how frequently females reject males displaying non-preferred trait (see hereafter). Because neither selection nor nonrandom mating generate association between *t_f_* and *p_f_*, following equation (4a) in Lande and Arnold [1985], we have

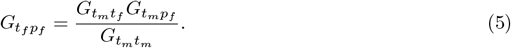

### Ancestral trait value *t_a_*

To investigate the effect of reproductive interference on the evolution of sexual dimorphism, we study the evolution of male and female traits (*t_m_* and *t_f_*) in the *focal* species, from an ancestral trait value initially shared between sexes (*t_a_*). This ancestral trait value *t_a_* represents the optimal trait value in the *focal* species, without interaction with the *model* species. This optimal value is assumed to be shaped by developmental as well as selective constraints, specific to the *focal* species. The natural selection exerted on males and females then depends on (1) departure from the ancestral trait value *t_a_*, inducing a selective cost *s*, as well as (2) protection against predators brought by mimicry, captured by the term 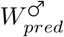 and 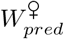 for males and females respectively. It is thus given by:

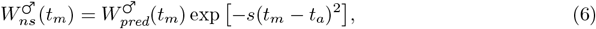

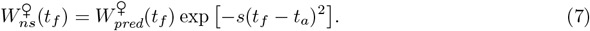

### Predation pressure exerted on warning trait

Predators exert a selection on individual trait promoting resemblance to the *model* species, resulting in an effect on fitness *W_pred_*. Müllerian mimicry indeed generates positive density-dependent selection [Benson, 1972, Mallet and Barton, 1989, Chouteau et al., 2016], due to predators learning. The density-dependence is modulated by the individual defence level λ, shaping predator deterrence: the higher the defence, the higher the defended individual contributes to the learning of predators. We note λ′ the defence level of an individual in the *model* species. We assume that harmless individuals (λ = 0) neither contribute to predators learning, nor impair it. The protection gained against predators then depends on the level of resemblance (as perceived by predators) among defended prey only, and on the number of defended individuals sharing the same signal. We note *N* and *N*′ the densities of individuals in the *focal* species and in the *model* species, respectively, and we assume a balanced sex ratio. The level of protection gained by an individual with trait *t* because of resemblance with other individuals is given by:

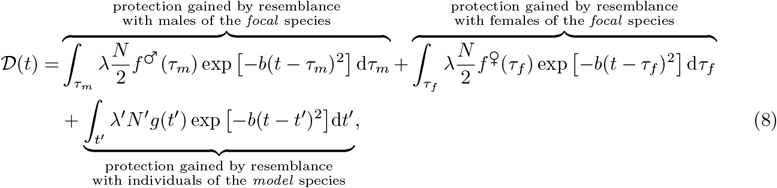

where exp [–*b*(*t* – *τ*)^2^] describes how much predators perceive the trait values *t* and *τ* as similar. The predators discrimination coefficient *b* thus quantifies how much predators discriminate different trait values displayed by prey. *f*^♂^, *f*^♀^ and *g* are the distribution of traits in males and females of the *focal* species and in the *model* species respectively.

Assuming that the distribution of traits has a low variance within both the *focal* and the *model* species leads to the following approximation (see Appendix 3):

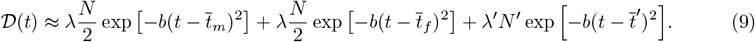

Because males and females can display different traits, the protection brought by mimicry might differ between sexes. Moreover, because males and females may have different behaviours and morphologies the strength of predation pressure can also vary between sexes. We note *d_m_*, *d_f_* ∈ (0, 1) the basal predation rates for males and females respectively. We assume these parameters to be of order *ε*, with *ε* small, ensuring that selection due to predation is weak (see Appendix 1 for analytical expression of selective coefficient). The impacts of predation on the fitness of a male and a female displaying the trait value *t_m_* and *t_f_* are given by:

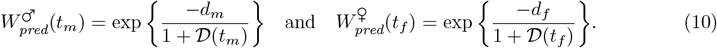

### Mating success modulating the evolution of female preference and male trait

The evolution of trait and preference also depends on the contribution to the next generation of crosses between males with trait *t_m_* and females with preference *p_f_*, *W_r_*(*t_m_*, *p_f_*). Because predators behaviour favours mimicry between sympatric species, substantial reproductive interference may occur in the *focal* species, because of erroneous species recognition during mate searching. Such reproductive interference depends on (1) females preference towards the warning trait displayed by males, (2) the distribution of this warning trait in males from both the *focal* and the *model* species and (3) the capacity of females to recognise conspecific males using alternative cues (pheromones for example). In the model, the investment of females in interspecific mating interaction is captured by the parameter *c_RI_* ∈ [0, 1]. This cost of reproductive interference incurred to the females can be reduced when female choice is also based on alternative cues differing between mimetic species. When a female with preference *p_f_* encounters a male displaying the trait value *t_m_*, the mating occurs with probability

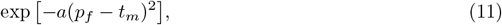

when the encountered male is a conspecific or

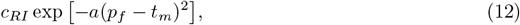

when the encountered male belongs to the *model* species. Females choosiness *a*, assumed constant among females, quantifies how frequently females reject males displaying a non-preferred trait.

During an encounter, the probability that a female with preference *p_f_* accepts a conspecific male is then given by [Otto et al., 2008]:

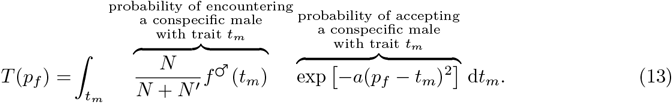

A female with preference *p_f_* may also accept an heterospecific male with probability:

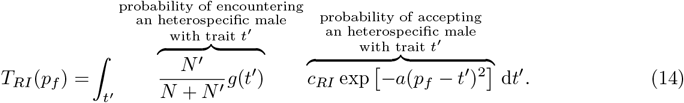

Assuming that the distribution of traits has a low variance within both the *focal* and the *model* species leads as before to the following approximations:

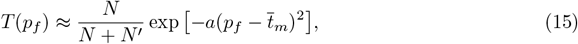

and

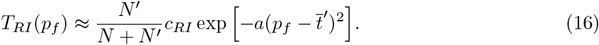

We assume that heterospecific crosses never produce any viable offspring, and that females engaged in such matings cannot recover this fitness loss (see Figure 1). Only crosses between conspecifics produce viable offspring (see Figure 1). Knowing that a female with preference *p_f_* has mated with a conspecific male, the probability that this male displays the trait *t_m_* is given by:

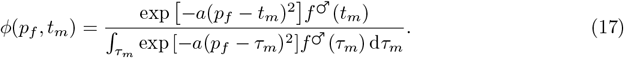

**Figure 1:**
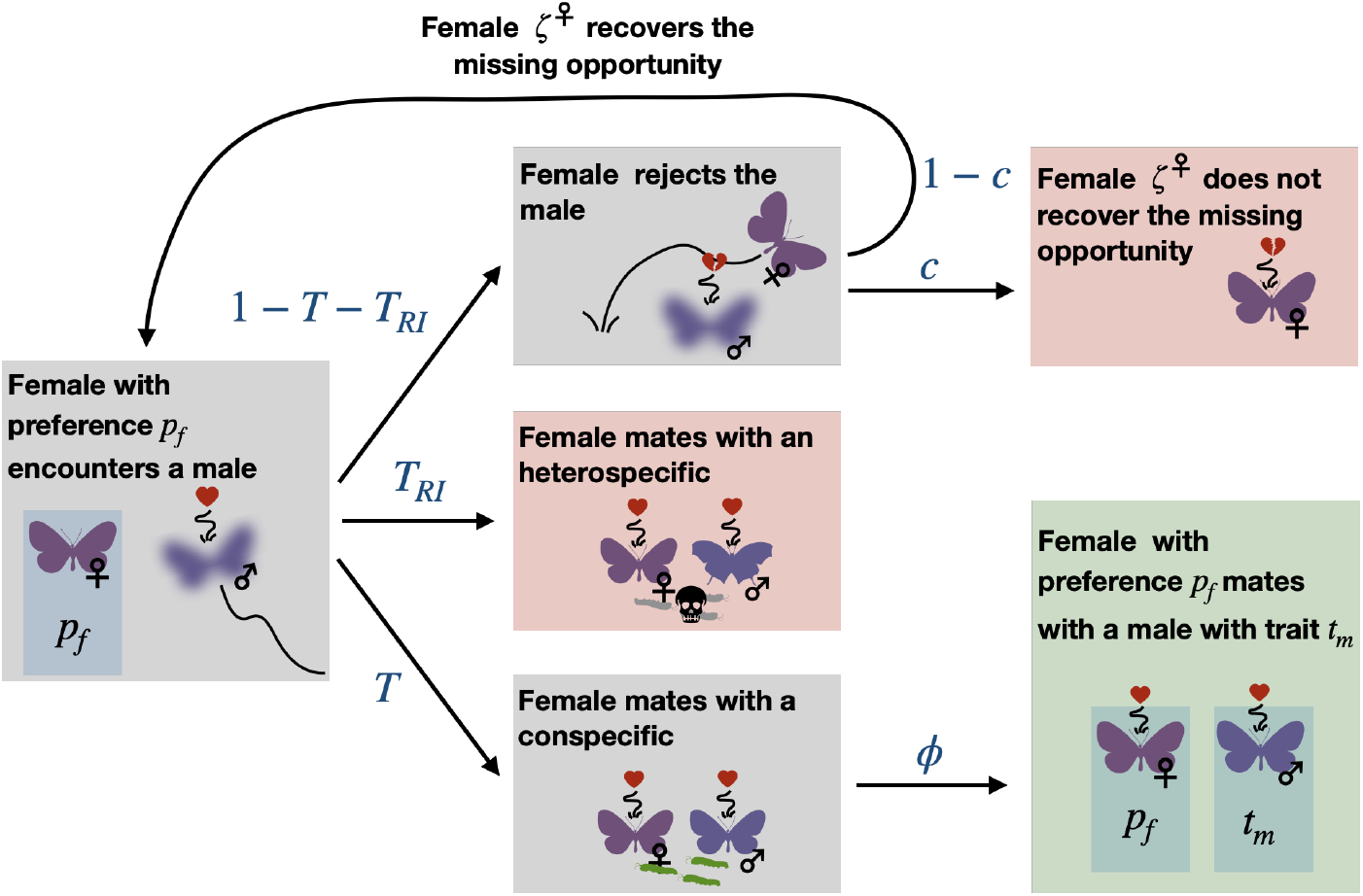
Computation of the contribution to the next generation of a mating. During an encounter, a female expresses her preference towards the warning trait displayed by the male and other cues that may differ between conspecific and heterospecific males. A female accepts a conspecific (resp. heterospecific) male with probability *T*(*p_f_*) (resp. *T_RI_*(*p_f_*)) (see Equation (13) (resp. (14))). A mating with an heterospecific male produces no viable offspring and the female cannot mate anymore. When the female mates with a conspecific of trait *t_m_*, the cross occurs with probability *ϕ*(*p_f_*, *t_m_*). During an encounter the female may refuse a mating opportunity with a male displaying a trait value *t_m_* distant from her preference *p_f_* and can subsequently encounter other males with probability 1 – *c*. Alternatively, she may not recover the fitness loss with probability *c*, resulting in an opportunity cost. The contribution to the next generation of a mating between a male with trait *t_m_* and a female with preference *p_f_* is thus given by *W_r_*(*t_m_*, *p_f_*) (see Equation (21)). Expressions in blue represent the probabilities associated with each arrow. In red, the female does not produce any offspring. In green, the mating between a male with trait *t_m_* and a female with preference *p_f_* happens and produces progeny.

Using again the assumption that the trait distribution has a low variance, this can be approximated by

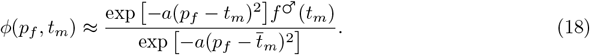

Considering that females only encounter one male, the proportion of crosses between a female with preference *p_f_* and a conspecific male with trait *t_m_* would be

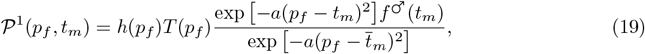

where *h* is the distribution of preferences in the population.

However, we assume that females refusing a mating opportunity can encounter another male with probability 1 – *c* (see Figure 1). We interpret *c* ∈ [0, 1] as the cost of choosiness (similar to the coefficient *c_r_* in [Otto et al., 2008]). The proportion of matings between a female with preference *p_f_* and a conspecific male with trait *t_m_* is thus given by

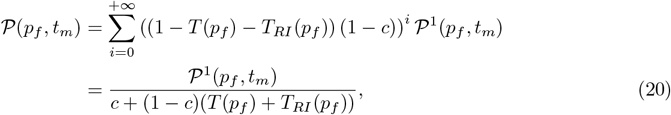

where ((1 – *T*(*p_f_*) – *T_RI_*(*p_f_*)) (1 – *c*))^*i*^ is the probability that a female with preference *p_f_* rejects the *i* males she first encounters and then encounters an (*i* + 1) – *th* male.

The contribution to the next generation of a mating between a male with trait *t_m_* and a female with preference *p_f_*, *W_r_* (*t_m_*, *p_f_*) is thus given by (see Figure 1)

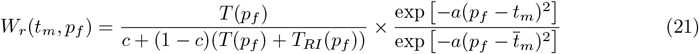

All variables and parameters used in the model are summed up in Table 1.

**Table 1:** Description of variables and parameters used in the model.

| Abbreviation | Description |
| --- | --- |
| $\bar{t}_m/\bar{t}_f$ | Mean trait value displayed in the *focal* species by males and females respectively |
| $\bar{p}_f$ | Mean female preference value in the *focal* species |
| $G$ | matrix of genetic covariance |
| $a$ | Females choosiness in the *focal* species |
| $s$ | Strength of developmental constraints in the *focal* species |
| $t_a$ | Ancestral trait favoured by developmental constraints in the *focal* species |
| $t'$ | Trait displayed in the *model* species |
| $d_m/d_f$ | Basal predation rate in males and females respectively |
| $b$ | Predators discrimination |
| $\lambda/\lambda'$ | Defence level of individuals of the *focal* and *model* species respectively |
| $N/N'$ | Density of the *focal* and *model* species respectively |
| $c_{RI}$ | Strength of reproductive interference |
| $c$ | Cost of choosiness |

### Relaxing the weak preference hypothesis

Because the stringency of females choice (*a*) is a key driver of the effect of reproductive interference on the convergence towards the trait displayed in the *model* species, we do not assume that *a* is always of order *ε*. Assuming such a strong sexual selection violates the weak selection hypothesis. However, because strong females choosiness leads to higher sexual selection, the discrepancy between females preference and males trait values 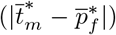 becomes limited. Therefore sexual selection and opportunity cost are actually weak and we can still estimate the matrix of genetic covariance and assume that the genetic variances of traits and preference are low.

### Model exploration

We assume that the *focal* species is ancestrally not in contact with the *model* species, and therefore the initial mean trait values displayed by males and females are equal to the optimal trait *t_a_*. We also assume that the mean female preference value is initially equal to the mean trait value displayed by males. At the initial time, we assume that the *focal* species enters in contact with the *model* species. The dynamics of traits and preference values then follow Equation (2). In Appendix 4 we explore two alternative scenarios: where the *focal* and the *model* species (1) ancestrally share common predators promoting mimicry before entering sexually in contact or (2) ancestrally interact sexually before sharing a common predator promoting mimicry.

#### Numerical simulations of the quantitative model

We use numerical simulations to estimate the traits and preference values at equilibrium 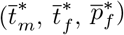. Numerically, we consider that the traits and preference are at equilibrium when

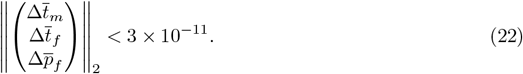

#### Individual-centred simulations

We also run individual-centred simulations with explicit genetic architecture to study the evolution of FLM with strong selection, as well as with high and fluctuating genetic variance of traits and preference. We assume two genetic architectures in an haploid population:

- Independent genetic basis of male and female trait: we assume three loci *T_m_*, *T_f_* and *P_f_* coding respectively for male trait, female trait and preference. We assume recombination rate between each loci *r_T_m_T_f__* and *r_T_f_P_f__*.
- Partially common genetic basis of male and female trait: we assume four loci *T*_1_, *T*_2_, *T*_3_ and *P_f_*. Locus *T*_2_ controls the trait variations shared by males and females and loci *T*_1_ and *T*_2_ (resp. *T*_2_ and *T*_3_) codes for specific male (resp. female) trait value with additive effect. *P_f_* codes for female preference value. We assume recombination rate between each loci *r*_*T*_1_*T*_2__, *r*_*T*_2_*T*_3__ and *r*_*T*_3_*P_f_*_.

We assume a constant standard deviation mutation effect across all loci *μ* and initial genetic variance of trait and preference *G*_0_ without genetic covariance. We also assume that population size stay constant. We run individual-centred simulations across 10,000 generations. Final traits and preference value are given by the mean value across the 1,000 last generations.

Scripts are available online at github.com/Ludovic-Maisonneuve/evo-flm.

#### Comparing alternative mechanisms inducing female-limited mimicry

First, we compare the evolutionary outcomes when assuming two alternative mechanisms generating FLM in an harmless species (λ = 0): (1) sexual selection generated by reproductive interference (*c_RI_* and *a* > 0) and (2) sexually contrasted predation (*d_f_* > *d_m_*). We thus compute the equilibrium traits and preference 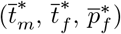 for different strengths of reproductive interference (*c_RI_* ∈ [0, 0.1]) or different basal predation rate sexual ratios between males and females *d_m_*/*d_f_* ∈ [0, 1]. Note that the two mechanisms are not mutually exclusive in natural populations. However here we investigate them separately to identify the specific evolutionary trajectories they generate. We then determine the range of key parameters enabling the evolution of FLM, under each mechanism assumed. We specifically follow the evolution of sexual dimorphism generated by each mechanism by comparing the level of sexual dimorphism at equilibrium defined by 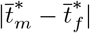.

#### Differential divergence from ancestral traits in male and female causing sexual dimorphism

To investigate whether the evolution of sexual dimorphism stems from increased divergence of traits from the ancestral states of one of the two sexes, we then compute the sexual bias in phenotypic divergence defined by

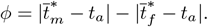

When *ϕ* < 0 we have 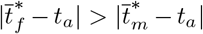 thus the trait diverged more in females than in males (see an illustration in Figure 2(a) and Figure 2(b)). By contrast *ϕ* > 0 indicates that the trait diverged more in males than in females (see an illustration in Figure 2(c)). We compare this sexual bias in phenotypic divergence under the two hypothetical mechanisms of FLM, to determine whether this criterium could be used to infer the actual evolutionary pressures involved in the emergence of FLM in natural populations.

**Figure 2:**
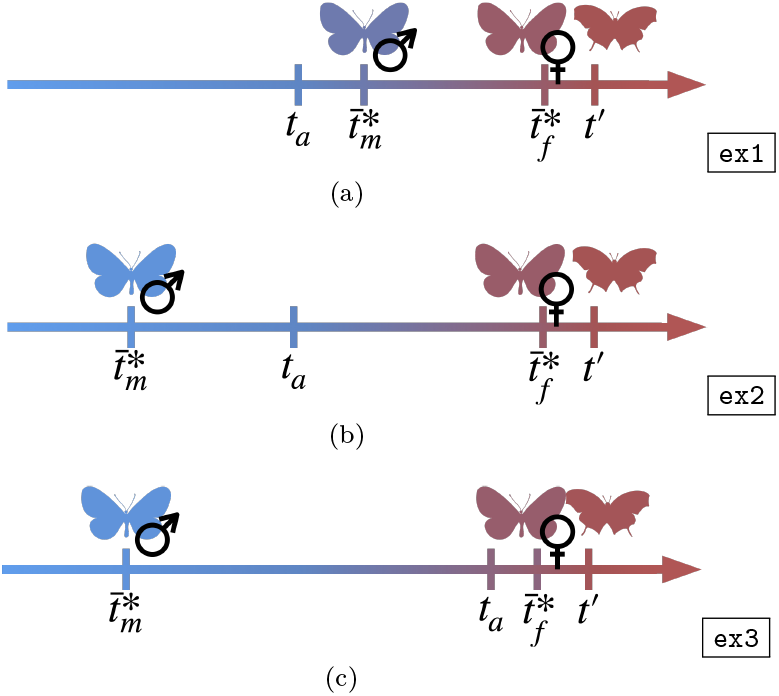
Illustration of the three main outcomes: (a) males trait value in the *focal* species gets closer to the value displayed in the *model* species *t*′, (b) males trait value in the *focal* species diverges away from the value displayed in the *model* species *t*′, (c) when the ancestral and the mimetic trait are close and males trait value in the *focal* species diverges away from the value displayed in the *model* species *t*′ then the phenotypic distance with the ancestral trait is higher in males than in females.

We first study the values of sexual bias in phenotypic divergence when reproductive interference causes FLM (*c_RI_* = 0.01), using numerical simulations. We investigate the effect of two key parameters: female choosiness *a* modulating cost of reproductive interference and the phenotypic distance between the ancestral trait *t_a_* and the mimetic trait *t*′. To investigate the impact of the phenotypic distance between the ancestral and the mimetic traits, we fixed the mimetic trait value to 1 (*t*′ = 1) and vary the ancestral trait value (*t_a_* ∈ [0, 1]) (see illustration in Figures 2(b) and 2(c)). We then study the sexual bias in phenotypic divergence when FLM stems from sexually contrasted predation (*d_f_* > *d_m_*), by deriving analytical results standing for all parameters value (see Appendix 5).

#### Investigating the impact of the defence level on the evolution of female-limited mimicry

Because FLM is usually reported for Batesian mimics, we then investigate the impact of the defence level (λ ∈ [0, 0.1]) on equilibrium traits 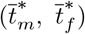 and the level of sexual dimorphism 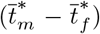. Because males and females in the *focal* species can display different traits, the level of protection gained by individuals of one sex through mimicry depends on males and females resemblance to the *model* species but also on the density of individuals of that sex within the *focal* species, modulated by the individual level of defence in the *focal* species (*λ*). When males from the *focal* species are non-mimetic, their defence level is given by the individual level of defence *λ* and the density of males *N*/2. To investigate the impact of defence level on the emergence of FLM, we thus explore not only the effect of the individual defence level *λ* but also of the density of the *focal* species (*N* ∈ [0, 20]).

The effects of all explored parameters and evolutionary forces on the evolution of FLM are summed up in Figure 3.

**Figure 3:**
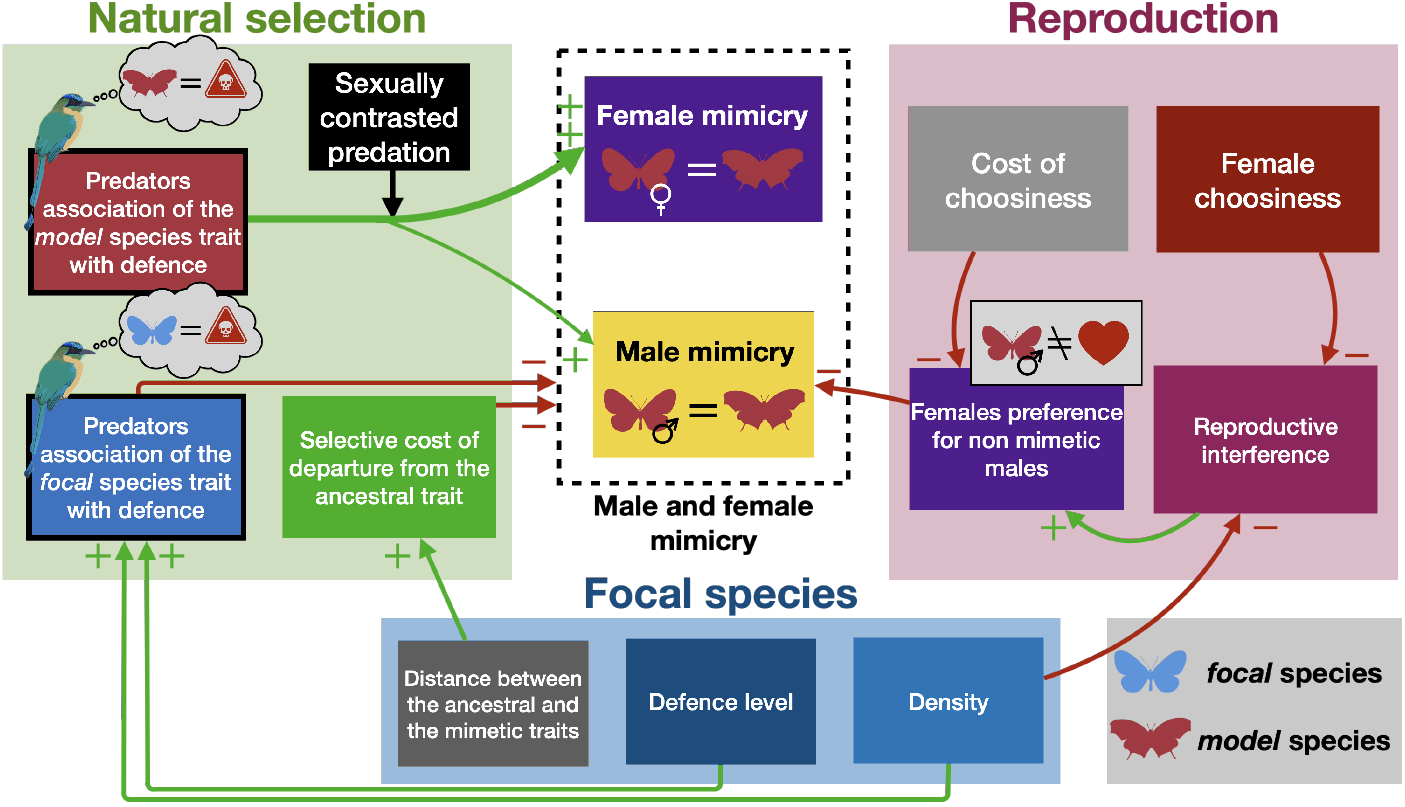
Summary of the impact of selective forces and parameters on the evolution of femalelimited mimicry. Green and red arrows represent the positive and negative impact respectively.

## Results

### Reproductive interference promotes female-limited mimicry in palatable species

We first test whether reproductive interference can generate FLM in a harmless species (λ = 0). We thus investigate the impact of the strength of reproductive interference (*c_RI_*) on the evolution of males trait 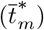, females trait and preference (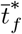 and 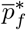), for different levels of females choosiness (*a*) modulating the costs generated by the strength of reproductive interference (Figure 4(a)). Without reproductive interference (*c_RI_* = 0), both males and females in the *focal* species are mimetic at equilibrium and the sexual dimorphism therefore does not emerge (Figure 4(a)). By contrast, when assuming reproductive interference (*c_RI_* > 0), FLM evolves in the *focal* species (Figure 4(a), see temporal dynamics in Figure A5(a)). Reproductive interference promotes a greater distance between final females preference 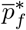 and the trait of the *model* species *t*′. Such females preference for non-mimetic males reduces costly sexual interactions with heterospecific males of the *model* species and generates sexual selection on males trait, inhibiting mimicry in males. Reproductive interference also promotes FLM in alternative scenarios when the *focal* and the *model* species (1) ancestrally share common predators promoting mimicry before entering sexually in contact or (2) ancestrally interact sexually before sharing a common predator promoting mimicry (see Appendix 4). Because FLM strongly depends on the evolution of females preference for potentially scarce non-mimetic males, it emerges only when the cost of choosiness (*c*) is low (see Appendix 7 for more details). FLM also evolves only when male and female traits have at least partially different genetic basis, allowing divergent evolution between sexes. The genetic covariance between males and females trait *G_t_m_t_f__* then only impacts the time to reach the equilibrium (see Appendix 8 for more details).

**Figure 4:**
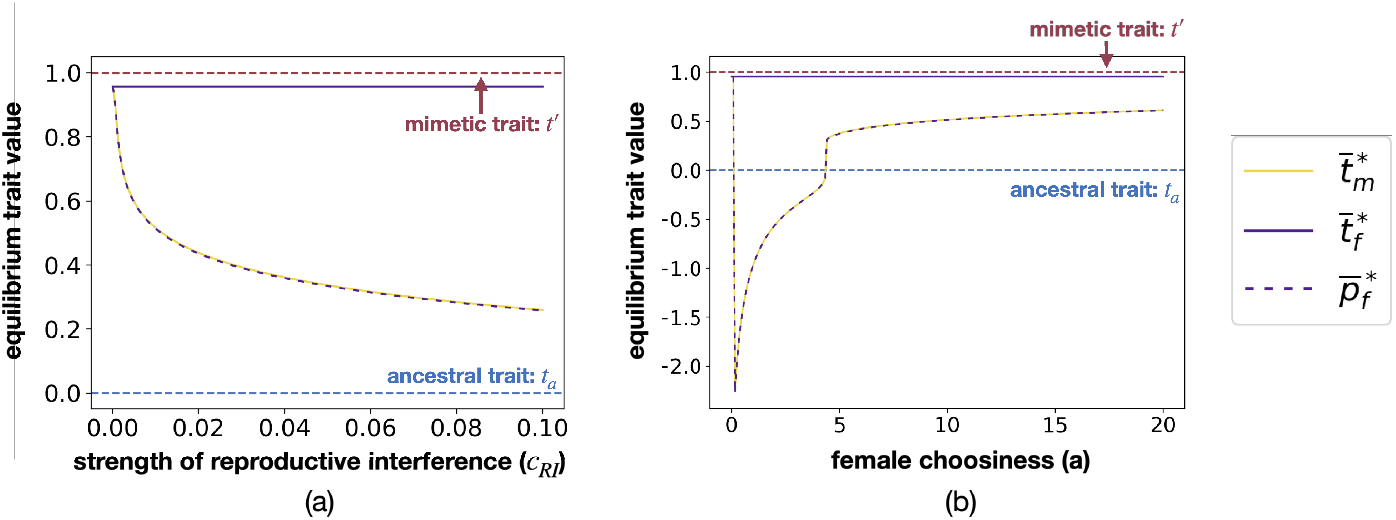
Influence of (a) the strength of reproductive interference *c_RI_* and (b) females choosiness *a* on the equilibrium values of males trait 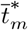 (yellow solid line), females trait 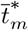 (purple solid line) and females preference 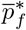 (purple dashed line). By default we assume: *G_t_m__* = *G_t_f__* = *G_p_f__* = 0.01, *G_t_m_t_f__* = 0.001, *c_RI_* = 0.01, *c* = 0.1, *a* = 10, *b* = 5, *d_m_* = *d_f_* = 0.05, λ = 0, *N* = 100, *λ*′ = 0.01, *N*′ = 200, *s* = 0.0025, *t_a_* = 0, 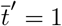.

We also investigate the impact of females choosiness (*a*) (modulating the stringency of sexual selection and cost of reproductive interference) on FLM, when there is reproductive interference (*c_RI_* > 0) (Figure 4(b)). The relationship between the final male trait value and the parameter *a* is sometimes discontinuous because for close value of parameters, the evolutionary dynamics can take different paths. When *a* is close to 0, both males and females become mimetic to the *model* species (Figure 4(b)). In this case, non-choosy females tend to accept almost all males, despite their preference *p_f_*. Thus selection on females preference *p_f_* is low because a change on preference hardly changes the mating behaviour and the resulting cost of reproductive interference. When *a* is higher than 0 and approximately lower than 5, selection due to reproductive interference on preference is important and reproductive interference promotes FLM. Furthermore, our results show that sexual selection does not only inhibit mimicry in males but may further promote divergence away from the ancestral trait *t_a_* (Figure 4(b), see Figure 2(b) for an illustration and Figure A5(b) for temporal dynamics). Such divergence from the ancestral trait in males does not occur when females choosiness is higher (*a* ≳ 5 in Figure 4(b) see Figure 2(a) for an illustration): when females are more picky, a small difference between female preference and the mimetic trait sufficiently reduces the cost of reproductive interference (Figure 4(b)). All results described in this section are confirmed in individual-centred simulations assuming simple genetic architecture of traits and preference (Figures A10 and A11), highlighting that the weak selection, constant and low genetic variance hypotheses does not preclude obtaining relevant analytical predictions.

### Sexually contrasted predation promotes female-limited mimicry in palatable species

Higher predation pressure acting on females has been proposed to explain FLM. Here we investigate the impact of the ratio of basal predation rate on males and females (*d_m_/d_f_*) on the evolution on FLM (Figure 5(a)) in case without reproductive interference and preference (*c_RI_* = 0, *a* = 0). When predation pressures are largely lower in males than in females (*i.e. d_m_/d_f_* ≲ 0.2), sexually contrasted predation promotes FLM (Figure 5(a), and see temporal dynamics in Figure A5(c)). Limited predation pressure in males implies low advantage to mimicry that is overcome by developmental constraints. By contrast, predation pressure is higher on females, resulting in a greater advantage to mimicry that overcomes costs of departure from ancestral trait value. However, when the predation ratio increases (*i.e. d_m_/d_f_* ≲ 0.2), sexual dimorphism is low, because advantage to mimicry in males becomes greater as compared to costs generated by developmental constraints (Figure 5(a)). When males and females suffer from similar predation pressure (*i.e. d_m_/d_f_* = 1), both sexes become mimetic (Figure 5(a)).

**Figure 5:**
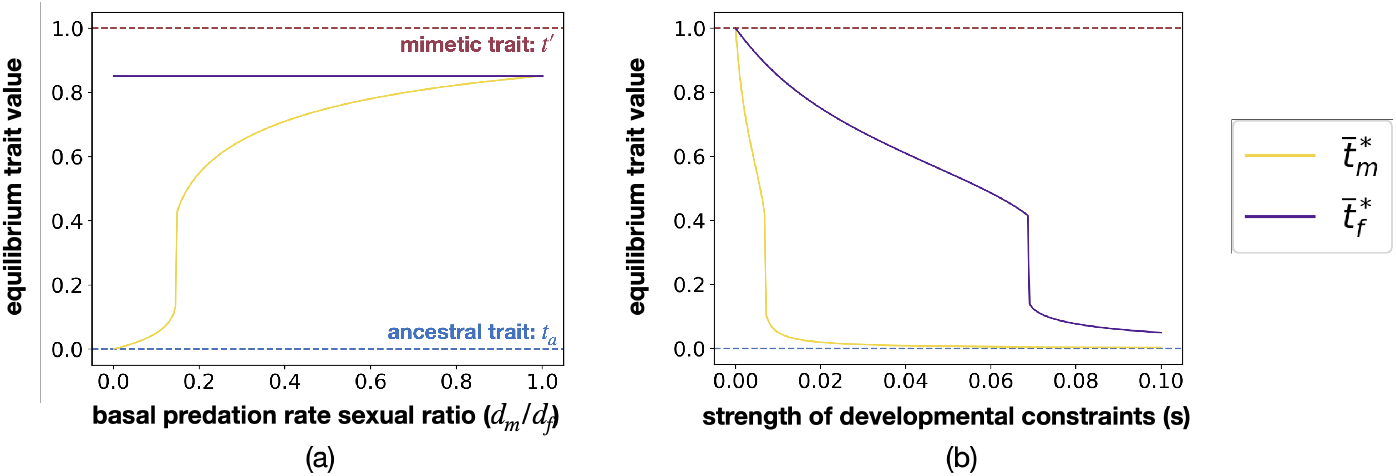
Influence of (a) the ratio of basal predation rate on males and females *d_m_/d_f_* and (b) the strength of developmental constraints *s* on the equilibrium values of males trait 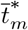 (yellow solid line), and females trait 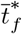 (purple solid line). By default we assume: *G_t_m__* = *G_t_f__* = *G_p_f__* = 0.01, *G_t_m_t_f__* = 0.001, *c_RI_* = 0, *c* = 0, *a* = 0, *b* = 5, *d_m_* = 0.005, *d_f_* = 0.05, λ = 0, *N* = 100, λ′ = 0.01, *N*′ = 200, *s* = 0.01, *t_a_* = 0, 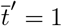.

Because developmental constraints are a major factor limiting mimicry, we then investigate the impact of the strength of developmental constraints (*s*) on FLM generated by a sexually contrasted predation (*d_m_/d_f_* = 0.1). When there is no developmental constraints (*s* = 0), FLM does not evolve, because males become mimetic even if they suffer for low predation. However, higher developmental constraints (0.1 ≲ *s* ≲ 0.7) limit mimicry in males, but not in females because of sexually contrasted predation (see previous paragraph). Important developmental constraints (*s* ≳ 0.7) overcome the advantages provided by mimicry in both sexes, and prevent the evolution of sexual dimorphism. Similarly to the previous section, all results shown in this section still hold in our individual-centred simulations (Figures A12 and A13)

### Different hypothetical causes of female-limited mimicry lead to different predictions

Here, we use our mathematical model to compare the effect of (1) reproductive interference and (2) sexually contrasted predation on the evolution of FLM. We specifically investigate in which sex the trait evolves away from the ancestral trait, depending on the selective mechanism causing FLM.

First, we focus on the evolution of FLM caused by reproductive interference via sexual selection (*a* > 0 and *d_f_* = *d_m_*). We specifically estimate how (1) the distance between the ancestral trait and the mimetic trait |*t_a_* – *t*′| and (2) the female choosiness *a* modulate sexual selection and shape the relative divergence of males and females from the ancestral trait value 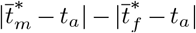. Figure 6 highlights that divergence from the ancestral trait can be stronger in males (yellow zone on figure 6(c)) or in females (purple zone on Figure 6(c)) depending on these parameters.

**Figure 6:**
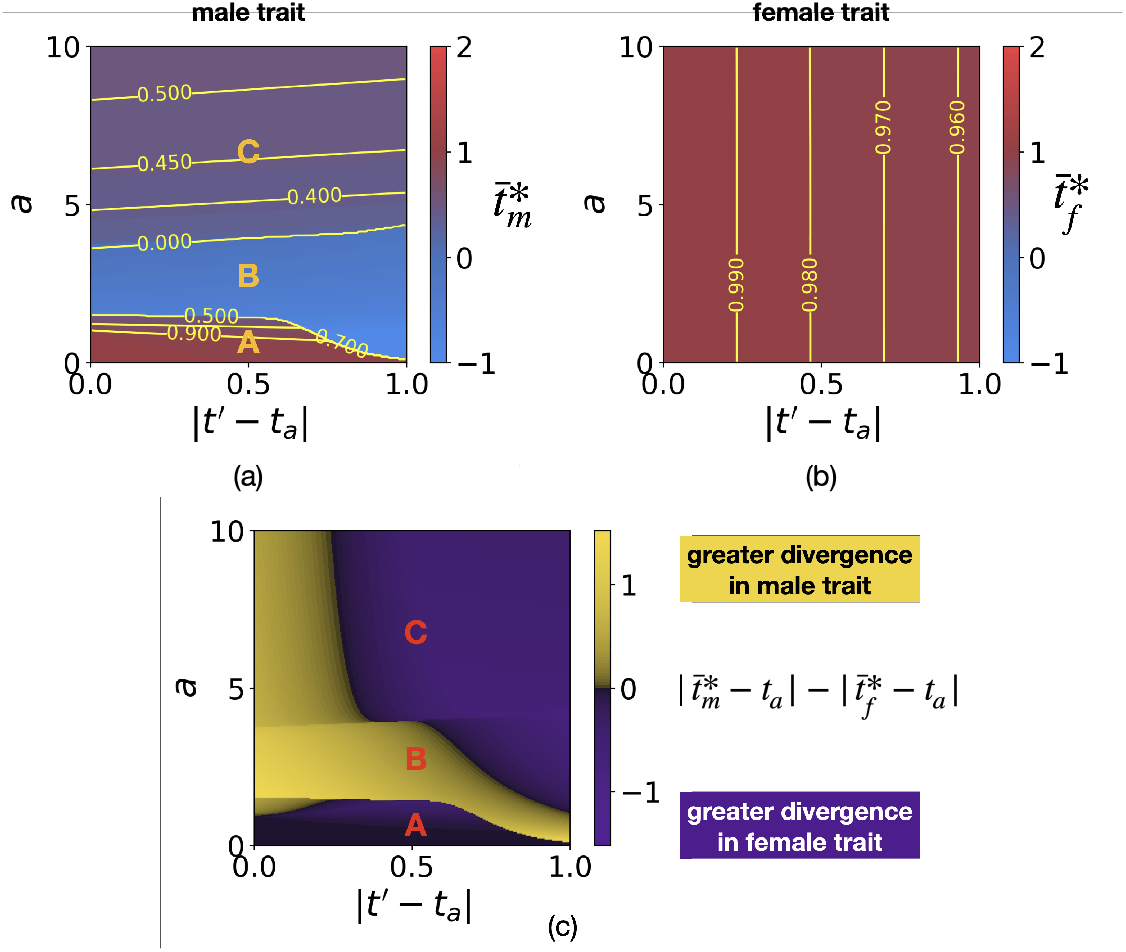
Influence of the distance between the ancestral and the mimetic traits |*t*′ – *t_a_*| and of females choosiness *a* on (a) final male trait 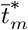, (b) final female trait 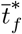 and (c) the difference between the level of divergence in males and females 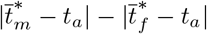. Note that Figure 6(c) results from Figures 6(a) and 6(b). Yellow lines indicate equal levels of trait value. We assume: *G_t_m__* = *G_t_f__* = *G_p_f__* = 0.01, *G_t_m_t_f__* = 0.001, *c_RI_* = 0.01, *c* = 0.1, *b* = 5, *d_m_* = *d_f_* = 0.05, λ = 0, *N* = 100, λ′ = 0.01, *N*′ = 200, *s* = 0.0025, *t*′ = 1.

The evolution of female trait only depends on the distance between the ancestral trait *t_a_* and the mimetic trait *t*′: because selection always promotes mimicry in females, divergence from the ancestral trait increases with the initial distance from the mimetic trait (Figure 6(b)). The level of mimicry in females slightly decreases with the ancestral level of mimicry because it increases the costs of developmental constraints. However, such costs are still overcame by the advantage of being mimetic. By contrast, the evolution of male trait depends on the interplay between the sexual selection generated by female preferences and the ancestral level of mimicry (Figure 6(a)).

The relationship between the final male trait and the parameters is discontinuous as previously highlighted, leading to three zones within where male trait vary continuously. When female choosiness is low (zone A, *a* ≲ 1.8), the selection caused by reproductive interference is mild: females are not very choosy and thus tend to accept almost all males despite their preference *p_f_*, therefore relaxing selection on females preference, and favouring the evolution of mimetic trait in males. Mimicry is nevertheless more accurate in females than in males, and males phenotype tends to stay closer to the ancestral trait value, and to display a so-called ”imperfect” mimicry. When the ancestral level of mimicry is poor (|*t_a_* – *t*′ | ~ 1), the slight advantage in sexual selection can then overcome the advantage of imperfect mimicry, resulting to divergence in males trait, even for low values of females choosiness (*a* ≲ 1.8).

However, when females choosiness has intermediate values (1.8 ≲ *a* ≲ 4, zone B), enhanced female choosiness increases selection due to reproductive interference and thus reduces mimicry in males. Nevertheless, when the distance between the ancestral and the mimetic trait is already large, divergence in male trait is limited, and the sexual dimorphism mainly stems from the evolution of mimicry in females. Using individual-centred simulations, we then show that stochastic variations may result in the divergence of male trait away from the ancestral trait, when the initial distance between the ancestral trait and the mimetic trait is low (|*t_a_* – *t*′| ≃ 0), (see Figure A19).

Contrastingly, high levels of choosiness in females (*a* ≳ 4, zone C) promote the evolution of more mimetic males because even a slight difference between the females preference and the mimetic trait allows to reduce cost of reproductive interference. Male divergence is then observed only when the ancestral level of resemblance between the *focal* and the *model* species is very high (*i.e* low |*t_a_* – *t*′|), and therefore induced cost of reproductive interference, despite the high pickiness (*i.e*. high *a*) of females.

The evolution of FLM caused by reproductive interference therefore leads to different divergence patterns, including divergence of male phenotypes away from the ancestral trait value. In contrast when FLM is caused by sexually contrasted predation (*d_f_* > *d_m_* and *a* = 0), sexual dimorphism always stems from the evolution of female phenotypes away from the ancestral trait, *i.e*. 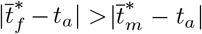 (see Appendix 5 and see Figure 2(a) for an illustration). Individual-centred simulations confirm this pattern, except when the distance between the ancestral trait and the mimetic trait is low (|*t_a_* – *t*′| ≃ 0). In this case, developmental constraints and predation promote the same trait value (*t_a_* ≃ *t*′). Higher stabilising selection in females due to higher predation pressure implies than females trait diverge less from the ancestral trait than males.

While both the reproductive interference and the sexually-contrasted predation may result in FLM, the evolutionary pathways causing the sexual dimorphism are strikingly different. These results are generally maintained when relaxing the weak selection, constant and low genetic variance hypotheses (see Appendix 11)

### The evolution of FLM depends on defence level

We then investigate the impact of the individual defence level (λ) and the density (*N*) in the *focal* species on the evolution of sexual dimorphism, when FLM is generated either (1) by sexually contrasted predation (Figure 7) or (2) by reproductive interference via sexual selection (Figure 8).

**Figure 7:**
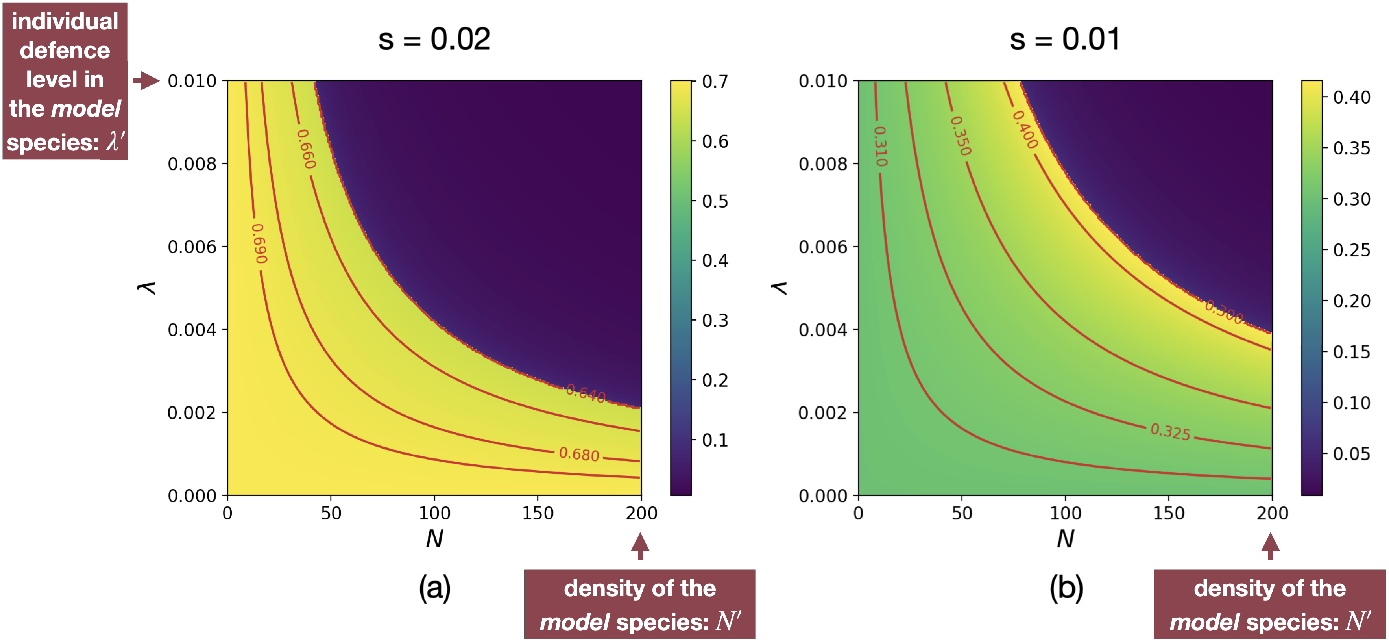
Influence of the density *N* and of the individual defence level λ in the *focal* species on the equilibrium values of the level of sexual dimorphism 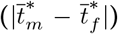 for different strength of developmental constraints ((a) *s* = 0.02 (b) *s* = 0.01) when femalelimited mimicry is caused by sexually contrasted predation (*d_f_* > *d_m_*, *a* = 0). Red lines indicate equal levels of sexual dimorphism. We assume: *G_t_m__* = *G_t_f__* = *G_p_f__* = 0.01, *G_t_m_t_f__* = 0.001, *c_RI_* = 0, *c* = 0, *a* = 0, *b* = 5, *d_m_* = 0.01, *d_f_* = 0.05, λ′ = 0.01, *N*′ = 200, *t_a_* = 0, *t*′ = 1.

**Figure 8:**
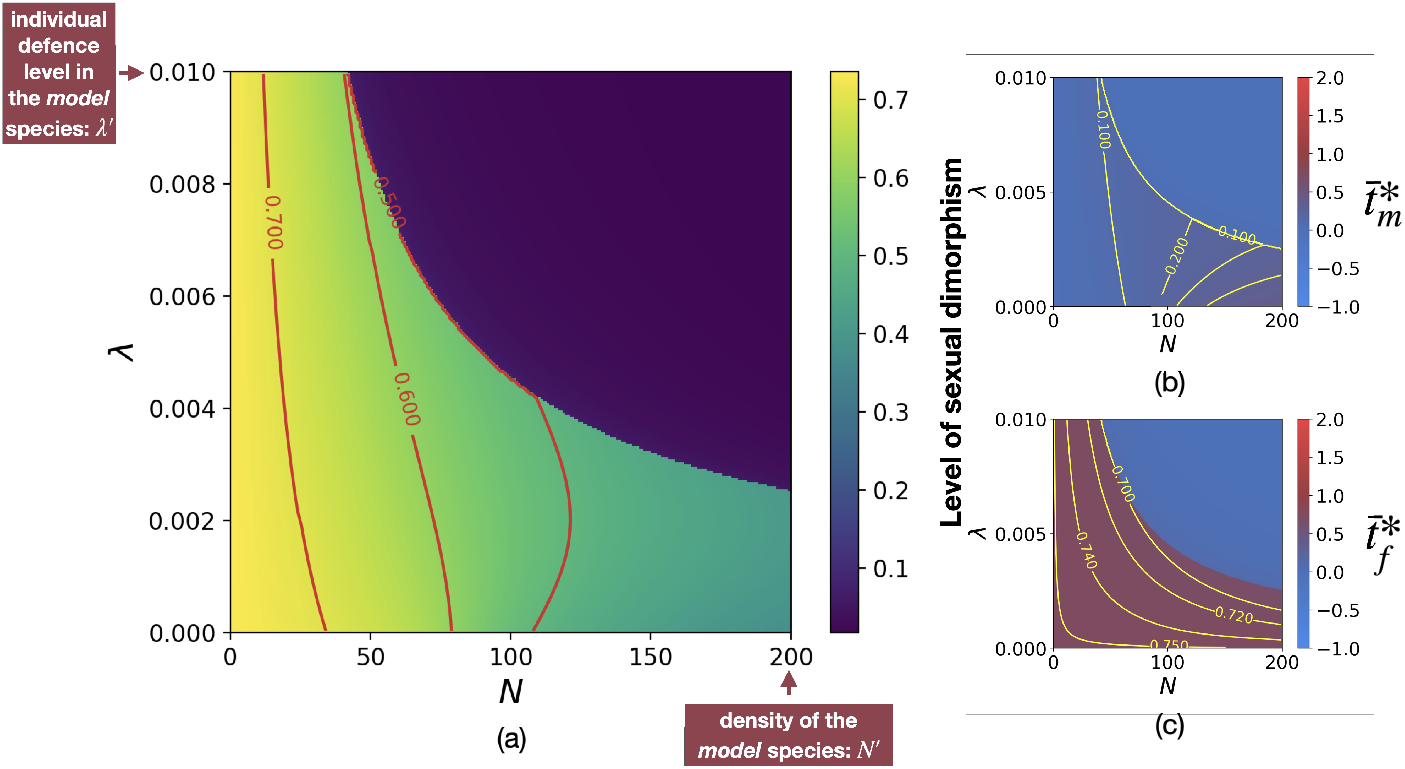
Influence of the density *N* and of the individual defence level λ in the *focal* species on the equilibrium values of (a) the level of sexual dimorphism 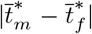, (b) males trait 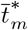 and (c) females trait 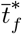 when female-limited mimicry is generated by sexual selection caused by reproductive interference (*c_RI_*, *a* > 0 and *d_f_* = *d_m_*). Red and yellow lines indicate equal levels of sexual dimorphism and trait value respectively. We assume: *G_t_m__* = *G_t_f__* = *G_p_f__* = 0.01, *G_t_m_t_f__* = 0.001, *c_RI_* = 0.01, *c* = 0.1, *a* = 5, *b* = 5, *d_m_* = *d_f_* = 0.05, λ′ = 0.01, *N*′ = 200, *s* = 0.02, *t_a_* = 0, *t*′ = 1.

Surprisingly, when FLM is caused by sexually-contrasted predation (*d_f_* > *d_m_*), the level of sexual dimorphism can either increase or decrease with defence levels in both males and females (λ*N*/2), depending on the strength of developmental constraints (Figure 7). In both sexes, the increase in defence levels indeed reduces selection favouring mimicry, while the developmental and selective constraints favour ancestral trait value. Great strength of developmental constraints (*s* = 0.02) then totally limits mimicry in males for every defence levels (Figure A20(a)). An increase in defence levels reduces mimicry in females (Figure A20(b)) but not in males that always displays the ancestral trait resulting in a decrease of the level of sexual dimorphism (Figure 7(a)). By contrast, low strength of developmental constraints (*s* = 0.01) allow the evolution of imperfect mimicry in males. However, the evolution of such mimicry in males is strongly impaired when defence level increases. In this range of mild levels of defence, mimicry is nevertheless advantageous in heavily-attacked females (Figure A21(b)), resulting in high level of sexual dimorphism (Figure 7(a)). However, when the defence level becomes very high, both males and females display the ancestral trait, and sexual dimorphism is no longer observed (Figures A21 and A20 at the top right). Because of the high level of defence, individuals of both sexes gain sufficient protection from similarity with their conspecifics, relaxing selection promoting mimicry towards the *model* species. Individual-centred simulations provide the same patterns. Interestingly, the only discrepancy is observed for the effect of the density of the *focal* species when developmental constraints are low: in this case, the level sexual dimorphism no longer increases with with density of the *focal* species(see Appendix 13), contrary to what was observed in the deterministic model (A20(a)). Stochasticity of population mean males and females trait value that is likely to increase sexual dimorphism. The amplitude of this stochastic effect reduce with population density that decrease the level of sexual dimorphism because when traits evolves randomly it is likely to produce sexual dimorphism (see figure A25).

Similarly, when FLM is caused by reproductive interference (*c_RI_* > 0) *via* sexual selection, the level of sexual dimorphism can also either increase or decrease with the individual defence level λ depending on the strength of developmental constraints (Figures 8(a) and A22(a)). In contrast with predation differences between sexes, sexual selection induced by reproductive interference generates markedly higher sexual dimorphism for low values of density of the *focal* species 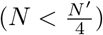 (Figure 8(a)). The relative density of the *focal* and the *model* species indeed determines the probability that a female of the *focal* species encounters a conspecific rather than an heterospecific male and thus modulates the costs of reproductive interference. Therefore, when the density of the *focal* species *N* is low, costs of reproductive interference are great, generating higher selection promoting sexual dimorphism. The density of the *focal* species therefore impacts much more the level of sexual dimorphism than the individual defence level λ.

Under both hypotheses explaining female limited-mimicry, when developmental constraints totally inhibit mimicry in males, sexual dimorphism decrease with the level of defence. Under the assumption of sexual selection generated by reproductive interference however, sexual dimorphism is higher when the *focal* species is rarer than the *model* species.

Under both selective hypotheses, mimicry toward the sympatric defended *model* species is no longer promoted in either sexes, when the level of defence within the *focal* species is high (Figures A21, A20 and 8(b)(c)) leading to sexual monomorphism. The distance between the ancestral and the mimetic traits |*t*′ – *t_a_* | limits mimicry in both sexes (Figure A23) highlighting the important role of the initial advantage and disadvantage of mimicry. Using individual-centred simulations, we nevertheless observed that males and females trait can get closer to the mimetic trait by stochasticity, enabling mimicry to be promoted, when the level of defence within the *focal* species is high (Figures A24, A26 and A28).

## Discussion

### Ancestral levels of resemblance, sexually-contrasted divergences and the evolution of female-limited mimicry

Our model highlights that both (1) sexually contrasted predation and (2) females preference generated by reproductive interference can favour the evolution of FLM. By explicitly studying how these contrasted selective pressures influence the divergence of males and females traits from a common ancestral trait, our model sheds light on contrasted evolutionary pathways towards sexual dimorphism. Empirical studies based on the estimation of the level of divergence in males and females traits usually interpret elevated divergence in males trait as compared to female trait, as a signature of sexual selection, causing sexual dimorphism [van der Bijl et al., 2020]. Focusing on FLM in *Papilio* butterflies, Kunte [2008] shows that sexual dimorphism is correlated with divergence in females trait, and concluded that FLM is caused by natural selection. However, our results show that when reproductive interference induces females preference, FLM can also stem from an increased divergence in female trait. Our results therefore highlight that higher divergence in female trait is not a reliable evidence of sexually-contrasted selection promoting FLM.

Contrary to reproductive interference, sexually-contrasted predation can generate FLM only when the *focal* and the *model* species have different ancestral traits. Such mechanism would thus be especially relevant for distantly-related co-mimetic species, that are more likely to have divergent ancestors. In contrast, the role of reproductive interference in generating FLM is probably more important in cases where mimetic and model species are more closely related. Our results also show that a non-mimetic ancestral state favour the emergence of FLM under sexually-contrasted selection. Therefore, the FLM observed in *Papilio garamas*, which likely derived from a sexually monomorphic and mimetic ancestor [Kunte, 2009], might be a good candidate to investigate the potential origin of FLM due to reproductive interference. Our results thus stress the need to infer the for ancestral levels of mimicry,as well as the phylogenetic distances between mimetic species and their co-mimics or model species to empirically investigate the effect of reproductive interference on the evolution of FLM.

### The level of investment of males in reproduction and the evolution of FLM caused by reproductive interference

Our results show that reproductive interference can generate females preference for non-mimetic males and therefore may cause FLM. Some studies already suggested that sexual selection may generate FLM [Belt, 1874., Turner, 1978], but the origin of females preferences for non-mimetic males was unidentified. Our model highlights that reproductive interference could be the driver of such females preferences.

Nevertheless, the emergence of sexual dimorphism stems from the assumption that female is the only choosy sex. This assumption is relevant when females invest much more in reproduction than males [Trivers, 1972, Balshine et al., 2002]. However, this asymmetrical investment in offspring between males and females can vary in different Lepidoptera species. In some species, butterfly males provide a nuptial gift containing nutriments during mating [Boggs and Gilbert, 1979]. Such elevated cost of mating in males could promote the evolution of choosiness in males. If the asymmetry in reproductive investment between sexes is limited, the evolution of FLM would then be impaired. Moreover, the investment of males in reproduction impacts the cost of choosiness for females, because females refusing a mating opportunity would be denied access to the nuptial gift. In Lepidoptera, females mating more that once have higher lifetime fecundity than females that mate only once, because nuptial gifts provide important metabolic resources [Wiklund et al., 1993, Lamunyon, 1997]. Such elevated cost of rejecting a potential mate may limit the evolution of preference in females, as highlighted by our model: our results indeed show that reproductive interference promotes FLM only when cost of choosiness is low. The evolution of female-mimicry is thus likely to be impaired when the costs of mating are elevated in males, and therefore (1) inducing male choosiness and (2) increasing the opportunity costs generated by female choosiness.

Even when females are the choosy sex, they can still have preference based on multiple cues reducing cost of reproductive interference. Butterflies express preference for pheromones that may strongly differ between closely related species [Darragh et al., 2017, González-Rojas et al., 2020] thus limiting cost of reproductive interference. Moreover, different micro-habitat preference may reduces interspecific interactions and then female probability of accepting a heterospecific male [Estrada and Jiggins, 2002]. In our model, the probability to reject an heterospecific male based on other trait than the warning trait is captured by the parameters *c_RI_*. Our results show that reproductive interference can promote FLM even when *c_RI_* is low. As soon as *c_RI_* is non-null, reproductive interference lead to selection on females preference and the evolution of FLM depends on the relative importance of each evolutionary forces.

Because few studies investigate the sexual selection origin of FLM, empirical studies estimating the reproductive costs and benefits in both sexes are strongly lacking. Here, we explicit a mechanism by which sexual selection can generate FLM. We thus hope our theoretical work will encourage experimental approaches investigating the link between reproductive costs and FLM. Such studies may shed light on the actual role of sexual selection generated by RI on the evolution of FLM.

### Relative species abundances and defences and the evolution of female-limited mimicry

Our results show that, for both causes of FLM (reproductive interference or sexually contrasted predation), the level of sexual dimorphism decreases with the individual level of defence when developmental constraints totally inhibit mimicry in males. This prediction is consistent with the empirical observation reporting FLM mostly in Batesian mimics, although FLM has still been reported in a few defended species [Nishida, 2017]. Our model stresses the need to precisely quantify the level of defences carried out by individuals from different species: important variations in the levels of defences within species have been documented in Müllerian mimics (*e.g*. in *Heliconius* butterflies, Sculfort et al. [2020]), as well as in Batesian mimics (e.g. viceroy butterfly, Prudic et al. [2019]). Empirical quantification of the level of deterrence induced by individuals from co-mimetic species would shed light on the evolutionary conditions favouring the evolution of FLM.

Our model also predicts that the emergence of FLM is strongly linked to the relative density between mimics and models, and our theoretical approach neglects the dynamics of population densities of the *focal* and the *model* species, that may depend on their individual defence level. Empirical studies usually report that the density of undefended mimics is low compared to those of the defended models [Long et al., 2015, Prusa and Hill, 2021]. Undefended mimics can have a negative effect predator’s learning [Rowland et al., 2010, Lindström et al., 1997], suggesting that Batesian mimicry could evolve and be maintained only in species with a low density compared to the *model* species. Moreover, a high abundance of the *model* species compared to the potential mimics also increases the protection of imperfect mimics allowing the evolution of gradual Batesian mimicry [Kikuchi and Pfennig, 2010]. The relative density between the *focal* and the *model* species is especially important when assuming reproductive interference, because the costs generated by heterospecific interactions depend on the proportion of heterospecific males encountered by females. Our results show that reproductive interference strongly promotes sexual dimorphism when the density of the *focal* species in low as compared to the *model* species. Considering that FLM is caused by reproductive interference, the lower relative density of undefended species may promote FLM, and therefore explain why FLM could be especially favoured in Batesian mimics is reserved to undefended species.

The reported difference in phenology between defended *models* emerging sooner than undefended *mimics* may further enhance the difference in relative abundances between *models* and *mimics*, therefore increasing the cost of reproductive interference for undefended females. Batesian mimics often emerge after their models, when the models warning trait is well known by predators [Prusa and Hill, 2021], and this might reinforce the evolution of FLM caused by reproductive interference in Batesian *mimics*. Overall, our theoretical study stresses the need of ecology studies quantifying relative densities of mimetic defended and palatable species through time. Such field studies, as well as chemical ecology studies quantifying defence variations, are now crucial needed to understand the evolution of FLM, in Batesian and Müllerian mimics.

### Sexual conflict limiting males adaptation

Our study highlight that different fitness optima among sexes, due to natural and sexual selections, drives the evolution of sexual dimorphism in both hypothesis explaining FLM. Different fitness optima may stem from sexually dimorphic morphology, leading to different flight ability and to sexually contrasted predation risk. But different sexual roles, such as different levels of physiological investments in offspring, may also leads to contrasted effect of trait variations on female and male fitness, generating so-called sexual conflicts [Parker, 2006]. Sexual conflicts classically involves the evolution of traits enhancing male mating success with multiple females, and of traits enhancing the rejection of non-preferred males in females (*e.g*. conflicting coevolution of genitalia in males and females Brennan et al. [2010]. FLM driven by reproductive interference provide an original example of sexual conflict: while mimicry would enhance survival in males, female preferences generated by reproductive interference and by their greater reproductive investment, prevent the evolution of mimetic trait in males. This is thus a relevant case-study of sexual conflict driving the evolution of sexual dimorphism. Similarly, costly exaggerated trait in males may be regarded as a results of sexual conflicts: female prefer this expensive trait sign of mate quality (handicap principle [Zahavi, 1975]) leading to maladaptive trait disfavoured by natural selection [Johnstone, 1995]. In black scavenger flies *Sepsis cynipsea* and *Sepsis neocynipsea* species differentiation of exaggerated male forelegs is higher in sympatric population [Baur et al., 2020], suggesting than species interactions may indeed be a key evolutionary force involved in the evolution of exaggerated trait in males. Reproductive interference is indeed expected to promote male exaggerated trait improving species recognition in females. However, evidences of the role of reproductive interference in the evolution of sexual dimorphism are still scarce. Our theoretical work on FLM highlights that conflict between natural selection promoting the same trait in different species and reproductive interference may generate sexual dimorphism. We thus hope our results will stimulate new research on the effect of ecological interactions between closely-related species on the evolution of sexual dimorphism.

## Conclusion

Our model show that both sexually contrasted predation and reproductive interference (by promoting preference for non-mimetic males) may generate FLM. Our results therefore show that the patterns of divergence of males and females traits from ancestral state should be interpreted in light from the selection regime involved. Our model also reveals the important role of ecological interactions between sympatric species on the evolution of sexual dimorphism, highlighting the need to consider the role of reproductive interference in the phenotypic diversification in sympatry.

## Acknowledgments

The authors would like to thank the ANR SUPERGENE (ANR-18-CE02-0019) for funding the PhD of LM. This work was partially supported by the Chair “Modélisation Mathématique et Biodiversité” of VEOLIA-Ecole Polytechnique-MNHN-F.X.

## Appendix

### 1 Selection vectors

In this part we detail the calculations to obtain the selection vector (Equation (2)).

#### 1.1 Selection acting on males trait *β_t_m__*

We compute the first component of the selection vector *β_t_m__* describing the selection acting on males trait. This coefficient is given by

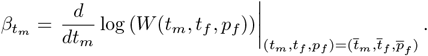

Using (1) and (6) we have

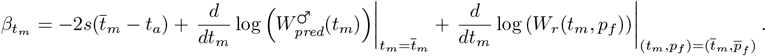

##### 1.1.1 Selection due to predation

First we compute the part of the selection coefficient due to predation. Using (10) we have:

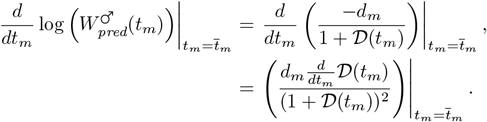

Using (9) we have

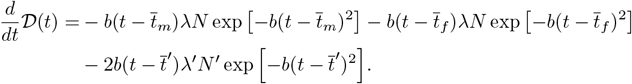

##### 1.1.2 Selection due to reproduction

We now compute the part of the selection coefficient due to reproduction. Using (21) we have:

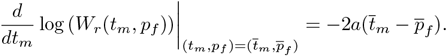

Therefore we have

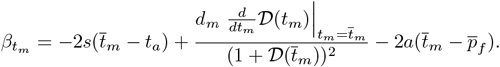

#### 1.2 Selection acting on females trait *β_t_f__*

The second component of the selection vector *β_t_f__* is given by

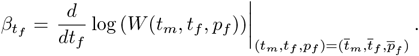

Using (1) and (7) we have

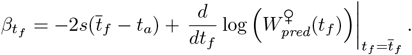

Similarly than with male traits we have

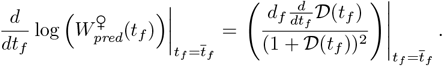

Thus we have

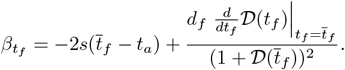

#### 1.3 Selection acting on females preference *β_p_f__*

The last component of the selection vector *β_t_f__* is given by

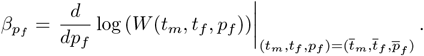

Using (1) we have

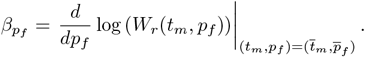

Using (21) we have

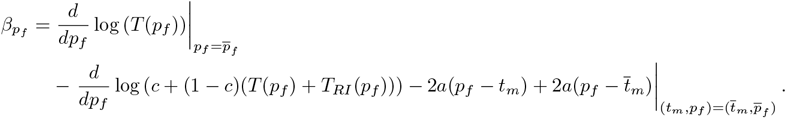

Using (15) and (16) we have

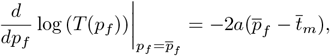

and

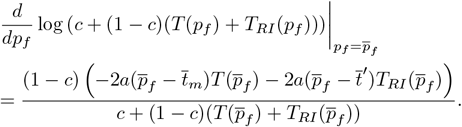

Thus

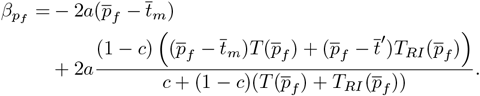

### 2 Computation of the matrix of correlation

In this part we approximate the genetic covariance between males trait and females preference *G_t_m_p_f__*, using the results from [Kirkpatrick et al., 2002]. Trait and preference are controled by different sets of unlinked loci with additive effects, denoted *T* and *P*, respectively. We note *T_m_* ⊆ *T* and *T_f_* ⊆ T the loci controlling trait in males and in females respectively. For each *i* in *T* (resp. *P*), we note 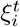 (resp.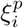) the contribution of the locus *i* on trait (resp. preference) value. The trait *t_m_* of a male is then given by

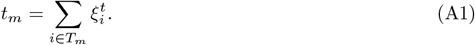

The trait *t_f_* and preference *p_f_* values of a female are given by

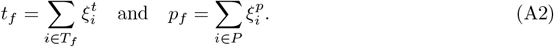

As in [Lande, 1981] we assume that the distributions of 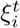 and 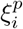 are multivariate Gaussian. Let *G_ij_* be the genetic covariance between loci *i* and *j*. Then the elements of the matrix of correlation are given by:

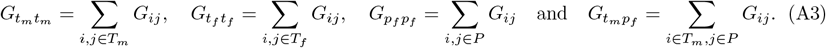

To compute the change on genetic correlation we need to identify various selection coefficients (see [Barton and Turelli, 1991, Kirkpatrick et al., 2002]). These coefficients are obtained using the contribution to the next generation of a mating between a male with trait *t_m_* and a female with trait *t_f_* and preference *p_f_* due to natural selection and mating preference (see equation 1).

For simplicity we consider only leading terms in the change in genetic correlation, computed with a Mathematica script (available online at https://github.com/Ludovic-Maisonneuve/evo-flm). For (*i, j*) ∈ *T_m_* × *P_f_*, combining Equations (9), (12), (15) from Kirkpatrick et al. [2002] gives the change in the genetic covariance between loci *i* and *j*:

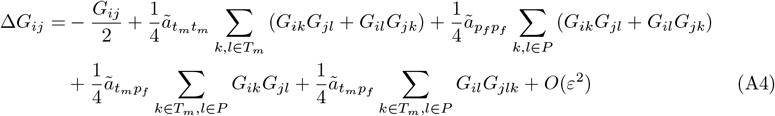

with 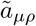 for (*μ*, *ρ*) ∈ {*t_m_*, *t_f_*, *p_f_* }^2^ being the leading term of the selection coefficients *a_μρ_* calculated from the contribution to the next generation:

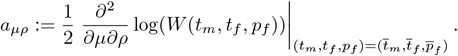

We obtain

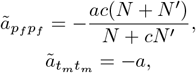

and

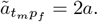

By summing Equations (A4) over each *i*, *j* in *T_m_* and *P* we obtain:

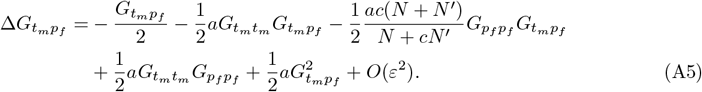

Under weak selection genetic correlations quickly reach equilibrium [Nagylaki, 1993]. For the sake of simplicity we assumed that the genetic correlations between traits and preferences are at equilibrium (as in [Barton and Turelli, 1991, Pomiankowski and Iwasa, 1993]). We obtain from (A5) that the two possible values at equilibrium are given by

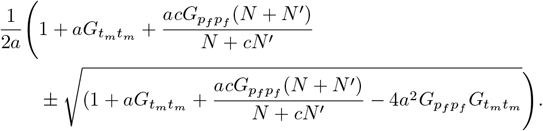

Only one of the two equilibrium values checks the Cauchy–Schwarz inequality 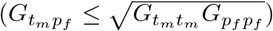. Therefore the equilibrium value is given by:

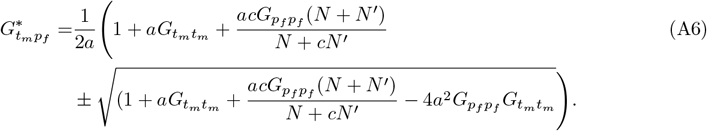

Because the genetic variance of traits and preferences is low, a Taylor expansion of (A6) gives

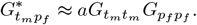

### 3 Low variance approximation

Because we assume that the variance of traits and preference is low we may use approximation in Equations (9), (15), (16) and (18). Here we detail how we obtained these approximations. The reasoning is similar for each approximation so we only explain how we get an approximation of 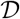 in (9). We recall that 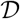 is defined by

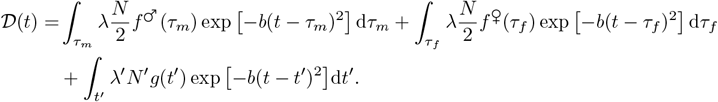

We first approximate the first term of 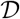. We have

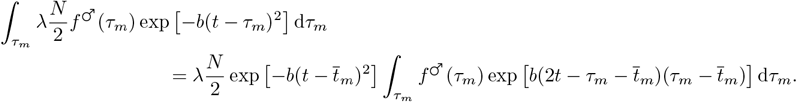

Using a Taylor expansion of 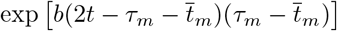 we have

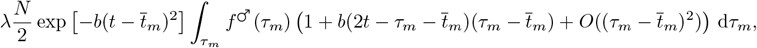

which is equal to

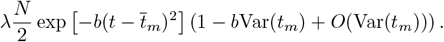

Hence when the variance of *t_m_* is low the first term of 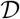 can be approximated by

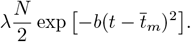

Similar computations for the other terms give the approximation in Equation (9).

### 4 Alternative scenarios

In the main document, we highlighted how the joint action of reproductive interference and predation may promote the evolution of FLM. We assumed that when the *focal* species enter in contact with *model*, reproductive interference and predation simultaneously exerted selection on individuals of the *focal* species (scenario 1). Here, we investigate the evolution of FLM under two other alternative scenarios. In scenario 2, we assume that the *focal* and the *model* species ancestrally shared common predators promoting mimicry, before sexual interactions happen between heterospecific individuals. In scenario 3, we assume the opposite sequences of events, whereby heterospecific sexual interactions occur before the two species start to share the same predators.

We compare the evolution of FLM under the three different scenarios using both the deterministic quantitative model (Figure A1) and individual-centred simulations assuming either independent genetic basis of male and female trait (Figure A2) or common genetic basis of male and female trait (Figure A3). Under scenario 2 (resp. 3) we let the traits in the *focal* species evolve with predation only (*d_m_* = *d_f_* > 0 and *c_RI_* = 0) (resp. reproductive interference only (*d_m_* = *d_f_* = 0 and *c_RI_* > 0)), until equilibrium using the deterministic quantitative model or after 10,000 generations using individual-centred simulations. Starting from the equilibria reached under each scenario, we assume that reproductive interference and predation then jointly influence the dynamics of traits in the *focal* species (*d_m_* = *d_f_* > 0 and *c_RI_* > 0). We compare the evolutionary outcomes observed when assuming either (1) that reproductive interference limits mimicry in males (*a* = 10) (Figure A1(a)(b)(c), Figure A2(a)(b)(c), Figure A3(a)(b)(c)) or (2) that reproductive interference promotes divergent evolution of male trait away from the ancestral value (*a* = 2.5) (Figure A1(d)(e)(f), Figure A2(d)(e)(f), Figure A3(d)(e)(f)).

Using the deterministic quantitative model, the three different scenarios leads to the same final male trait and female trait and preference values (Figure A1). Similarly, using individual-centred simulations male trait and female trait and preference values generally oscillate around the same value under the three scenarios (Figure A2 and A3), with few notable exceptions (Figure A4). When mimicry evolve first (scenario 2) male trait and female trait and preference values first oscillates around the trait displayed in the *model* species. If species enter sexually in contact when male trait is superior to the trait displayed in the *model* species, male trait increases and oscillates around a trait value that differs from the value observed under the other scenarios (Figure A4(b)).

**Figure A1:**
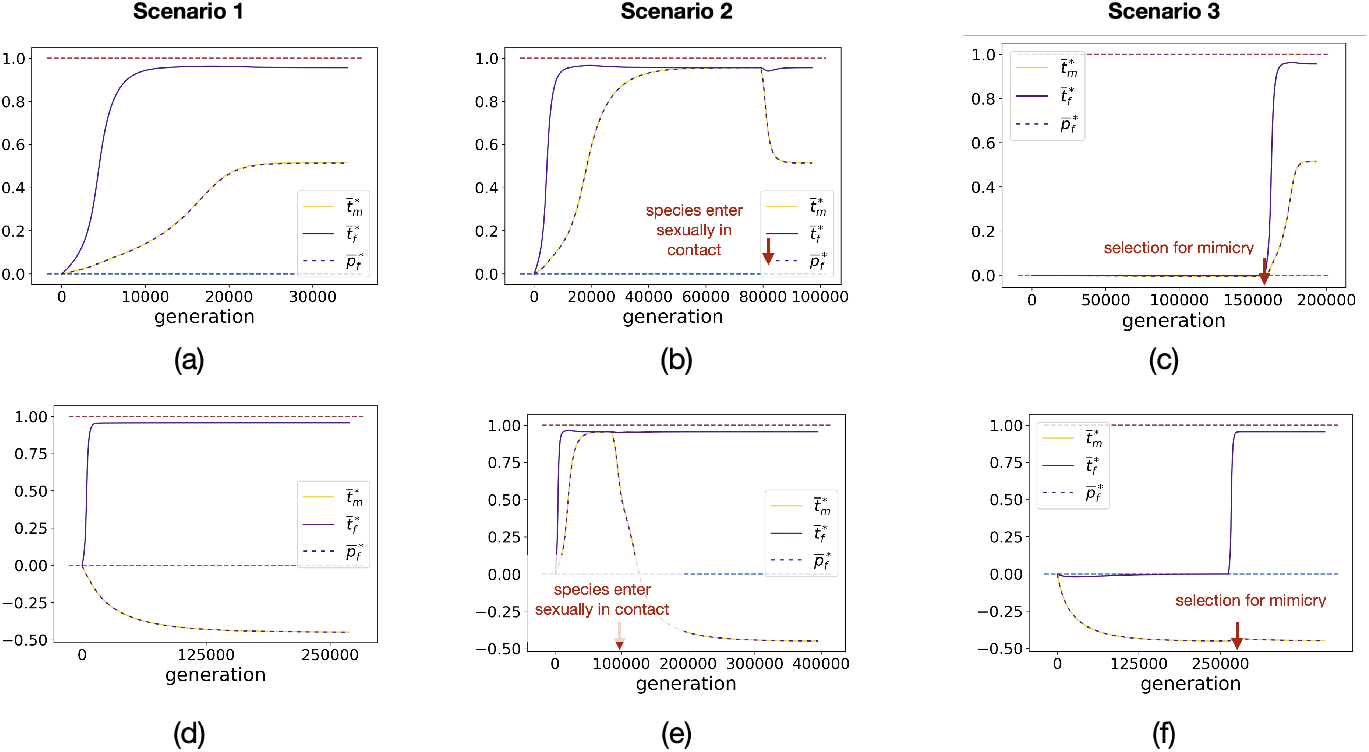
Effect of the history of species interactions on the dynamics of the mean males trait and females trait and preference values across generations given by the deterministic quantitative model. Different scenarios ((a)(d) simultaneous heterospecific sexual interactions and mimicry, (b)(e) initial mimicry, (c)(f) initial heterospecific sexual interactions) are explored when (a)(b)(c) reproductive interference limits mimicry in males (*a* = 10) and when (d)(e)(f) reproductive interference promotes divergent evolution of male trait away from the ancestral value (*a* = 2.5). We assume: *G_t_m__* = *G_t_f__* = *G_p_f__* = 0.01, *G_t_m_t_f__* = 0.001, *c* = 0.1, *c_RI_* = 0.01, *b* = 5, *d_m_* = *d_f_* = 0.05, λ = 0, *N* = 100, λ′ = 0.01, *N*′ = 200, *s* = 0.0025, *t_a_* = 0, 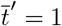.

**Figure A2:**
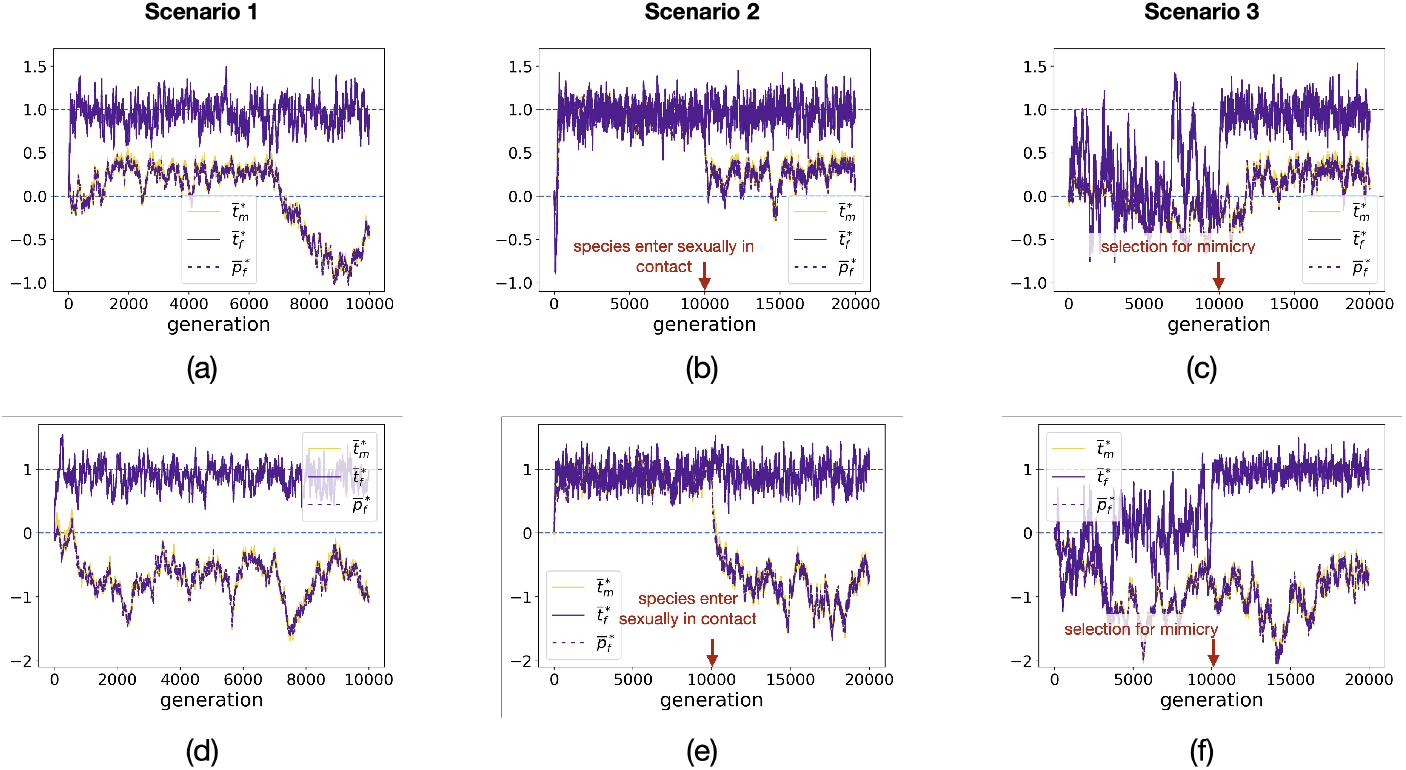
Effect of the history of species interactions on the dynamics of the mean males trait and females trait and preference values across generations given by individual-centred simulations assuming independent genetic basis of male and female trait. Different scenarios ((a)(d) simultaneous heterospecific sexual interactions and mimicry, (b)(e) initial mimicry, (c)(f) initial heterospecific sexual interactions) are explored when (a)(b)(c) reproductive interference limits mimicry in males (*a* = 10) and when (d)(e)(f) reproductive interference promotes divergent evolution of male trait away the ancestral value (*a* = 2.5). We assume: *G*_0_ = 0.0025, *μ* = 0.05, *r_T_m_T_f__* = 0.25, *r_T_f_P_f__* = 0.25, *c* = 0.1, **c_RI_** = 0.5, *b* = 5, *d_m_* = *d_f_* = 0.5, λ = 0, *N* = 100, λ′ = 0.01, *N*′ = 200, *s* = 0.025, *t_a_* = 0, 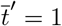.

**Figure A3:**
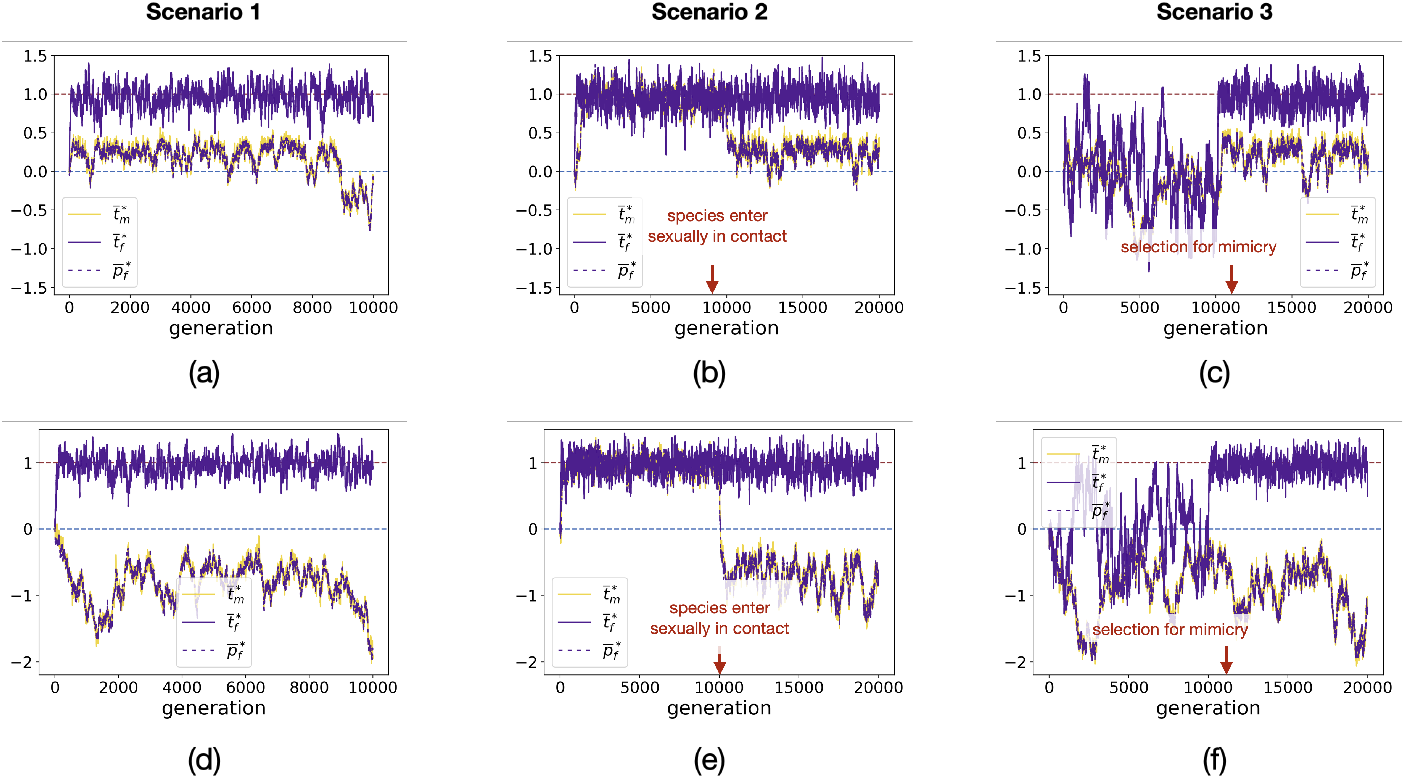
Effect of the history of species interactions on the dynamics of the mean males trait and females trait and preference values across generations given by individual-centred simulations assuming independent genetic basis of male and female trait. Different scenarios ((a)(d) simultaneous heterospecific sexual interactions and mimicry, (b)(e) initial mimicry, (c)(f) initial heterospecific sexual interactions) are explored when (a)(b)(c) reproductive interference limits mimicry in males (*a* = 10) and when (d)(e)(f) reproductive interference promotes divergent evolution of male trait away the ancestral value (*a* = 2.5). We assume: *G*_0_ = 0.0025, *μ* = 0.05, *r*_*T*_1_*T*_2__ = 0.25, *r*_*T*_2_*T*_3__ = 0.25, *r*_*T*_3_*P_f_*_ = 0.25, *c* = 0.1, *c_RI_* = 0.5, *b* = 5, *d_m_* = *d_f_* = 0.5, λ = 0, *N* = 100, λ′ = 0.01, *N*′ = 200, *s* = 0.025, *t_a_* = 0, 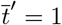.

**Figure A4:**
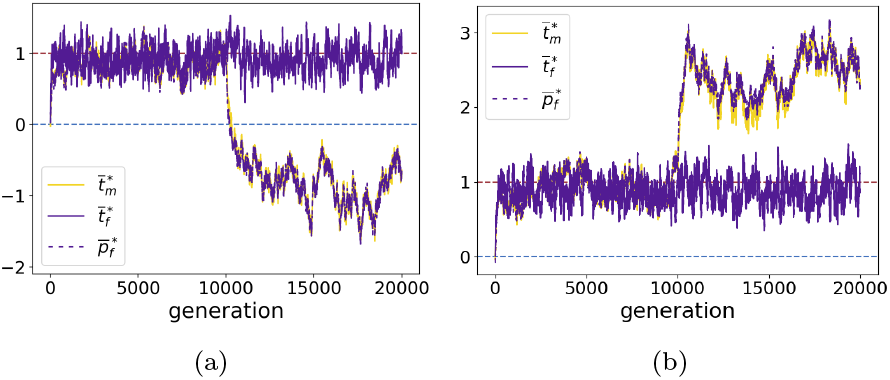
Two independent replicates of the dynamics of the mean males trait and females trait and preference values across generations given by individual-centred simulations assuming independent genetic basis of male and female trait when mimicry evolves first (scenario 2). We assume: *G*_0_ = 0.0025, *μ* = 0.05, *r_T_m_T_f__* = 0.25, *r_T_f_P_f__* = 0.25, *c* = 0.1, *a* = 2.5, *c_RI_* = 0.5, *b* = 5, *d_m_* = *d_f_* = 0.5, λ = 0, *N* = 100, λ′ = 0.01, *N*′ = 200, *s* = 0.025, *t_a_* = 0, 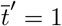.

### 5 Sexually contrasted predation promotes higher trait divergence in females

In this part, we show that if FLM in a palatable species (λ = 0) is not caused by sexual selection (*a* = 0) but by sexually contrasted predation (*d_f_* > *d_m_*) then at the final state females trait 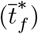 diverges more from the ancestral trait than male trait 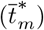. In mathematical terms, we prove that if *a* = 0 and *d_f_* > *d_m_* we have

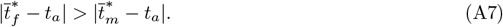

For simplicity we assume that *t*′ > *t_a_*, the other case being obtained by symmetry.

At final state we have 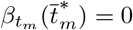 (*β_t_m__* is given in Equation (3). Because we have

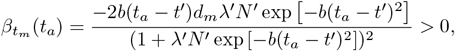

and

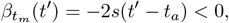

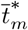 is bounded by *t_a_* and *t*′. Similar arguments give that final females trait is bounded by *t_a_* and *t*′. Because 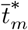 is the final trait we have 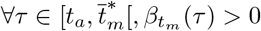.

For all trait *τ* we have

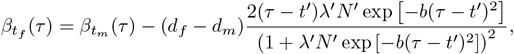

which implies that 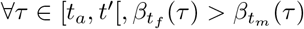. Then 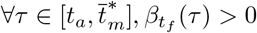. Therefore 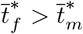 and then we have (A7).

### 6 Temporal dynamics of sexual dimorphism

Here, we illustrate the temporal dynamics of sexual dimorphism when

- reproductive interference limits mimicry in males (Figure A5(a)).
- reproductive interference promotes divergence from the ancestral trait in males (Figure A5(b)).
- sexually contrasted predation promotes mimicry in females only (Figure A5(c)).

**Figure A5:**
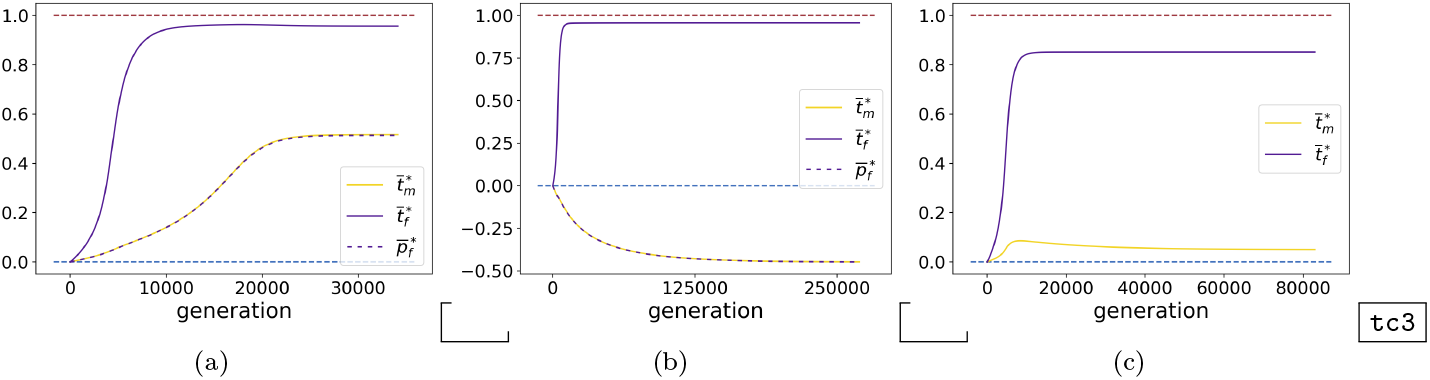
Evolution of the mean males trait and females trait and preference values across generations (a)(b) when reproductive interference or (c) sexually contrasted predation promotes sexual dimorphism. We assume: (a) *c_RI_* = 0.01, *a* = 10, *s* = 0.0025, *d_m_* = 0.05, (b) *c_RI_* = 0.01, *a* = 2.5, *s* = 0.0025, *d_m_* = *d_f_* = 0.05 (c) *c_RI_* = 0, *a* = 0, *s* = 0.01, *d_m_* = 0.005. We assume for the other parameters: *G_t_m__* = *G_t_f__* = *G_p_f__* = 0.01, *G_t_m_t_f__* = 0.001, *c* = 0.1, *b* = 5, *d_f_* = 0.05, λ = 0, *N* = 100, λ′ = 0.01, *N*′ = 200, *t_a_* = 0, 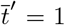. The curves stop when the males trait and females trait and preference values reach equilibrium.

### 7 Reproductive interference promotes female-limited mimicry in palatable species when females have sufficiently low cost of choosiness

The evolution of FLM strongly depends on the evolution of females preference. As we have already seen the evolution of females preference depends on reproductive interference promoting preferences for non-mimetic males. However such preferences may cause females to seek for rarer males in the population. The evolution of preference limiting the cost of reproductive interference may thus be limited by the cost of choosiness described by the parameter *c*. We thus investigate the impact of the strength of reproductive interference (*c_RI_*) promoting FLM and the cost of choosiness (*c*) on the final level of sexual dimorphism given by 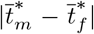 (Figure A6 (a)) and on final females preference 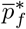 (Figure A6 (b)). Cost of choosiness limits the evolution of sexual dimorphism due to reproductive interference (Figure A6 (a)) because it limits the evolution of females preference (Figure A6 (b)). In natural population, reproductive interference may explain FLM in populations where females have low cost of choosiness.

**Figure A6:**
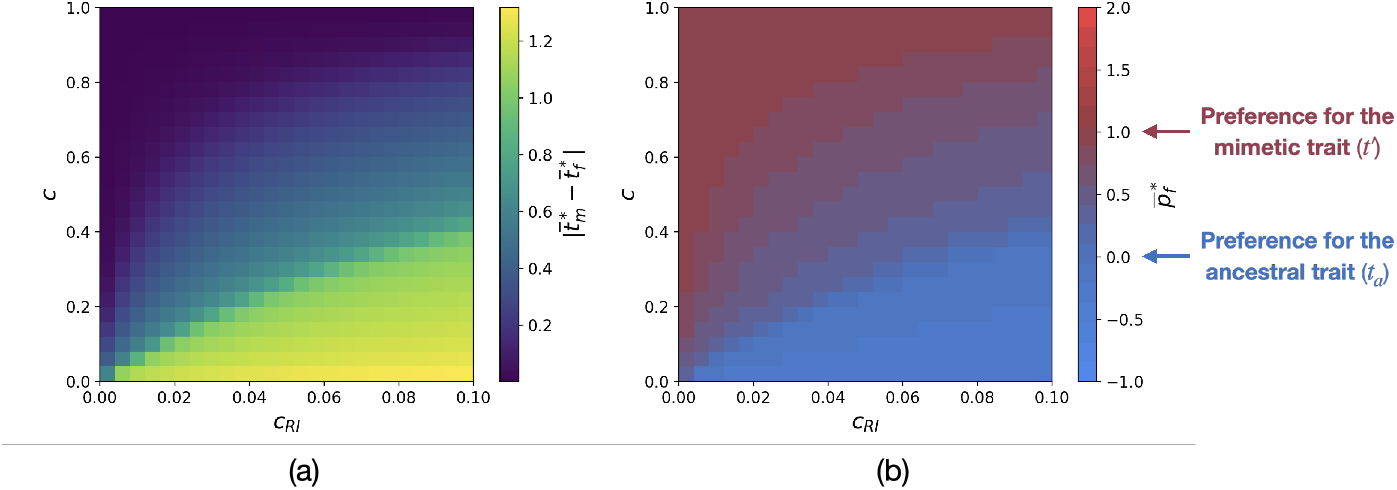
Influence of the strength of reproductive interference *c_RI_* and of the cost of choosiness *c* on the final level of sexual dimorphism 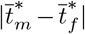 and final preference 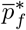. We assume: *G_t_m__* = *G_t_f__* = *G_p_f__* = 0.01, *G_t_m_t_f__* = 0.001, *a* = 5, *b* = 5, *d_m_* = *d_f_* = 0.05, λ = 0, *N* = 100, λ′ = 0.01, *N*′ = 200, *s* = 0.0025, *t_a_* = 0, 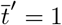.

### 8 Impact of the genetic correlation between males and females traits *C_t_m_t_f__*

The evolution of the mean males and females trait values (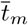 and 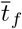) depends on the genetic covariance between males and females traits (*G_t_m_t_f__*) (see equation (2)). We investigate the impact of this genetic covariance and of the strength of reproductive interference (*c_RI_*) on the level of sexual dimorphism (Figure A7). The level of sexual dimorphism is not impacted by the genetic covariance unless this quantity is at its maximum value 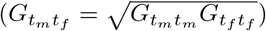. Indeed when the genetic covariance is at it maximum value males and females traits have the same genetic basis, therefore the evolution of sexual dimorphism is not possible. By contrast when males and females traits have at least partially different genetic basis 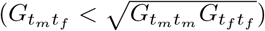 the non-shared genetic basis allows the level of sexual dimorphism to increase.

**Figure A7:**
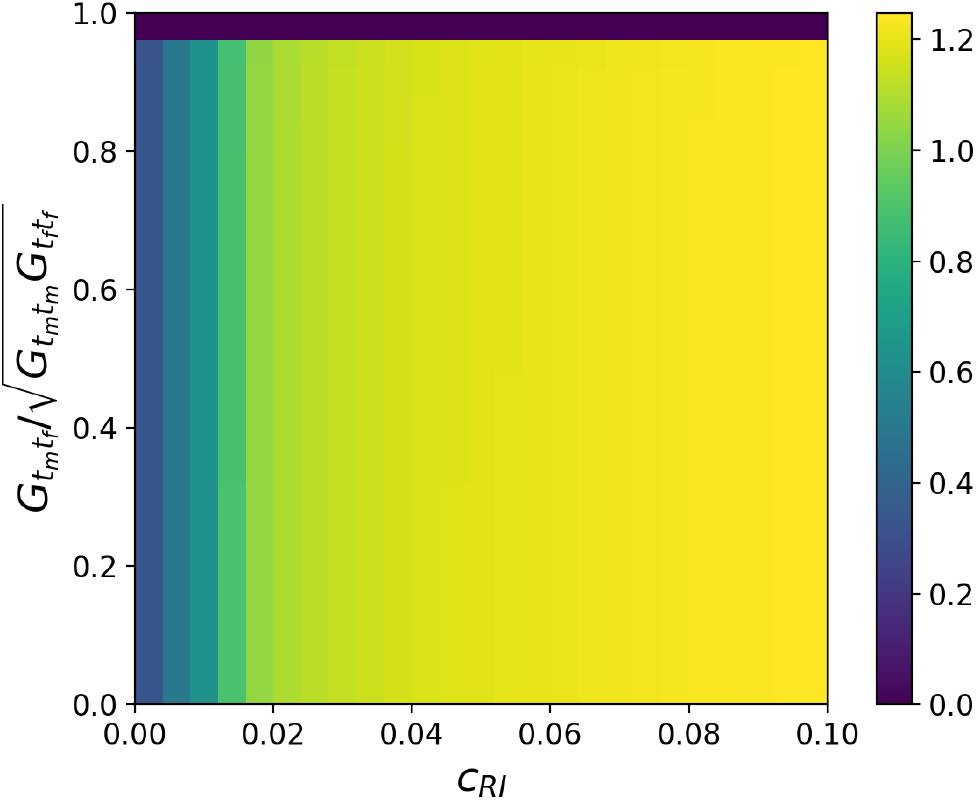
Influence of the strength of reproductive interference *c_RI_* and of the genetic covariance between males and females traits normalized by its maximum value 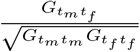 on the final level of sexual dimorphism 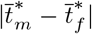. We assume: *G_t_m__* = *G_t_f__* = *G_p_f__* = 0.01, *c* = 0.1, *a* = 5, *b* = 5, *d_m_* = *d_f_* = 0.05, λ = 0, *N* = 100, λ′ = 0.01, *N*′ = 200, *s* = 0.0025, *t_a_* = 0, 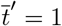.

However *G_t_m_t_f__* impacts the speed at which the equilibrium is reached. When males trait in the *focal* species gets closer to the mimetic trait the genetic correlation increases the speed of convergence because selection on females trait also favours mimicry and also acts on males trait. By contrast when males trait diverges away from the mimetic trait the genetic correlation decreases the speed of convergence.

**Figure A8:**
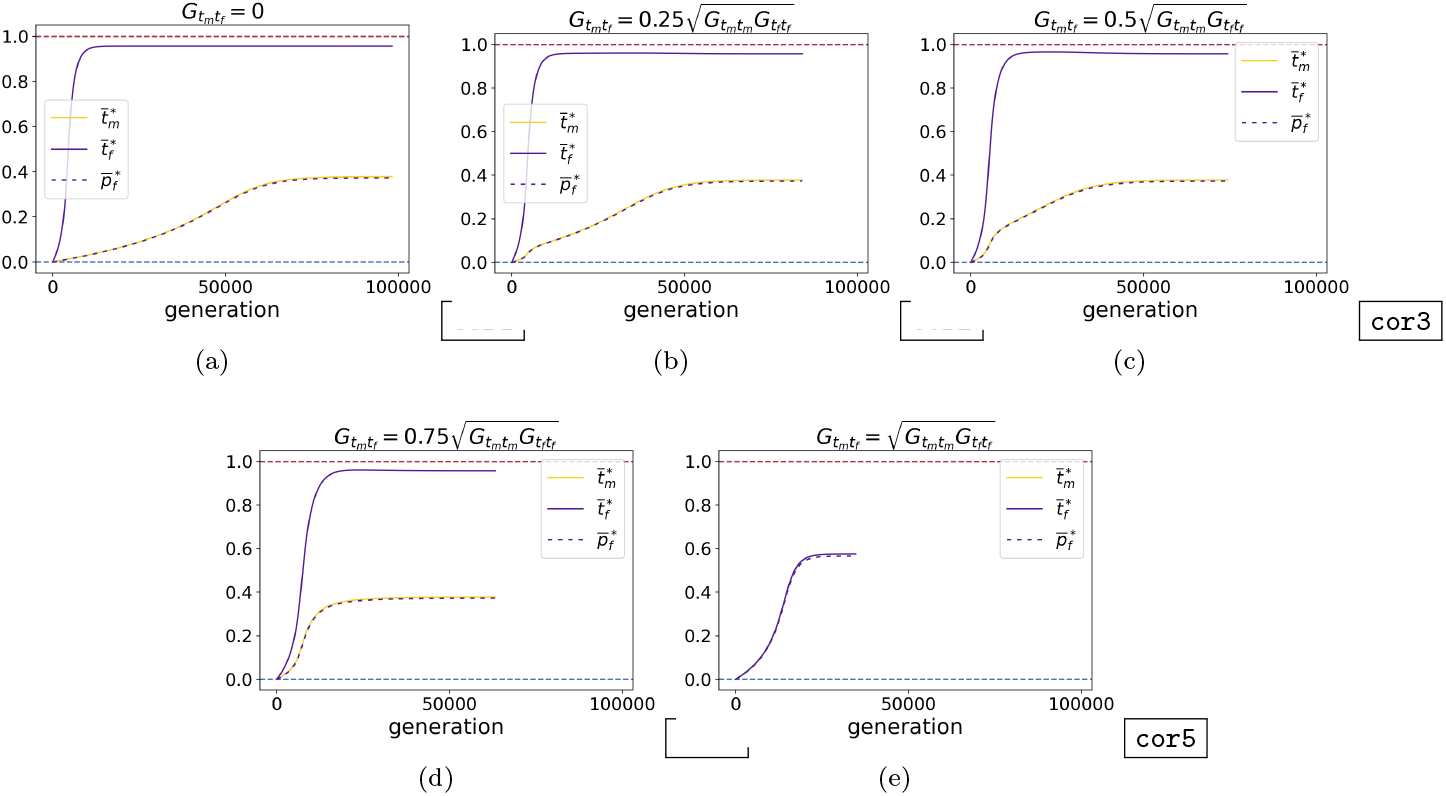
Evolution of the mean males trait and females trait and preference values across generations for different genetic covariances between males and females traits *G_t_m_t_f__* when males trait gets closer to the mimetic trait. We assume different values of the genetic covariance between male and female traits: (a) *G_t_m_t_f__* = 0, (b) 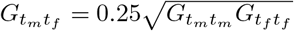, (c) 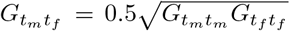, (d) 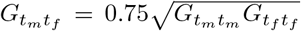, (e) 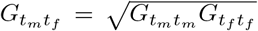. We assume: *G_t_m__* = *G_t_f__* = *G_p_f__* = 0.01, *G_t_m_t_f__* = 0.001, *c_RI_* = 0.01, *c* = 0.1, *a* = 5, *b* = 5, *d_m_* = *d_f_* = 0.05, λ = 0, *N* = 100, λ′ = 0.01, *N*′ = 200, *s* = 0.0025, *t_a_* = 0, 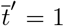. The curves stop when the males trait and females trait and preference values reach equilibrium.

**Figure A9:**
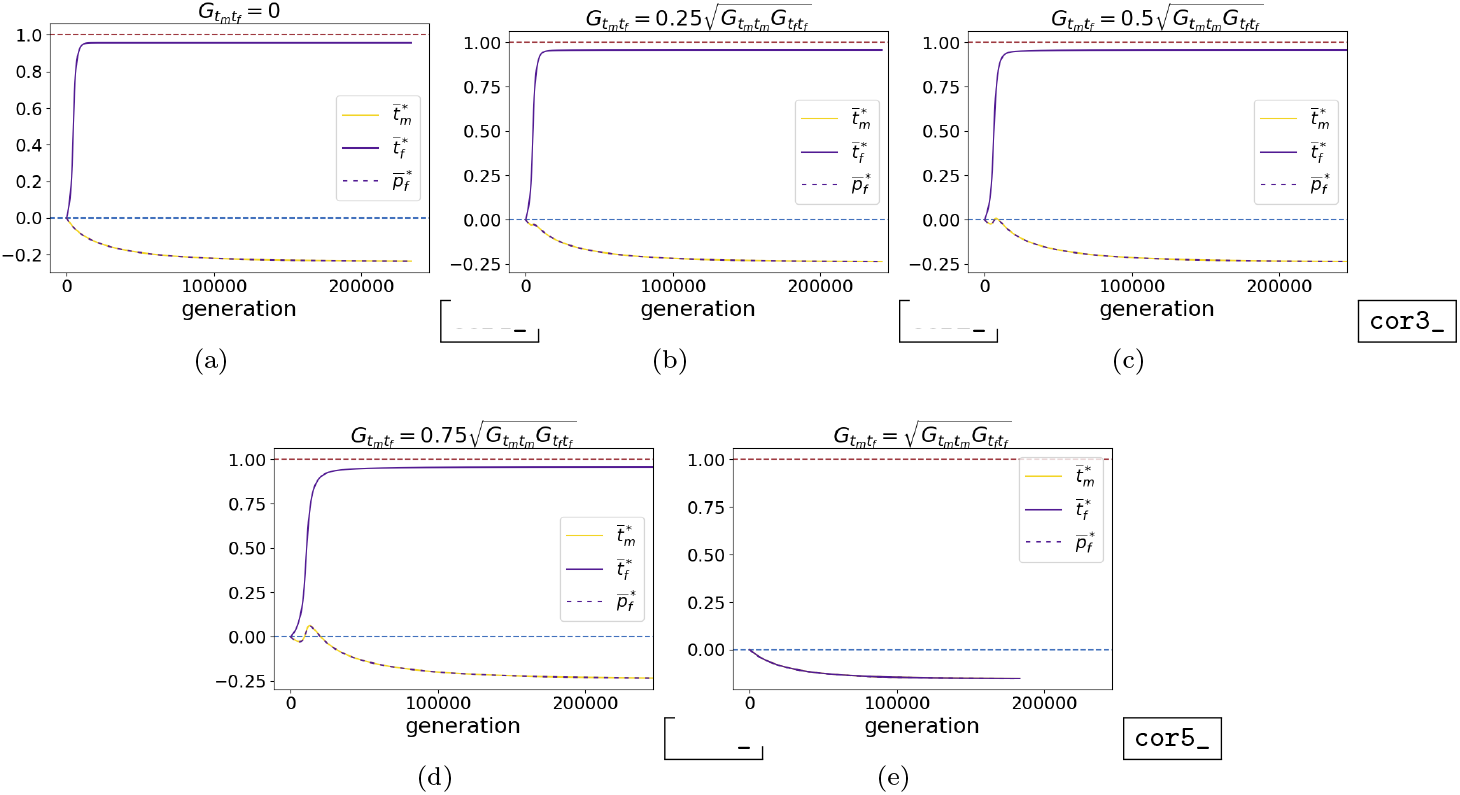
Evolution of the mean males trait and females trait and preference values across generations for different genetic covariances between male and female traits *G_t_m_t_f__* when reproductive interference promotes divergence of males trait away from the mimetic trait. We assume different value of the genetic covariance between of male and female trait: (a) *G_t_m_t_f__* = 0, (b) 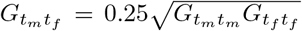, (c) 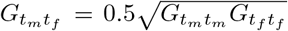, (d) 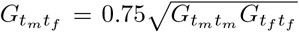, (e) 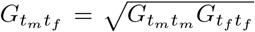. We assume: *G_t_m__* = *G_t_f__* = *G_p_f__* = 0.01, *G_t_m_t_f__* = 0.001, *c_RI_* = 0.05, *c* = 0.1, *a* = 5, *b* = 5, *d_m_* = *d_f_* = 0.05, λ = 0, *N* = 100, λ′ = 0.01, *N*′ = 200, *s* = 0.0025, *t_a_* = 0, 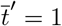.

### 9 Investigation of the effect of reproductive interference on the evolution of FLM using individual-centred simulations

**Figure A10:**
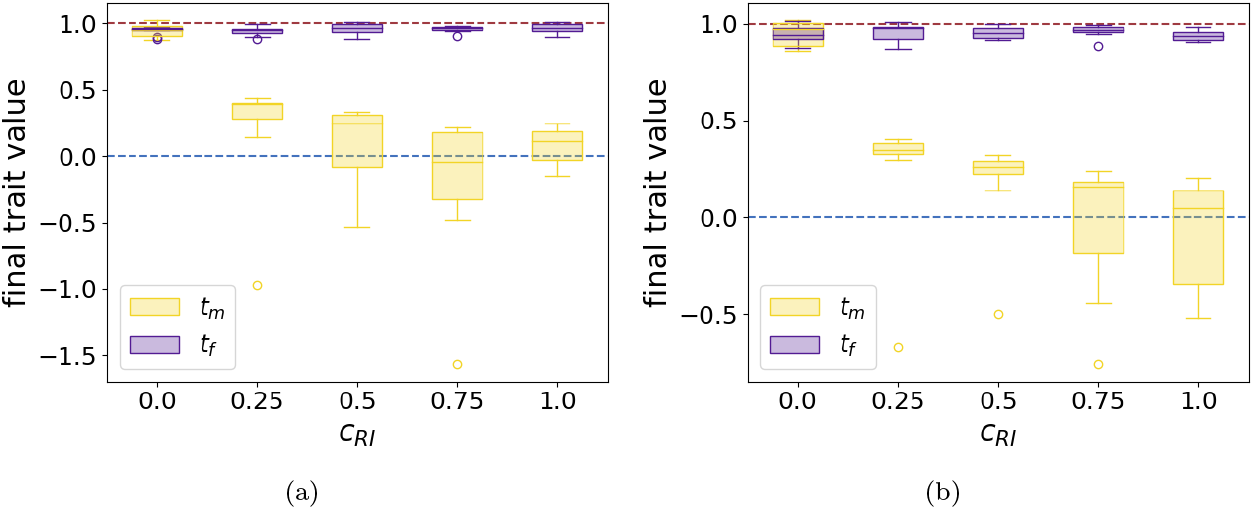
Boxpolts of final mean male (yellow) and female (purple) traits values for different strength of reproductive interference *c_RI_* using individual-centred simulations assuming (a) independent genetic basis or (b) partially common genetic basis of male and female trait. We assume: (a) *r_T_m_T_f__* = 0.25, *r_T_f_ P_f__* = 0.25 and (b) *r*_*T*_1_*T*_2__ = 0.25, *r*_*T*_2_*T*_3__ = 0.25, *r*_*T*_3_*P_f_*_ = 0.25. We also assume: *G*_0_ = 0.0025, *μ* = 0.05, *c* = 0.1, *a* = 10, *b* = 5, *d_m_* = *d_f_* = 0.5, λ = 0, *N* = 100, λ′ = 0.01, *N*′ = 200, *s* = 0.025, *t_a_* = 0, 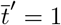.

**Figure A11:**
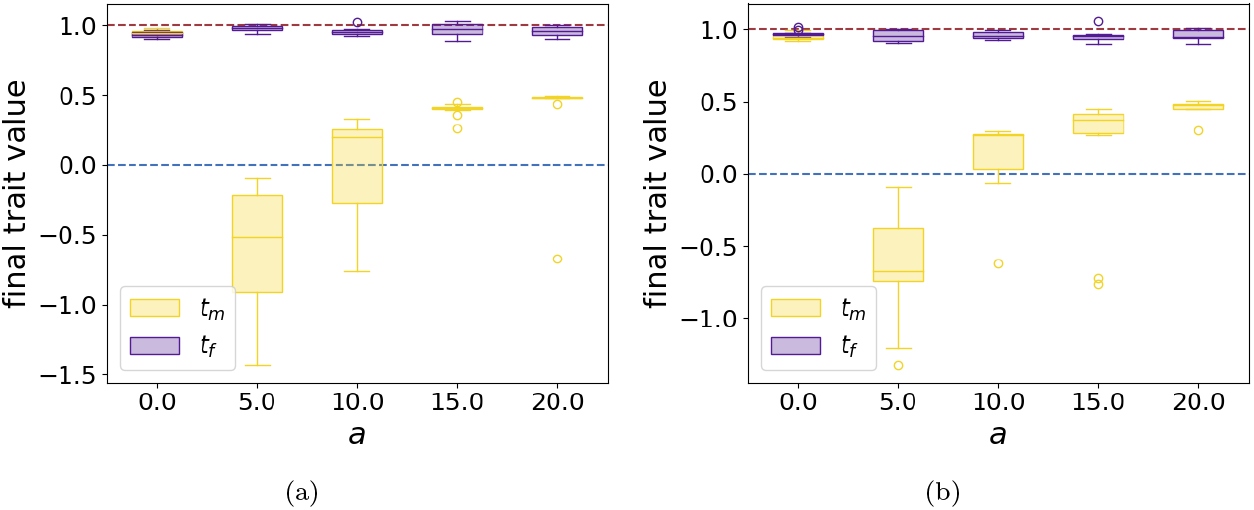
Boxpolts of final mean male (yellow) and female (purple) traits values for different females choosiness *a* using individual-centred simulations assuming (a) independent genetic basis or (b) partially common genetic basis of male and female trait. We assume: (a) *r_T_m_T_f__* = 0.25, *r_T_f_ P_f__* = 0.25 and (b) *r*_*T*_1_*T*_2__ = 0.25, *r*_*T*_2_*T*_3__ = 0.25, *r*_*T*_3_*P_f_*_ = 0.25. We also assume: *G*_0_ = 0.0025, *μ* = 0.05, *c* = 0.1, *c_RI_* = 0.5, *b* = 5, *d_m_* = *d_f_* = 0.5, λ = 0, *N* = 100, λ′ = 0.01, *N*′ = 200, *s* = 0.025, *t_a_* = 0, 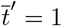.

### 10 Investigation of the effect of sexually contrasted predation on the evolution of FLM using individual-centred simulations

**Figure A12:**
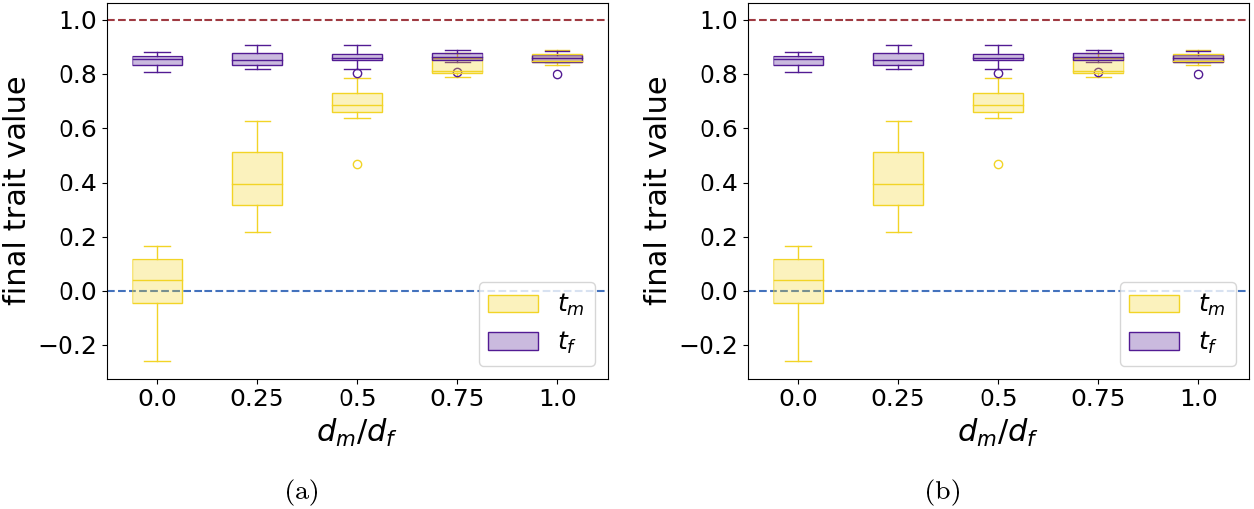
Boxpolts of final mean male (yellow) and female (purple) traits values for different ratio of basal predation rate on males and females *d_m_/d_f_* using individual-centred simulations assuming either (a) independent genetic basis or (b) partially common genetic basis of male and female trait. We assume: (a) *r_T_m_T_f__* = 0.25, *r_T_f_ P_f__* = 0.25 and (b) *r*_*T*_1_*T*_2__ = 0.25, *r*_*T*_2_*T*_3__ = 0.25, *r*_*T*_3_*P_f_*_ = 0.25. We also assume: *G*_0_ = 0.0025, *μ* = 0.05, *c* = 0, *a* = 0, *c_RI_* = 0, *b* = 5, *d_f_* = 0.5, λ = 0, *N* = 100, λ′ = 0.01, *N*′ = 200, *s* = 0.1, *t_a_* = 0, 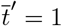.

**Figure A13:**
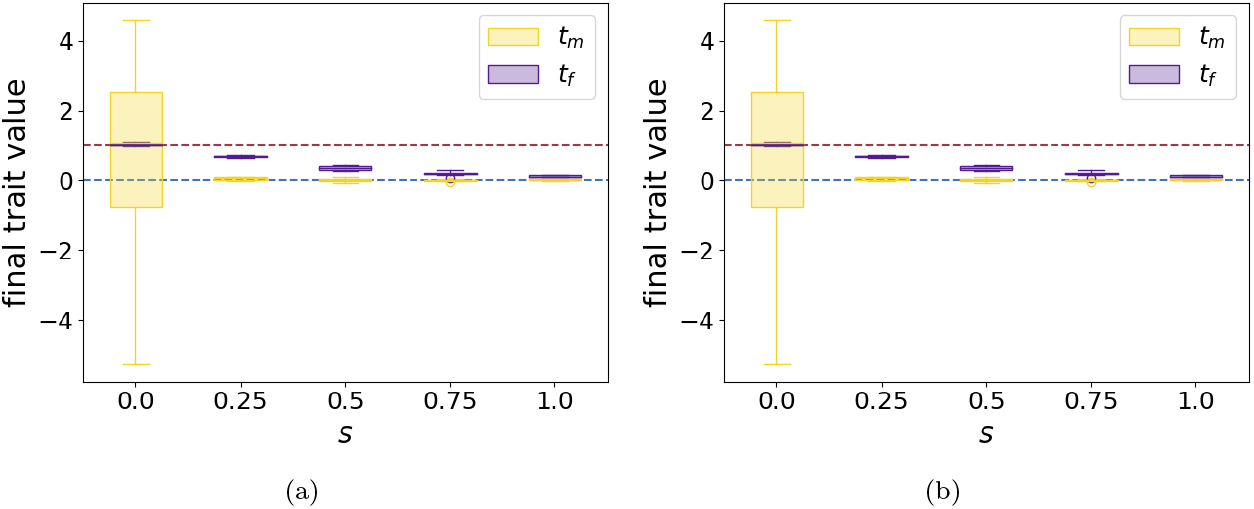
Boxpolts of final mean male (yellow) and female (purple) traits values for different strength of developmental constraints *s* using individual-centred simulations assuming either (a) independent genetic basis or (b) partially common genetic basis of male and female trait. We assume: (a) *r_T_m_T_f__* = 0.25, *r_T_f_ P_f__* = 0.25 and (b) *r*_*T*_1_*T*_2__ = 0.25, *r*_*T*_2_*T*_3__ = 0.25, *r*_*T*_3_*P_f_*_ = 0.25. We also assume: *G*_0_ = 0.0025, *μ* = 0.05, *c* = 0, *a* = 0, *c_RI_* = 0, *b* = 5, *d_m_* = 0.05, *d_f_* = 0.5, λ = 0, *N* = 100, λ′ = 0.01, *N*′ = 200, *t_a_* = 0, 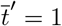.

### 11 Exploring the relative divergence of males and females from the ancestral trait using individual-centred simulations

#### 11.1 FLM caused by reproductive interference

**Figure A14:**
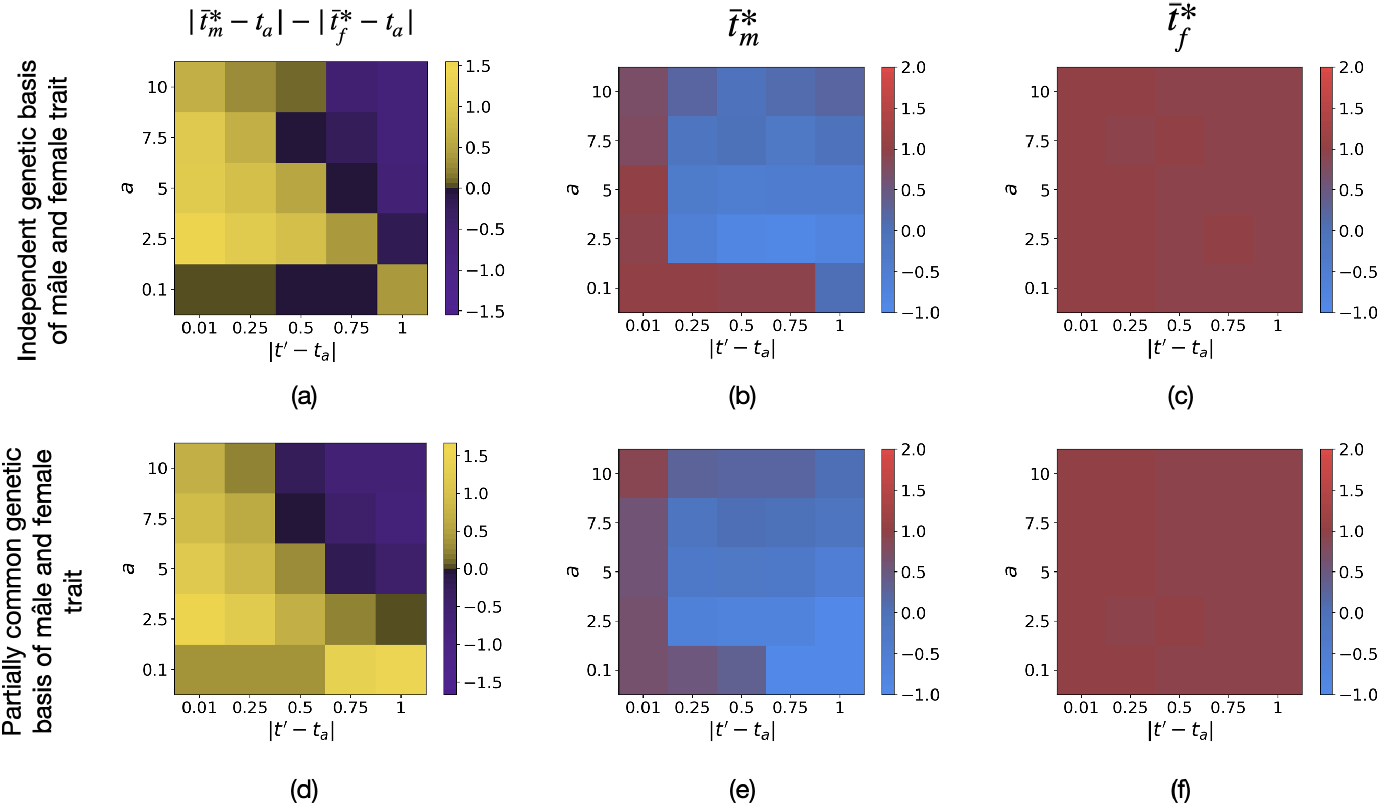
Influence of the distance between the ancestral and the mimetic traits |*t*′–*t_a_*| and of females choosiness *a* on (a)(d) the difference between the level of divergence in males and females 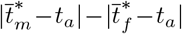, (b)(e) final male trait 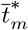 and (c)(f) final female trait 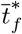 using individual-centred simulations assuming either (a)(b)(c) independent genetic basis or (d)(e)(f) partially common genetic basis of male and female trait. We assume: (a) *r_T_m_T_f__* = 0.25, *r_T_f_ P_f__* = 0.25 and (b) *r*_*T*_1_*T*_2__ = 0.25, *r*_*T*_2_*T*_3__ = 0.25, *r*_*T*_3_*P_f_*_ = 0.25. We also assume: *G*_0_ = 0.0025, *μ* = 0.05, *c* = 0.1, *c_RI_* = 0.5, *b* = 5, *d_m_* = *d_f_* = 0.5, λ = 0, *N* = 100, λ′ = 0.01, *N*′ = 200, *s* = 0.025, 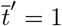.

The deterministic quantitative model and individuals-centred simulations show the same impact of the distance between the ancestral and the mimetic traits |*t*′ – *t_a_* | and of females choosiness a on (a)(d) the difference between the level of divergence in males and females 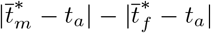 (Figures 6(c) and A14(a)(d)). However, when the ancestral trait is close to the trait displayed in the *model* species (*t_a_* = 0.99, *t*′ = 1), the different models then predict a different evolution of mean male trait value:

- Using the deterministic quantitative model, male traits value diverge from the mimetic trait towards the ancestral trait value (Figure 6(a)).
- Using individuals-centred simulations, final male trait values are centred around the mimetic trait (Figure A14(b)(e)). Male traits also diverge but not necessarily toward the ancestral trait because stochasticity allows male trait to reach higher values than the mimetic trait value (Figure A16).

is centred around the mimetic trait (*t*′) using individuals-centred simulations whereas male trait using thedeterministic quantitative model.

**Figure A15:**
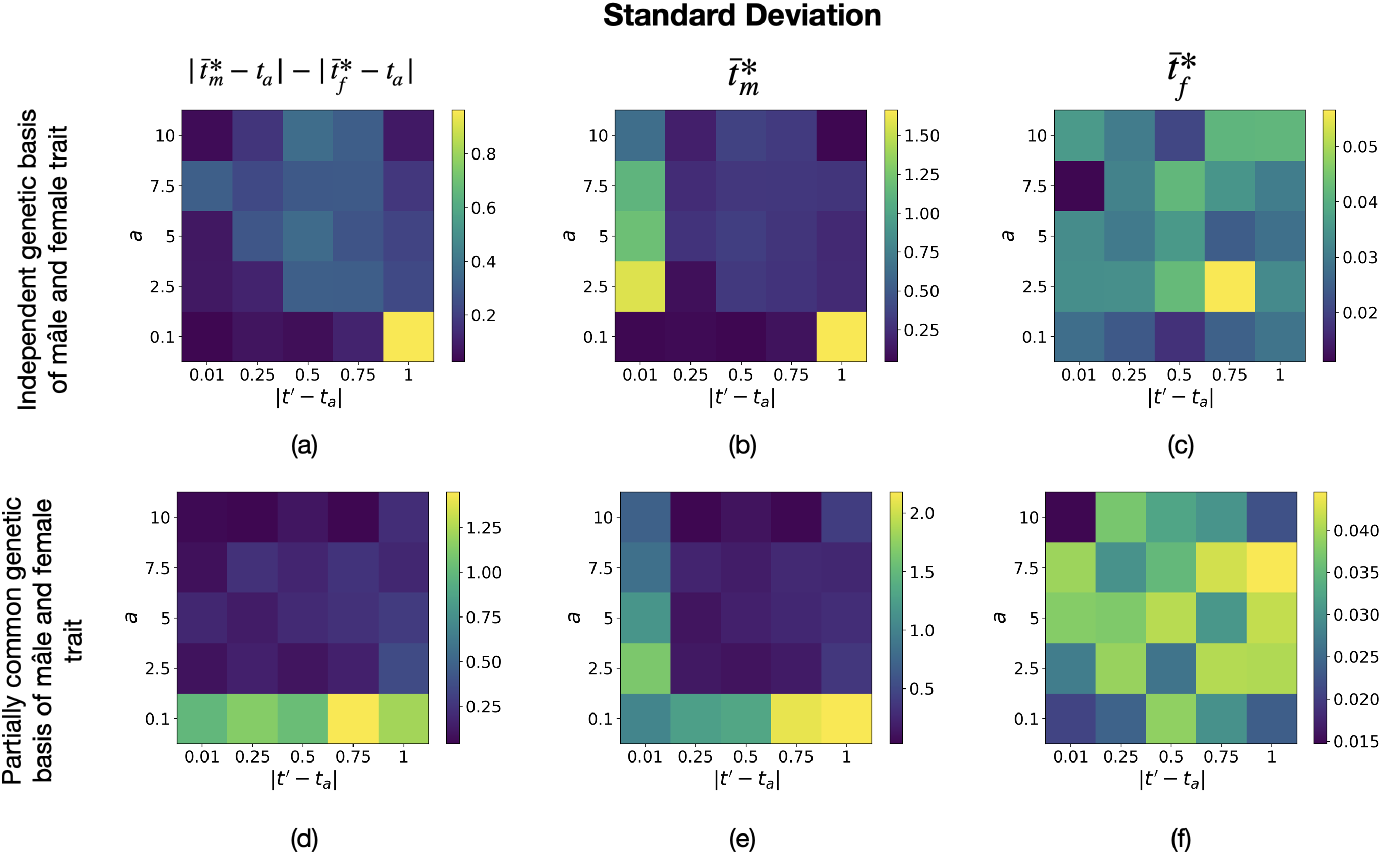
Standard deviation associated with Figure A14 of (a)(d) the difference between the level of divergence in males and females 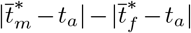, (b)(e) final male trait 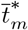 and (c)(f) final female trait 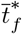 using individual-centred simulations assuming (a)(b)(c) independent genetic basis or (d)(e)(f) partially common genetic basis of male and female trait.

**Figure A16:**
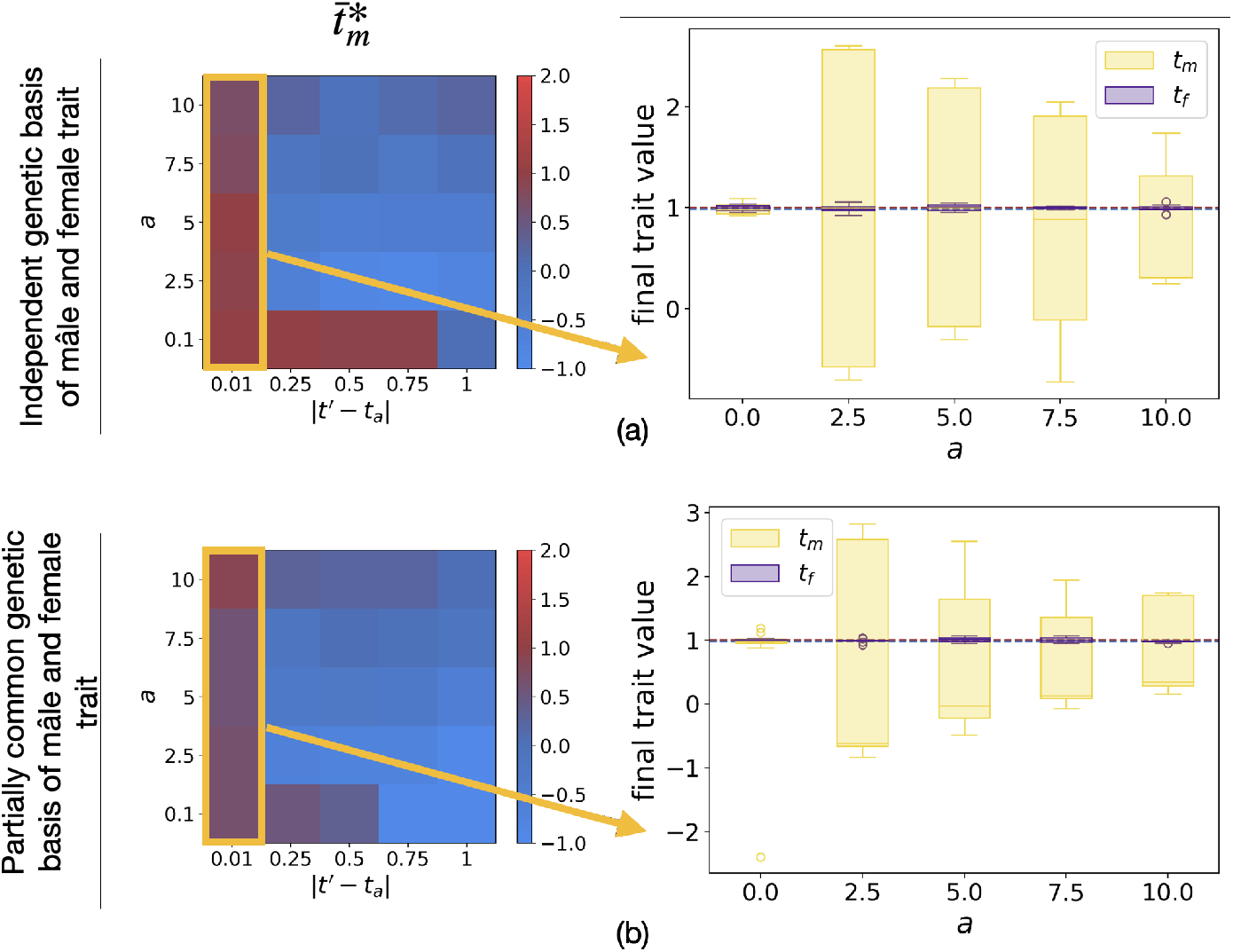
Boxplots of final mean male (yellow) and female (purple) traits values for different females choosiness *a* using individual-centred simulations assuming either (a) independent genetic basis or (b) partially common genetic basis of male and female trait. We assume: (a) *r_t_m_t_f__* = 0.25, *r_T_f_ P_f__* = 0.25 and (b) *r*_*T*_1_*T*_2__ = 0.25, *r*_*T*_2_*T*_3__ = 0.25, *r*_*T*_3_*P_f_*_ = 0.25. We also assume: *G*_0_ = 0.0025, *μ* = 0.05, *c* = 0.1, *c_RI_* = 0.5, *b* = 5, *d_m_* = *d_f_* = 0.5, λ = 0, *N* = 100, λ′ = 0.01, *N*′ = 200, *s* = 0.025, *t_a_* = 0.99, 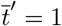.

#### 11.2 FLM caused by sexually contrasted predation

**Figure A17:**
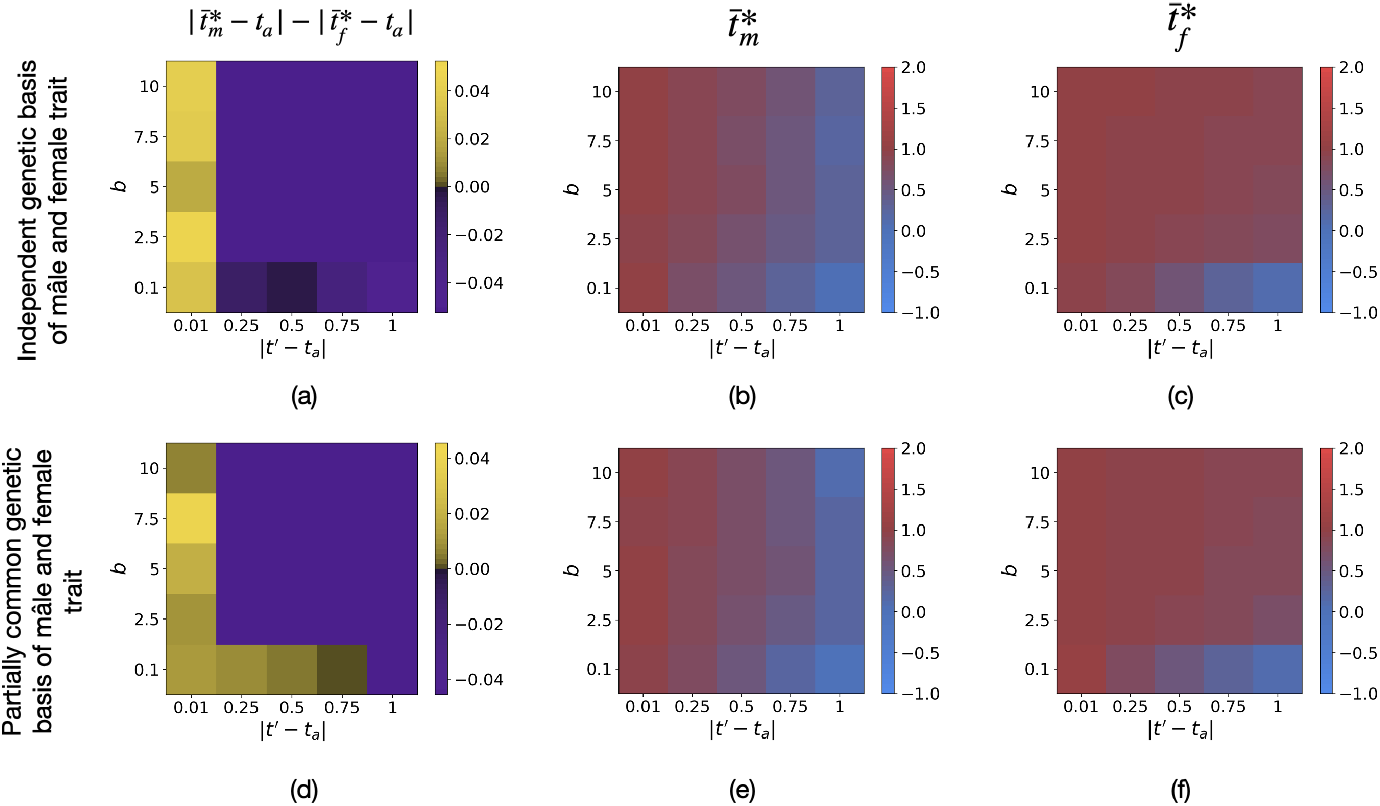
Influence of the distance between the ancestral and the mimetic traits |*t*′ – *t_a_*| and of predators discrimination *b* on (a)(d) the difference between the level of divergence in males and females 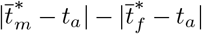, (b)(e) final male trait 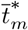 and (c)(f) final female trait 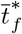 using individual-centred simulations assuming either (a)(b)(c) independent genetic basis or (d)(e)(f) partially common genetic basis of male and female trait. We assume: (a) *r_T_m_T_f__* = 0.25, *r_T_f_P_f__* = 0.25 and (b) *r*_*T*_1_*T*_2__ = 0.25, *r*_*T*_2_*T*_3__ = 0.25, *r*_*T*_3_*P_f_*_ = 0.25. We also assume: *G*_0_ = 0.0025, *μ* = 0.05, *c* = 0, *a* = 0, *c_RI_* = 0, *d_m_* = 0.1, *d_f_* = 0.5, λ = 0, *N* = 100, λ′ = 0.01, *N*′ = 200, *s* = 0.1, 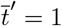.

**Figure A18:**
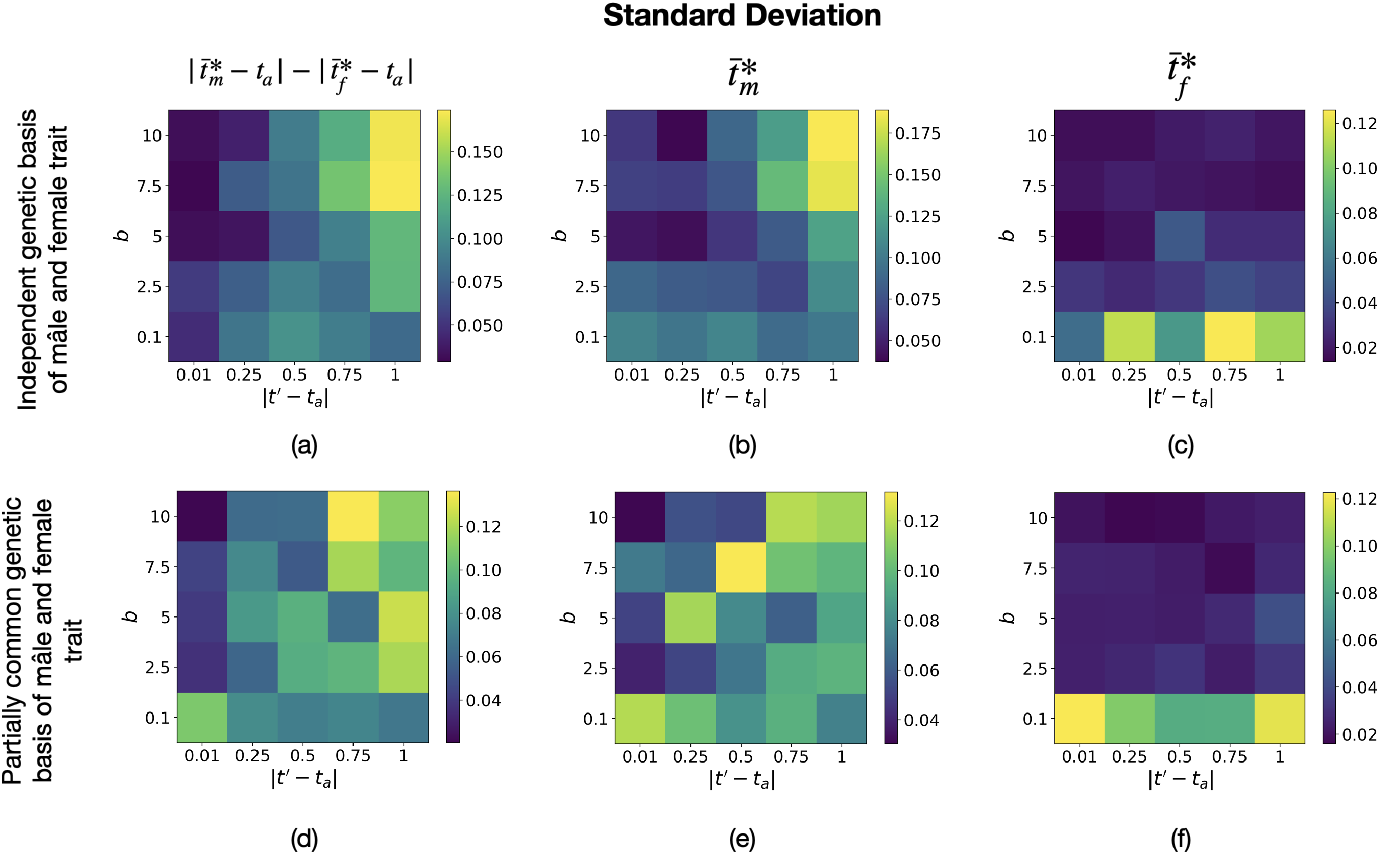
Standard deviation associated with Figure A17 of (a)(d) the difference between the level of divergence in males and females 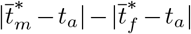, (b)(e) final male trait 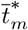 and (c)(f) final female trait 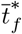 using individual-centred simulations assuming either (a)(b)(c) independent genetic basis or (d)(e)(f) partially common genetic basis of male and female trait.

**Figure A19:**
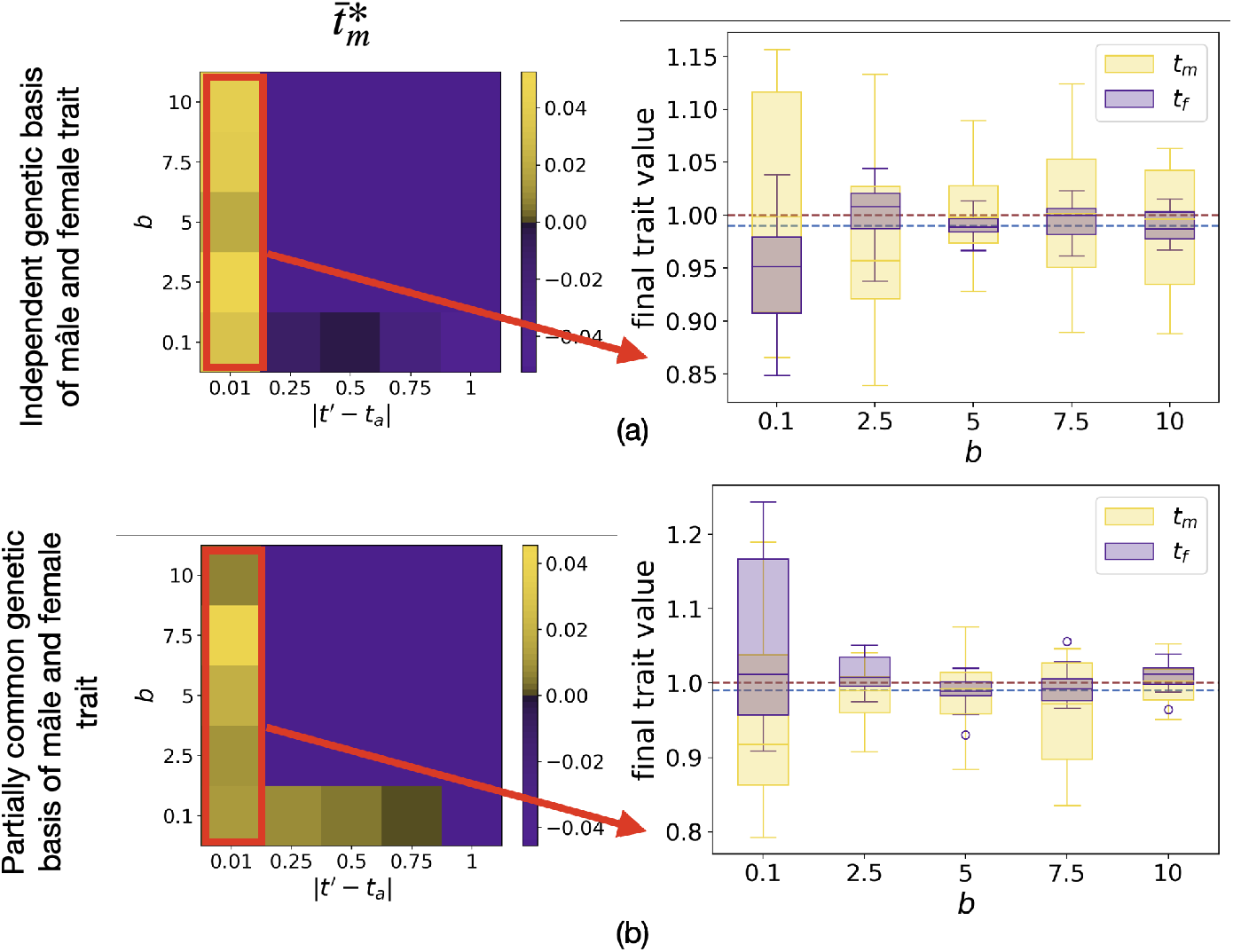
Boxplots of final mean male (yellow) and female (purple) traits values for different females choosiness *a* using individual-centred simulations assuming either (a) independent genetic basis or (b) partially common genetic basis of male and female trait. We assume: (a) *r_T_m_T_f__* = 0.25, *r_T_f_ P_f__* = 0.25 and (b) *r*_*T*_1_ *T*_2__ = 0.25, *r*_*T*_2_*T*_3__ = 0.25, *r*_*T*_3_*P_f_*_ = 0.25. We also assume: *G*_0_ = 0.0025, *μ* = 0.05, *c* = 0, *a* = 0, *c_RI_* = 0, *d_m_* = 0.1, *d_f_* = 0.5, λ = 0, *N* = 100, λ′ = 0.01, *N*′ = 200, *s* = 0.1, *t_a_* = 0.99, 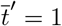.

### 12 Additional figures: The evolution of FLM depends on defence level

**Figure A20:**
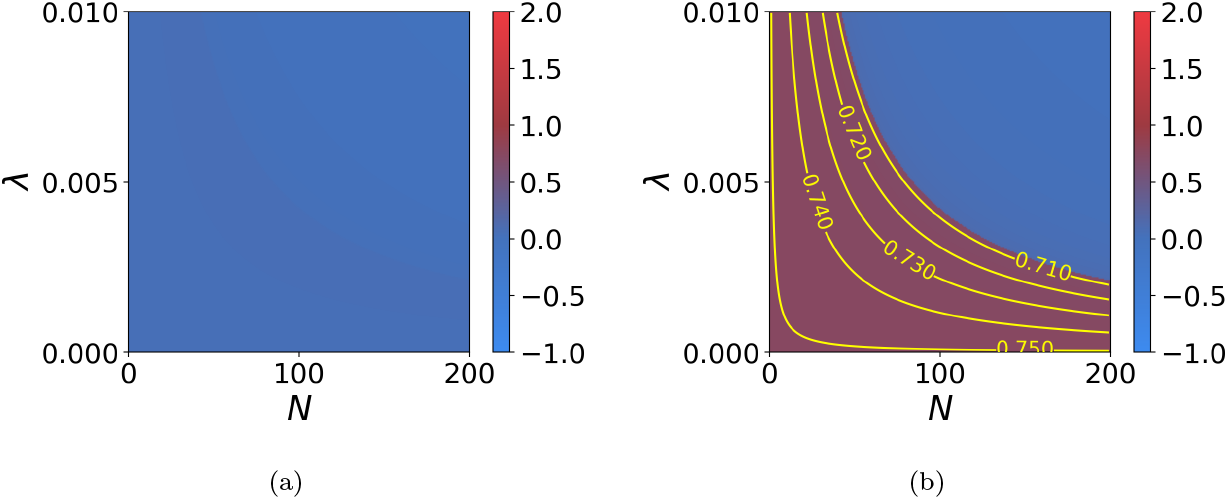
Influence of the density *N* and of the individual defence level λ in the *focal* species on the equilibrium values of (a) males trait 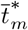 and (b) females trait 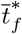 when female-limited mimicry is caused by sexually contrasted predation (*d_f_* > *d_m_*, *a* = 0). Yellow lines indicate equal trait value. We assume: *G_t_m__* = *G_t_f__* = *G_p_f__* = 0.01, *G_t_m_t_f__* = 0.001, *c_RI_* = 0, *c* = 0, *a* = 0, *b* = 5, *d_m_* = 0.01, *d_f_* = 0.05, λ′ = 0.01, *N*′ = 200, *s* = 0.02, *t_a_* = 0, *t*′ = 1.

**Figure A21:**
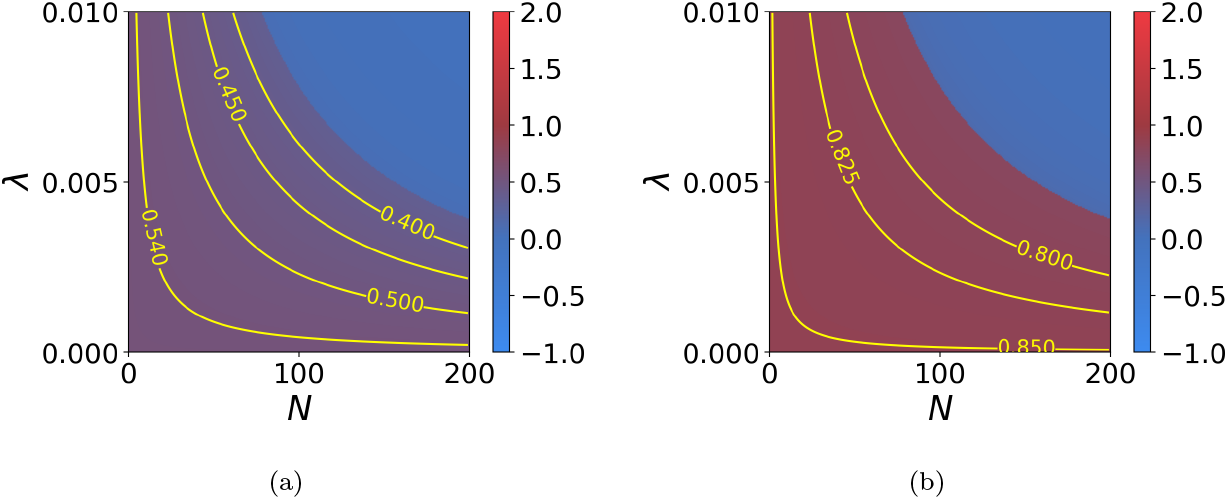
Influence of the density *N* and of the individual defence level λ in the *focal* species on the equilibrium values of (a) males trait 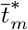 and (b) females trait 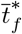 when female-limited mimicry is caused by sexually contrasted predation (*d_f_* > *d_m_*, *a* = 0). Yellow lines indicate equal trait value. We assume: *G_t_m__* = *G_t_f__* = *G_p_f__* = 0.01, *G_t_m_t_f__* = 0.001, *c_RI_* = 0, *c* = 0, *a* = 0, *b* = 5, *d_m_* = 0.01, *d_f_* = 0.05, λ′ = 0.01, *N*′ = 200, *s* = 0.01, *t_a_* = 0, *t*′ = 1.

**Figure A22:**
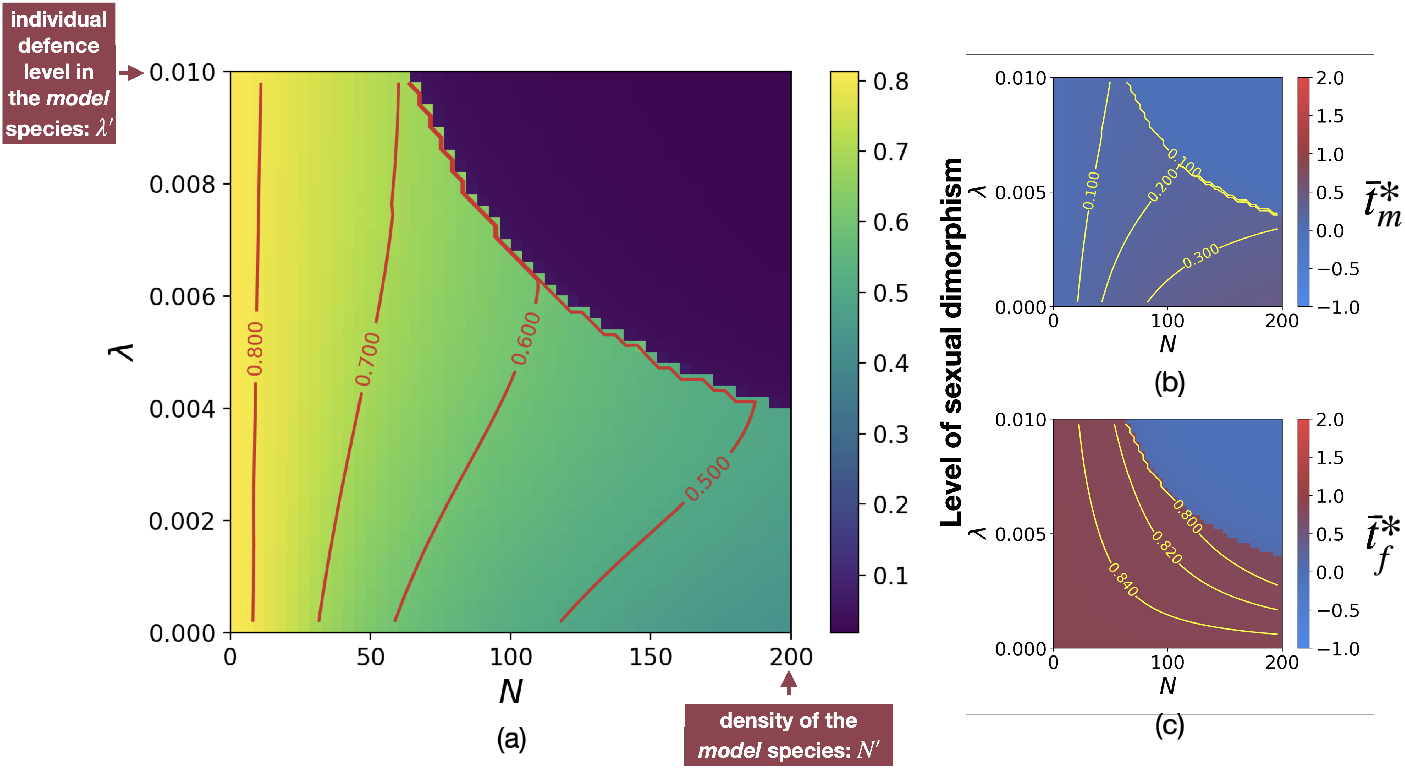
Influence of the density *N* and of the individual defence level λ in the *focal* species on the equilibrium values of (a) the level of sexual dimorphism 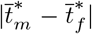, (b) males trait 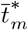 and (c) females trait 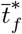 when female-limited mimicry is generated by sexual selection caused by reproductive interference (*c_RI_*, *a* > 0 and *d_f_* = *d_m_*). Red and yellow lines indicate equal levels of sexual dimorphism and trait value respectively. We assume: *G_t_m__* = *G_t_f__* = *G_p_f__* = 0.01, *G_t_m_t_f__* = 0.001, *c_RI_* = 0.01, *c* = 0.1, *a* = 5, *b* = 5, *d_m_* = *d_f_* = 0.05, λ′ = 0.01, *N*′ = 200, *s* = 0.01, *t_a_* = 0, *t*′ = 1.

**Figure A23:**
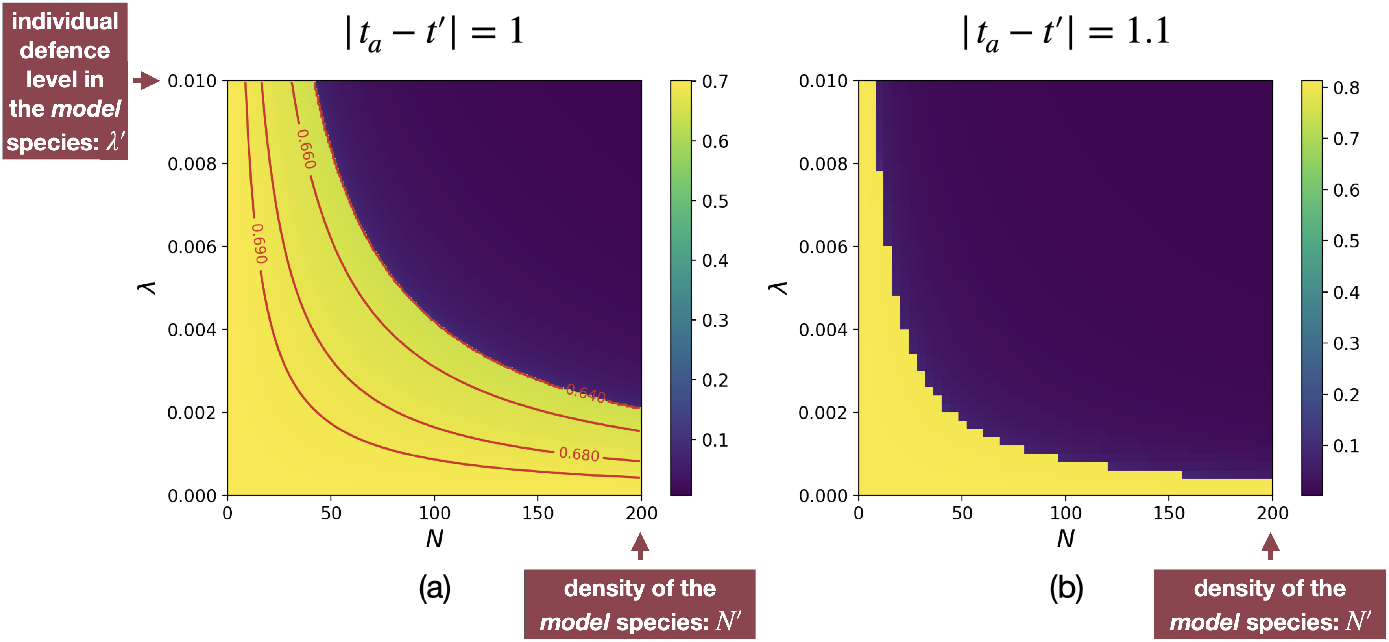
Influence of the density *N* and of the individual defence level λ in the *focal* species on the equilibrium values of the level of sexual dimorphism 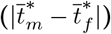 for different distances between the ancestral and the mimetic traits ((a) |*t_a_* – *t*′| = 1 (b) |*t_a_* – *t*′| = 1.1) when female-limited mimicry is caused by sexually contrasted predation (*d_f_* > *d_m_*, *a* = 0). Red lines indicate equal levels of sexual dimorphism. We assume: *G_t_m__* = *G_t_f__* = *G_p_f__* = 0.01, *G_t_m_t_f__* = 0.001, *c_RI_* = 0, *c* = 0, *a* = 0, *b* = 5, *d_m_* = 0.01, *d_f_* = 0.05, λ′ = 0.01, *N*′ = 200, *s* = 0.02, *t*′ = 1.

### 13 Investigation of the effect of defence level on the evolution of FLM using individual-centred simulations

#### 13.1 FLM caused by sexually constrasted predation

**Figure A24:**
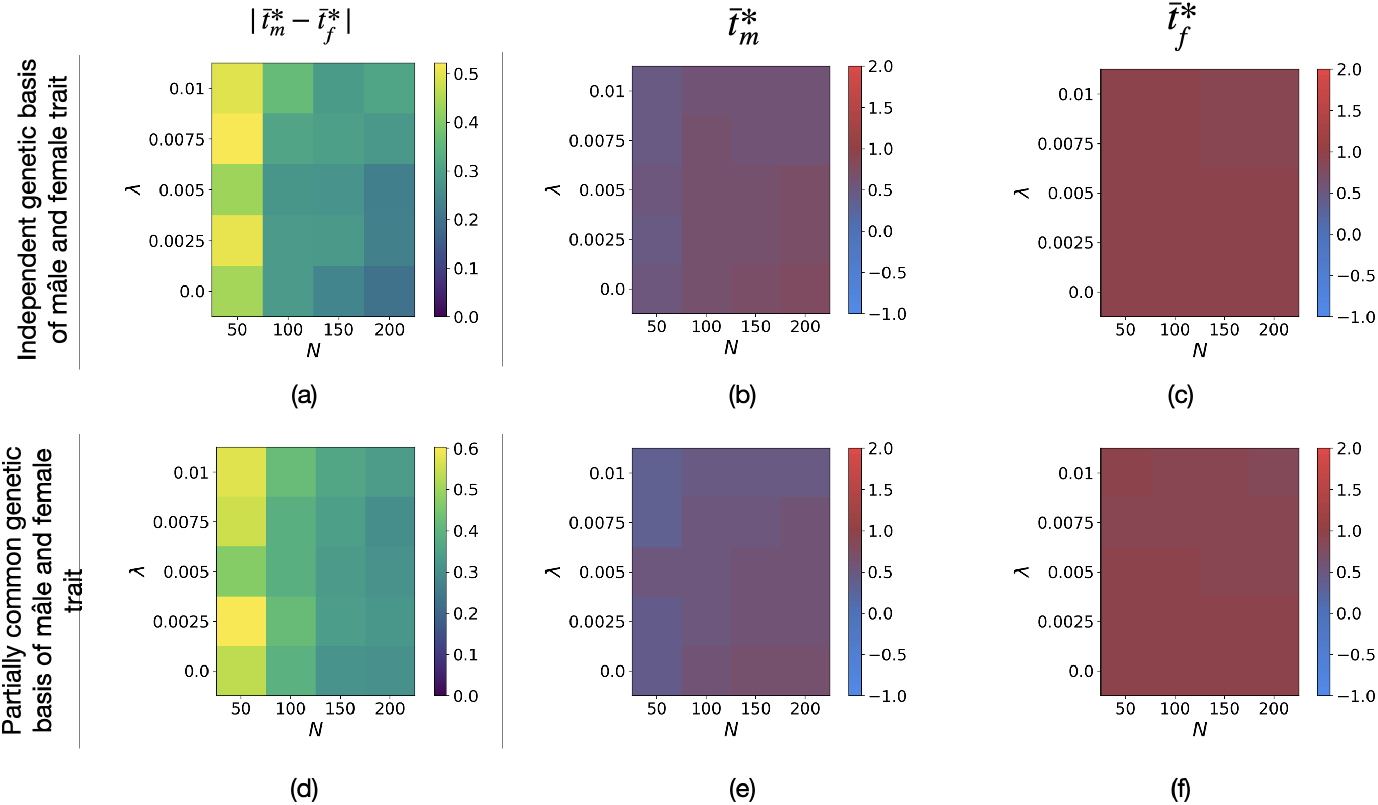
Influence of the density *N* and of the individual defence level λ in the *focal* species on the equilibrium values of (a)(d) the level of sexual dimorphism 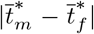, (b)(e) males trait 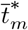 and (c)(f) females trait 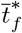 when selective constraints are low using individual-centred simulations assuming either (a)(b)(c) independent genetic basis or (d)(e)(f) partially common genetic basis of male and female trait. We assume: (a) *r_T_m_T_f__* = 0.25, *r_T_f_ P_f__* = 0.25 and (b) *r*_*T*_1_ *T*_2__ = 0.25, r_*T*_2_*T*_3__ = 0.25, *r*_*T*_3_*P_f_*_ = 0.25. We also assume: *G*_0_ = 0.0025, *μ* = 0.05, *c* = 0, *a* = 0, *c_RI_* = 0, *b* = 5, *d_m_* = 0.1, *d_f_* = 0.5, λ′ = 0.01, *N*′ = 200, *s* = 0.05, *t_a_* = 0, 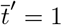.

**Figure A25:**
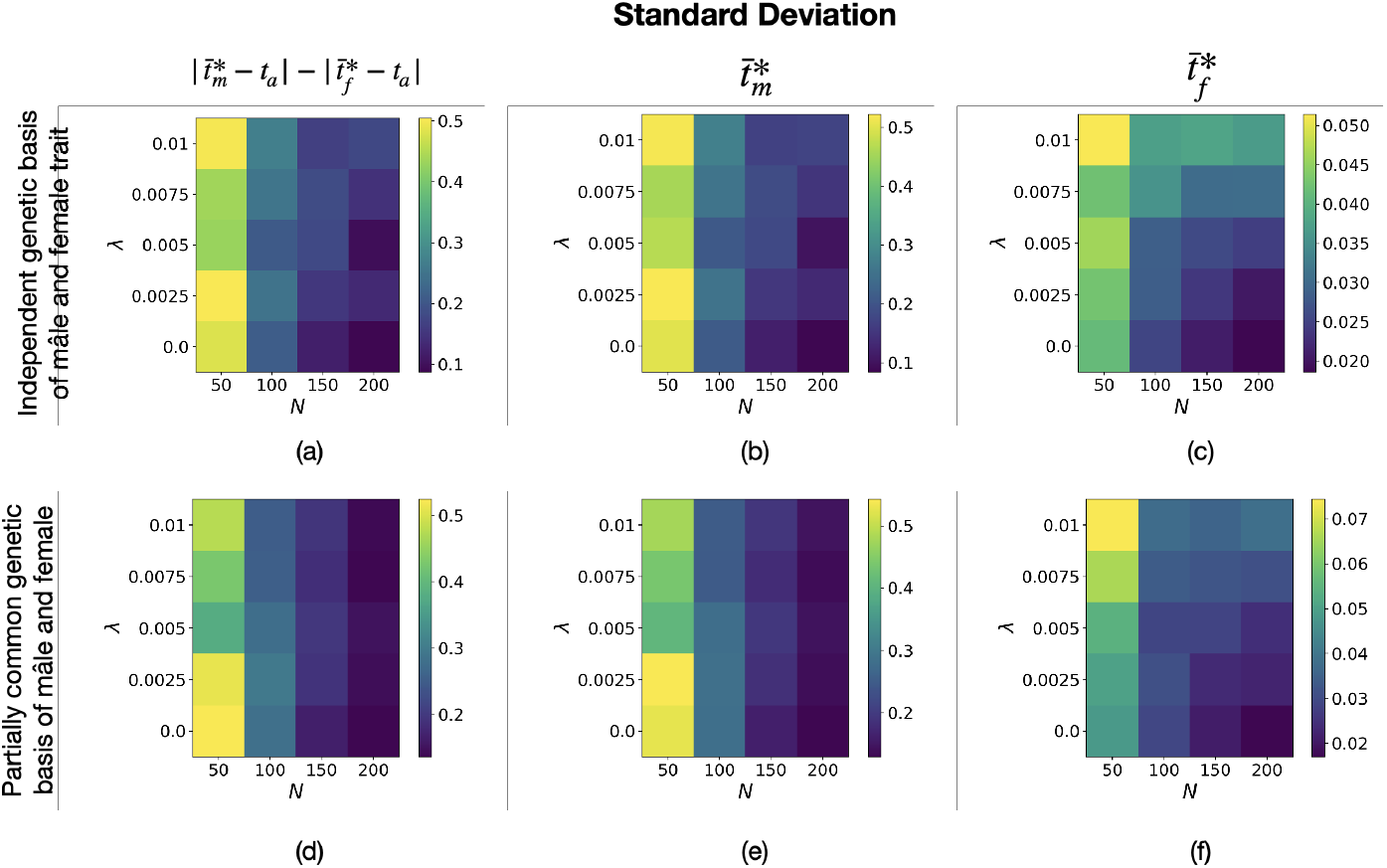
Standard deviation associated with Figure A24 of (a)(d) the level of sexual dimorphism 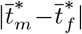, (b)(e) final male trait 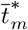 and (c)(f) final female trait 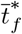 when selective constraints are low using individual-centred simulations assuming either (a)(b)(c) independent genetic basis or (d)(e)(f) partially common genetic basis of male and female trait.

**Figure A26:**
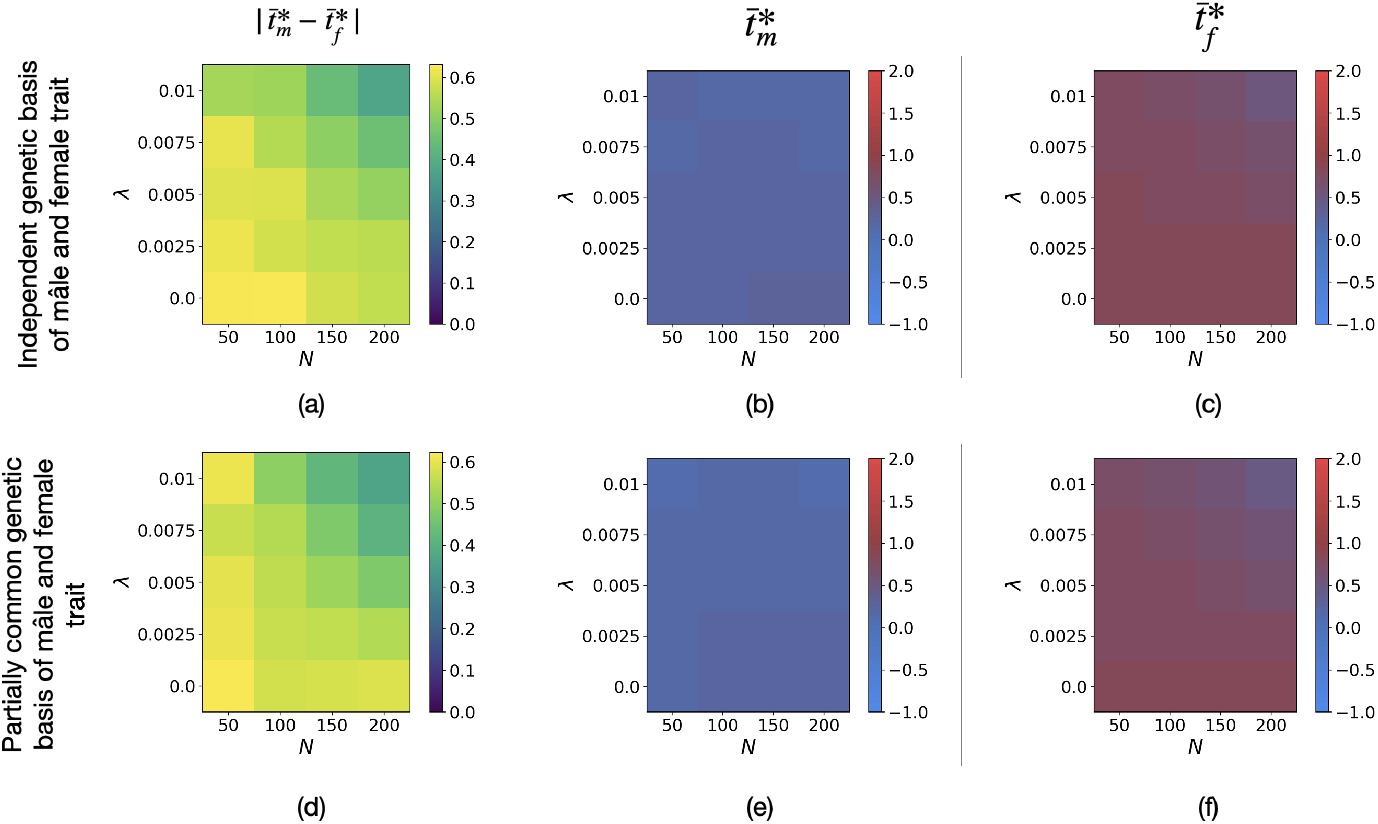
Influence of the density *N* and of the individual defence level λ in the *focal* species on the equilibrium values of (a)(d) the level of sexual dimorphism 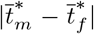, (b)(e) males trait 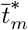 and (c)(f) females trait 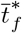 when selective constraints are high using individual-centred simulations assuming either (a)(b)(c) independent genetic basis or (d)(e)(f) partially common genetic basis of male and female trait. We assume: (a) *r_T_m_T_f__* = 0.25, *r_T_f_ P_f__* = 0.25 and (b) *r*_*T*_1_*T*_2__ = 0.25, *r*_*T*_2_*T*_3__ = 0.25, *r*_*T*_3_*P_f_*_ = 0.25. We also assume: *G*_0_ = 0.0025, *μ* = 0.05, *c* = 0, *a* = 0, *c_RI_* = 0, *b* = 5, *d_m_* = 0.1, *d_f_* = 0.5, λ′ = 0.01, *N*′ = 200, *s* = 0.1, *t_a_* = 0, 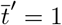.

**Figure A27:**
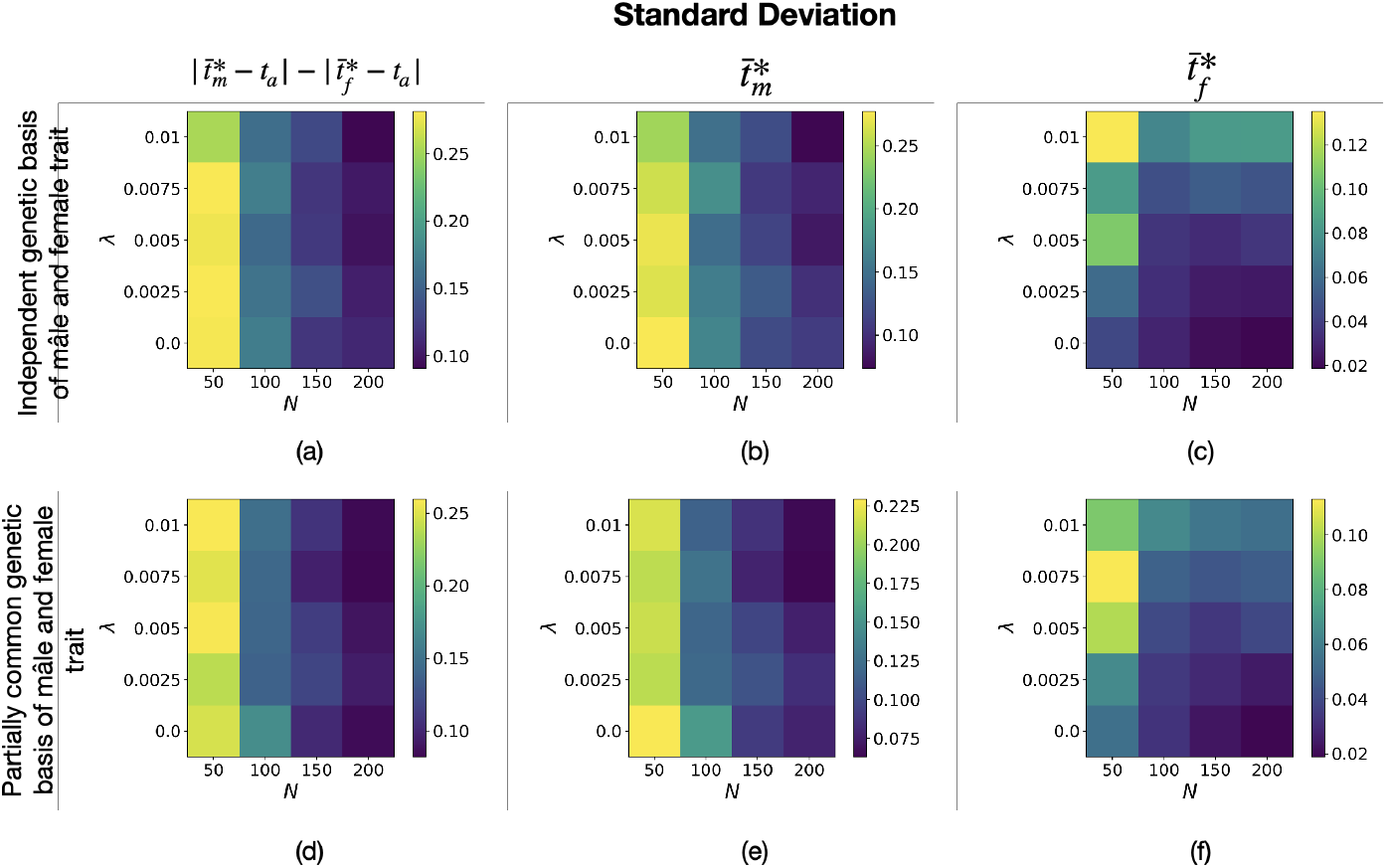
Standard deviation associated with Figure A26 of (a)(d) the level of sexual dimorphism 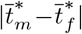, (b)(e) final male trait 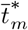 and (c)(f) final female trait 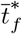 when selective constraints are high using individual-centred simulations assuming either (a)(b)(c) independent genetic basis or (d)(e)(f) partially common genetic basis of male and female trait.

#### 13.2 FLM caused by reproductive interference

**Figure A28:**
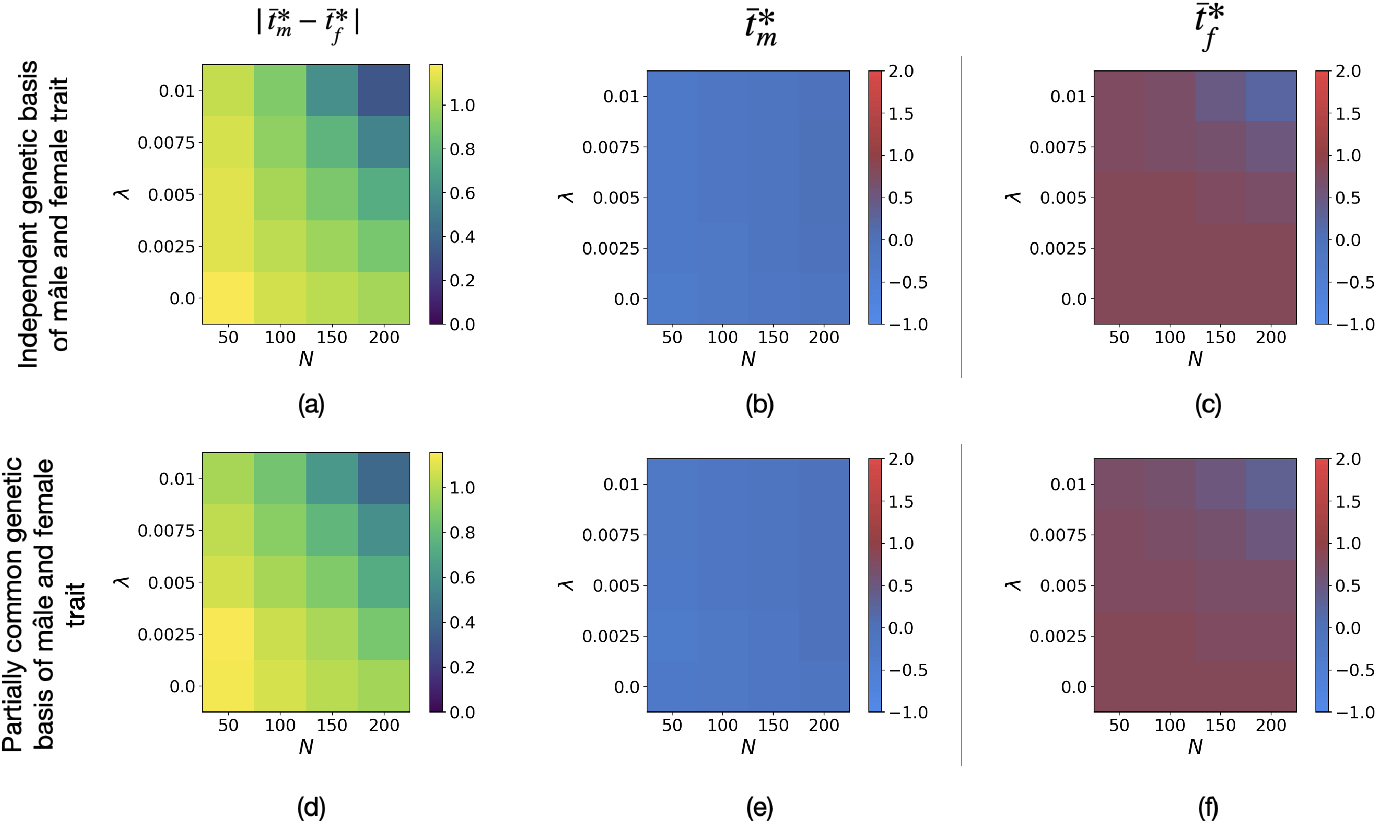
Influence of the density *N* and of the individual defence level λ in the *focal* species on the equilibrium values of (a)(d) the level of sexual dimorphism 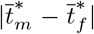, (b)(e) males trait 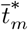 and (c)(f) females trait 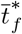 when selective constraints are high using individual-centred simulations, assuming either (a)(b)(c) independent genetic basis or (d)(e)(f) partially common genetic basis of male and female trait. We assume: (a) *r_T_m_T_f__* = 0.25, *r_T_f_ P_f__* = 0.25 and (b) *r*_*T*_1_ *T*_2__ = 0.25, *r*_*T*_2_*T*_3__ = 0.25, *r*_*T*_3_*P_f_*_ = 0.25. We also assume: *G*_0_ = 0.0025, *μ* = 0.05, *c* = 0.1, *a* = 5, *c_RI_* = 0.5, *b* = 5, *d_m_* = 0.5, *d_f_* = 0.5, λ′ = 0.01, *N*′ = 200, *s* = 0.1, *t_a_* = 0, 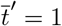.

**Figure A29:**
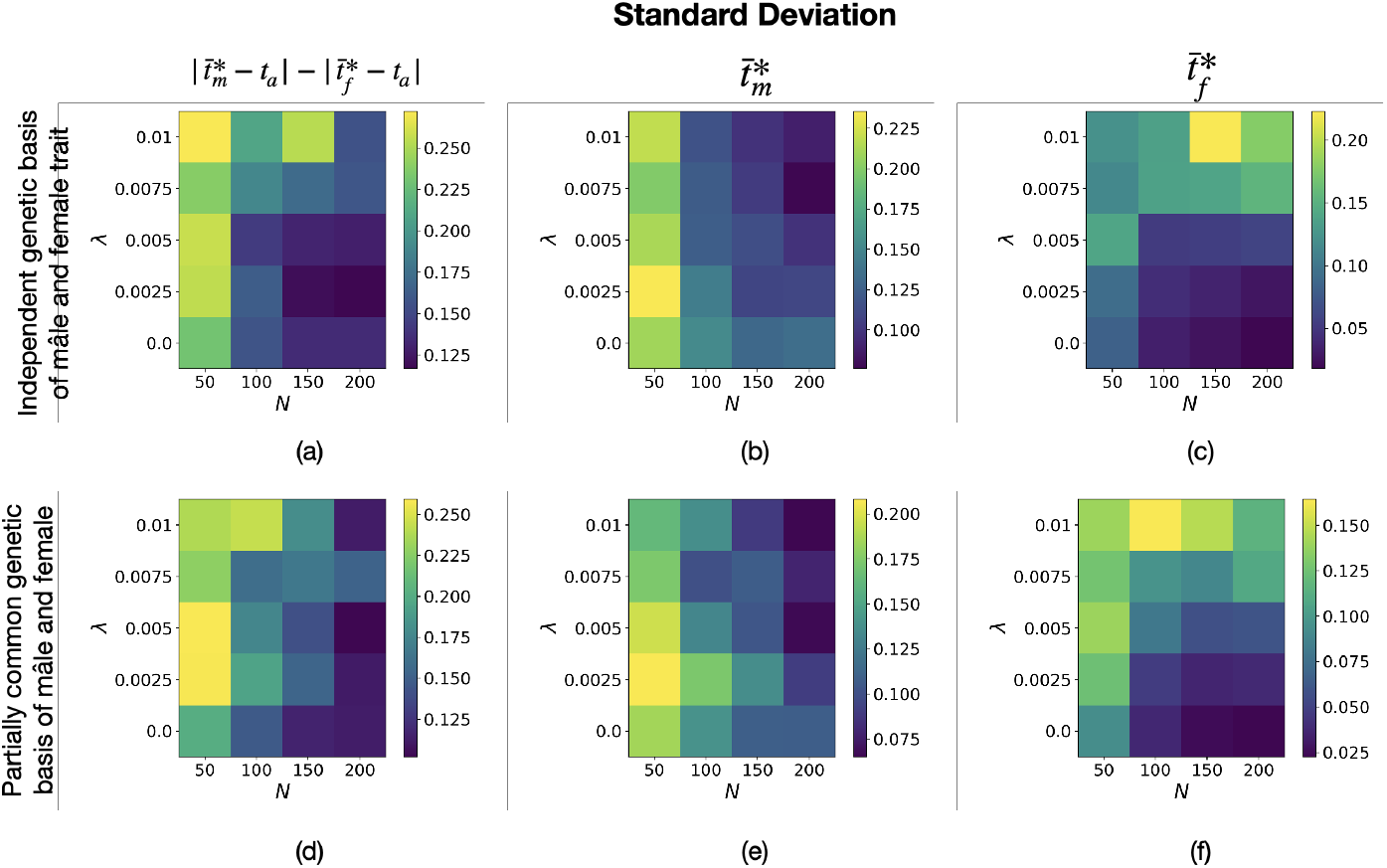
Standard deviation associated with Figure A28 of (a)(d) the level of sexual dimorphism 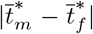, (b)(e) final male trait 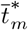 and (c)(f) final female trait 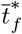 when selective constraints are high, using individual-centred simulations assuming either (a)(b)(c) independent genetic basis or (d)(e)(f) partially common genetic basis of male and female trait.

## References

S. Balshine, B. Kempenaers, T. Székely, H. Kokko, and R. A. Johnstone. Why is mutual mate choice not the norm? operational sex ratios, sex roles and the evolution of sexually dimorphic and monomorphic signalling. Philosophical Transactions of the Royal Society of London. Series B: Biological Sciences, 357(1419):319–330, 2002. 10.1098/rstb.2001.0926. URL https://royalsocietypublishing.org/doi/abs/10.1098/rstb.2001.0926.

N. H. Barton and M. Turelli. Natural and sexual selection on many loci. Genetics, 127(1):229–255, 1991. ISSN 0016-6731. URL https://www.genetics.org/content/127/1/229.

H. W. Bates. Contributions to an insect fauna of the amazon valley (lepidoptera: Heliconidae). Biological Journal of the Linnean Society, 16(1):41–54, 1981. https://doi.org/10.1111/j.1095-8312.1981.tb01842.x. URL https://onlinelibrary.wiley.com/doi/abs/10.1111/j.1095-8312.1981.tb01842.x.

J. Baur, A. Giesen, P. T. Rohner, W. U. Blanckenhorn, and M. A. Schäfer. Exaggerated male forelegs are not more differentiated than wing morphology in two widespread sister species of black scavenger flies. Journal of Zoological Systematics and Evolutionary Research, 58(1):159–173, 2020. https://doi.org/10.1111/jzs.12327. URL https://onlinelibrary.wiley.com/doi/abs/10.1111/jzs.12327.

T. Belt. The naturalist in Nicaragua: A narrative of a residence at the gold mines of Chontales; journeys in the savannahs and forests. With observations on animals and plants in reference to the theory of evolution of living forms. London, J. Murray,, 1874. URL https://www.biodiversitylibrary.org/item/16263. https://www.biodiversitylibrary.org/bibliography/1390.

W. W. Benson. Natural selection for mullerian mimicry in heliconius erato in costa rica. Science, 176(4037):936–939, 1972. ISSN 00368075, 10959203. URL http://www.jstor.org/stable/1733812.

C. L. Boggs and L. E. Gilbert. Male contribution to egg production in butterflies: Evidence for transfer of nutrients at mating. Science, 206(4414):83–84, 1979. ISSN 0036-8075. 10.1126/science.206.4414.83. URL https://science.sciencemag.org/content/206/4414/83.

G. Boussens-Dumon and V. Llaurens. Sex, competition and mimicry: an eco-evolutionary model reveals unexpected impacts of ecological interactions on the evolution of phenotypes in sympatry. Oikos, 130(11):2028–2039, 2021. https://doi.org/10.1111/oik.08139. URL https://onlinelibrary.wiley.com/doi/abs/10.1111/oik.08139.

P. L. Brennan, C. J. Clark, and R. O. Prum. Explosive eversion and functional morphology of the duck penis supports sexual conflict in waterfowl genitalia. Proceedings of the Royal Society B: Biological Sciences, 277(1686):1309–1314, 2010.

P. Chai and R. B. Srygley. Predation and the flight, morphology, and temperature of neotropical rain-forest butterflies. The American Naturalist, 135(6):748–765, 1990. ISSN 00030147, 15375323. URL http://www.jstor.org/stable/2462312.

M. Chouteau, M. Arias, and M. Joron. Warning signals are under positive frequency-dependent selection in nature. Proceedings of the National Academy of Sciences, 113(8):2164–2169, 2016. ISSN 0027-8424. 10.1073/pnas.1519216113. URL https://www.pnas.org/content/113/8/2164.

K. Darragh, S. Vanjari, F. Mann, M. F. Gonzalez-Rojas, C. R. Morrison, C. Salazar, C. Pardo-Diaz, R. M. Merrill, W. O. McMillan, S. Schulz, and C. D. Jiggins. Male sex pheromone components in *Heliconius* butterflies released by the androconia affect female choice. PeerJ, 5:e3953, Nov. 2017. ISSN 2167-8359. 10.7717/peerj.3953. URL https://doi.org/10.7717/peerj.3953.

C. Darwin. The descent of man and se1ection inre1ation to sex. The origin of species and the descent of man and selection in relation, 1871.

C. Estrada and C. Jiggins. Patterns of pollen feeding and habitat preference among heliconius species. Ecological Entomology, 27:448 – 456, 08 2002. 10.1046/j.1365-2311.2002.00434.x.

C. Estrada and C. D. Jiggins. Interspecific sexual attraction because of convergence in warning colouration: is there a conflict between natural and sexual selection in mimetic species? Journal of Evolutionary Biology, 21(3):749–760, 2008. https://doi.org/10.1111/j.1420-9101.2008.01517.x. URL https://onlinelibrary.wiley.com/doi/abs/10.1111/j.1420-9101.2008.01517.x.

E. Ford. Ecological Genetics. Springer Netherlands, 1975. ISBN 9789400958258. URL https://www.springer.com/gp/book/9780412161308.

G. W. Gilchrist. The consequences of sexual dimorphism in body size for butterfly flight and thermoregulation. Functional Ecology, 4(4):475–487, 1990. ISSN 02698463, 13652435. URL http://www.jstor.org/stable/2389315.

M. F. González-Rojas, K. Darragh, J. Robles, M. Linares, S. Schulz, W. O. McMillan, C. D. Jiggins, C. Pardo-Diaz, and C. Salazar. Chemical signals act as the main reproductive barrier between sister and mimetic ¡i¿heliconius¡/i¿ butterflies. Proceedings of the Royal Society B: Biological Sciences, 287(1926):20200587, 2020. 10.1098/rspb.2020.0587. URL https://royalsocietypublishing.org/doi/abs/10.1098/rspb.2020.0587.

J. Gröning and A. Hochkirch. Reproductive interference between animal species. The Quarterly Review of Biology, 83(3):257–282, 2008. 10.1086/590510. URL https://doi.org/10.1086/590510. PMID: 18792662.

Y. Iwasa, A. Pomiankowski, and S. Nee. The evolution of costly mate preferences ii. the “handicap” principle. Evolution, 45(6):1431–1442, 1991. 10.1111/j.1558-5646.1991.tb02646.x. URL https://onlinelibrary.wiley.com/doi/abs/10.1111/j.1558-5646.1991.tb02646.x.

C. D. Jiggins, R. E. Naisbit, R. L. Coe, and J. Mallet. Reproductive isolation caused by colour pattern mimicry. Nature, 411(6835):302–305, 2001. 10.1038/35077075. URL https://doi.org/10.1038/35077075.

R. A. Johnstone. Sexual selection, honest advertisement and the handicap principle: reviewing the evidence. Biological Reviews, 70(1):1–65, 1995. https://doi.org/10.1111/j.1469-185X.1995.tb01439.x. URL https://onlinelibrary.wiley.com/doi/abs/10.1111/j.1469-185X.1995.tb01439.x.

B. Karlsson and P.-O. Wickman. Increase in reproductive effort as explained by body size and resource allocation in the speckled wood butterfly, pararge aegeria (l.). Functional Ecology, 4(5):609–617, 1990. ISSN 02698463, 13652435. URL http://www.jstor.org/stable/2389728.

M. Katoh, H. Tatsuta, and K. Tsuji. Mimicry genes reduce pre-adult survival rate in papilio polytes: A possible new mechanism for maintaining female-limited polymorphism in batesian mimicry. Journal of Evolutionary Biology, 33(10):1487–1494, 2020. https://doi.org/10.1111/jeb.13686. URL https://onlinelibrary.wiley.com/doi/abs/10.1111/jeb.13686.

D. W. Kikuchi and D. W. Pfennig. High-model abundance may permit the gradual evolution of batesian mimicry: an experimental test. Proceedings of the Royal Society B: Biological Sciences, 277(1684):1041–1048, 2010. 10.1098/rspb.2009.2000. URL https://royalsocietypublishing.org/doi/abs/10.1098/rspb.2009.2000.

M. Kirkpatrick, T. Johnson, and N. Barton. General models of multilocus evolution. Genetics, 161(4):1727–1750, 2002. ISSN 0016-6731. URL https://www.genetics.org/content/161/4/1727.

S. Komata, C.-P. Lin, and T. Sota. Do juvenile developmental and adult body characteristics differ among genotypes at the doublesex locus that controls female-limited batesian mimicry polymorphism in papilio memnon?: A test for the “cost of mimicry” hypothesis. Journal of Insect Physiology, 107:1–6, 2018. ISSN 0022-1910. https://doi.org/10.1016/j.jinsphys.2018.02.001. URL https://www.sciencedirect.com/science/article/pii/S0022191017304420.

S. Komata, T. Kitamura, and H. Fujiwara. Batesian mimicry has evolved with deleterious effects of the pleiotropic gene doublesex. Scientific Reports, 10(1):21333, 2020.

R. A. Krebs and D. A. West. Female mate preference and the evolution of female-limited batesian mimicry. Evolution, 42(5):1101–1104, 1988. ISSN 00143820, 15585646. URL http://www.jstor.org/stable/2408927.

M. R. Kronforst, L. G. Young, D. D. Kapan, C. McNeely, R. J. O’Neill, and L. E. Gilbert. Linkage of butterfly mate preference and wing color preference cue at the genomic location of wingless. Proceedings of the National Academy of Sciences, 103(17):6575–6580, 2006. 10.1073/pnas.0509685103. URL https://www.pnas.org/content/103/17/6575.

K. Kunte. Mimetic butterflies support wallace’s model of sexual dimorphism. Proceedings of the Royal Society B: Biological Sciences, 275(1643):1617–1624, 2008. 10.1098/rspb.2008.0171. URL https://royalsocietypublishing.org/doi/abs/10.1098/rspb.2008.0171.

K. Kunte. The diversity and evolution of batesian mimicry in papilio swallowtail butterflies. Evolution, 63(10):2707–2716, 2009. 10.1111/j.1558-5646.2009.00752.x. URL https://onlinelibrary.wiley.com/doi/abs/10.1111/j.1558-5646.2009.00752.x.

C. Lamunyon. Increased fecundity, as a function of multiple mating, in an arctiid moth, utetheisa ornatrix. Ecological Entomology, 22(1):69–73, 1997. https://doi.org/10.1046/j.1365-2311.1997.00033.x. URL https://onlinelibrary.wiley.com/doi/abs/10.1046/j.1365-2311.1997.00033.x.

R. Lande. Models of speciation by sexual selection on polygenic traits. Proceedings of the National Academy of Sciences, 78(6):3721–3725, 1981. ISSN 0027-8424. 10.1073/pnas.78.6.3721. URL https://www.pnas.org/content/78/6/3721.

R. Lande and S. J. Arnold. Evolution of mating preference and sexual dimorphism. Journal of Theoretical Biology, 117(4):651–664, 1985. ISSN 0022-5193. https://doi.org/10.1016/S0022-5193(85)80245-9. URL https://www.sciencedirect.com/science/article/pii/S0022519385802459.

R. C. Lederhouse and J. M. Scriber. Intrasexual selection constrains the evolution of the dorsal color pattern of male black swallowtail butterflies, papilio polyxenes. Evolution, 50(2):717–722, 1996. https://doi.org/10.1111/j.1558-5646.1996.tb03881.x. URL https://onlinelibrary.wiley.com/doi/abs/10.1111/j.1558-5646.1996.tb03881.x.

L. Lindström, R. V. Alatalo, and J. Mappes. Imperfect batesian mimicry—the effects of the frequency and the distastefulness of the model. Proceedings of the Royal Society B: Biological Sciences, 264(1379):149–153, Feb 1997. ISSN 0962-8452. 10.1098/rspb.1997.0022. URL https://www.ncbi.nlm.nih.gov/pmc/articles/PMC1688248/. PMC1688248[pmcid].

E. C. Long, T. P. Hahn, and A. M. Shapiro. Variation in wing pattern and palatability in a female-limited polymorphic mimicry system. Ecology and evolution, 4(23):4543–4552, 12 2014.

E. C. Long, K. F. Edwards, and A. M. Shapiro. A test of fundamental questions in mimicry theory using long-term datasets. Biological Journal of the Linnean Society, 116(3):487–494, 2015. https://doi.org/10.1111/bij.12608. URL https://onlinelibrary.wiley.com/doi/abs/10.1111/bij.12608.

X. H. Low and A. Monteiro. Dorsal forewing white spots of male papilio polytes(lepidoptera: Papilionidae) not maintained by female mate choice. Journal of Insect Behavior, 31(1):29–41, 2018.

L. Maisonneuve, C. Smadi, and V. Llaurens. The limits of evolutionary convergence in sympatry: reproductive interference and developmental constraints leading to local diversity in aposematic signals. bioRxiv, 2021. 10.1101/2021.01.22.427743. URL https://www.biorxiv.org/content/early/2021/01/22/2021.01.22.427743.

J. Mallet and N. H. Barton. Strong natural selection in a warning-color hybrid zone. Evolution, 43(2):421–431, 1989. ISSN 00143820, 15585646. URL http://www.jstor.org/stable/2409217.

J. Mallet and M. Joron. Evolution of diversity in warning color and mimicry: Polymorphisms, shifting balance, and speciation. Annual Review of Ecology and Systematics, 30(1):201–233, 1999. 10.1146/annurev.ecolsys.30.1.201. URL https://doi.org/10.1146/annurev.ecolsys.30.1.201.

J. H. Marden and P. Chai. Aerial predation and butterfly design: How palatability, mimicry, and the need for evasive flight constrain mass allocation. The American Naturalist, 138(1):15–36, 1991. ISSN 00030147, 15375323. URL http://www.jstor.org/stable/2462530.

S. Marsteller, D. C. Adams, M. L. Collyer, and M. Condon. Six cryptic species on a single species of host plant: morphometric evidence for possible reproductive character displacement. Ecological Entomology, 34(1):66–73, 2009. https://doi.org/10.1111/j.1365-2311.2008.01047.x. URL https://onlinelibrary.wiley.com/doi/abs/10.1111/j.1365-2311.2008.01047.x.

M. A. McPeek and S. Gavrilets. The evolution of female mating preferences: Differentiation from species with promiscious males can promotes speciation. Evolution, 60(10):1967 – 1980, 2006. 10.1554/06-184.1. URL https://doi.org/10.1554/06-184.1.

R. M. Merrill, A. Chia, and N. J. Nadeau. Divergent warning patterns contribute to assortative mating between incipient heliconius species. Ecology and Evolution, 4(7):911–917, 2014. https://doi.org/10.1002/ece3.996. URL https://onlinelibrary.wiley.com/doi/abs/10.1002/ece3.996.

C. Mérot, B. Frérot, E. Leppik, and M. Joron. Beyond magic traits: Multimodal mating cues in heliconius butterflies. Evolution, 69(11):2891–2904, 2015. https://doi.org/10.1111/evo.12789. URL https://onlinelibrary.wiley.com/doi/abs/10.1111/evo.12789.

T. Nagylaki. The evolution of multilocus systems under weak selection. Genetics, 134(2):627–647, 1993. ISSN 0016-6731. URL https://www.genetics.org/content/134/2/627.

R. E. Naisbit, C. D. Jiggins, and J. Mallet. Disruptive sexual selection against hybrids contributes to speciation between ¡i¿heliconius cydno¡/i¿ and ¡i¿heliconius melpomene¡/i¿. Proceedings of the Royal Society of London. Series B: Biological Sciences, 268(1478):1849–1854, 2001. 10.1098/rspb.2001.1753. URL https://royalsocietypublishing.org/doi/abs/10.1098/rspb.2001.1753.

R. Nishida. Chemical Ecology of Poisonous Butterflies: Model or Mimic? A Paradox of Sexual Dimorphisms in Müllerian Mimicry, pages 205–220. Springer Singapore, Singapore, 2017. ISBN 978-981-10-4956-9. 10.1007/978-981-10-4956-9_11. URL https://doi.org/10.1007/978-981-10-4956-9_11.

H. Nishikawa, T. Iijima, R. Kajitani, J. Yamaguchi, T. Ando, Y. Suzuki, S. Sugano, A. Fujiyama, S. Kosugi, H. Hirakawa, S. Tabata, K. Ozaki, H. Morimoto, K. Ihara, M. Obara, H. Hori, T. Itoh, and H. Fujiwara. A genetic mechanism for female-limited batesian mimicry in papilio butterfly. Nature Genetics, 47(4):405–409, 2015. 10.1038/ng.3241. URL https://doi.org/10.1038/ng.3241.

N. Ohsaki. Preferential predation of female butterflies and the evolution of batesian mimicry. Nature, 378(6553):173–175, Nov 1995. ISSN 1476-4687. 10.1038/378173a0. URL https://doi.org/10.1038/378173a0.

N. Ohsaki. A common mechanism explaining the evolution of female-limited and both-sex batesian mimicry in butterflies. Journal of Animal Ecology, 74(4):728–734, 2005. https://doi.org/10.1111/j.1365-2656.2005.00972.x. URL https://besjournals.onlinelibrary.wiley.com/doi/abs/10.1111/j.1365-2656.2005.00972.x.

S. P. Otto, M. R. Servedio, and S. L. Nuismer. Frequency-dependent selection and the evolution of assortative mating. Genetics, 179(4):2091–2112, 2008. ISSN 0016-6731. 10.1534/genetics.107.084418. URL https://www.genetics.org/content/179/4/2091.

G. Parker. Sexual conflict over mating and fertilization: an overview. Philosophical Transactions of the Royal Society B: Biological Sciences, 361(1466):235–259, 2006. 10.1098/rstb.2005.1785. URL https://royalsocietypublishing.org/doi/abs/10.1098/rstb.2005.1785.

K. S. Pfennig and D. W. Pfennig. CHARACTER DISPLACEMENT AS THE “BEST OF A BAD SITUATION”: FITNESS TRADE-OFFS RESULTING FROM SELECTION TO MINIMIZE RESOURCE AND MATE COMPETITION. Evolution, 59(10):2200 – 2208, 2005. 10.1554/05-263.1. URL https://doi.org/10.1554/05-263.1.

A. Pomiankowski and Y. Iwasa. Evolution of multiple sexual preferences by fisher’s runaway process of sexual selection. Proceedings: Biological Sciences, 253(1337):173–181, 1993. ISSN 09628452. URL http://www.jstor.org/stable/49806.

K. L. Prudic, B. N. Timmermann, D. R. Papaj, D. B. Ritland, and J. C. Oliver. Mimicry in viceroy butterflies is dependent on abundance of the model queen butterfly. Communications Biology, 2(1):68, Feb 2019. ISSN 2399-3642. 10.1038/s42003-019-0303-z. URL https://doi.org/10.1038/s42003-019-0303-z.

L. A. Prusa and R. I. Hill. Umbrella of protection: spatial and temporal dynamics in a temperate butterfly Batesian mimicry system. Biological Journal of the Linnean Society, 04 2021. ISSN 0024-4066. 10.1093/biolinnean/blab004. URL https://doi.org/10.1093/biolinnean/blab004.blab004.

S. H. Rice. Evolutionary theory: mathematical and conceptual foundations. Sinauer Associates, Sunderland, Mass., USA, 2004. ISBN 9780878937028.

H. M. Rowland, J. Mappes, G. D. Ruxton, and M. P. Speed. Mimicry between unequally defended prey can be parasitic: evidence for quasi-batesian mimicry. Ecology Letters, 13(12):1494–1502, 2010. https://doi.org/10.1111/j.1461-0248.2010.01539.x. URL https://onlinelibrary.wiley.com/doi/abs/10.1111/j.1461-0248.2010.01539.x.

G. Ruxton, W. Allen, T. Sherratt, and M. Speed. Avoiding Attack: The Evolutionary Ecology of Crypsis, Aposematism, and Mimicry. OUP Oxford, 2019. ISBN 9780191002632. URL https://books.google.fr/books?id=SiKFDwAAQBAJ.

K. Sanders, A. Malhotra, and R. Thorpe. Evidence for a müllerian mimetic radiation in asian pitvipers. Proceedings of the Royal Society B: Biological Sciences, 273(1590):1135–1141, 2006. 10.1098/rspb.2005.3418. URL https://royalsocietypublishing.org/doi/abs/10.1098/rspb.2005.3418.

O. Sculfort, E. C. P. de Castro, K. M. Kozak, S. Bak, M. Elias, B. Nay, and V. Llaurens. Variation of chemical compounds in wild heliconiini reveals ecological factors involved in the evolution of chemical defenses in mimetic butterflies. Ecology and Evolution, 10(5):2677–2694, 2020. https://doi.org/10.1002/ece3.6044. URL https://onlinelibrary.wiley.com/doi/abs/10.1002/ece3.6044.

T. N. Sherratt. The evolution of müllerian mimicry. Die Naturwissenschaften, 95(8):681–695, 08 2008. 10.1007/s00114-008-0403-y. URL https://pubmed.ncbi.nlm.nih.gov/18542902.

S. Su, M. Lim, and K. Kunte. Prey from the eyes of predators: Color discriminability of aposematic and mimetic butterflies from an avian visual perspective. Evolution, 69(11):2985–2994, 2015. https://doi.org/10.1111/evo.12800. URL https://onlinelibrary.wiley.com/doi/abs/10.1111/evo.12800.

M. J. T. N. Timmermans, M. J. Thompson, S. Collins, and A. P. Vogler. Independent evolution of sexual dimorphism and female-limited mimicry in swallowtail butterflies (papilio dardanus and papilio phorcas). Mol Ecol, 26(5):1273–1284, Feb. 2017.

R. Trivers. Parental Investment and Sexual Selection, page 378. 01 1972.

J. R. G. Turner. Why male butterflies are non-mimetic: natural selection, sexual selection, group selection, modification and sieving*. Biological Journal of the Linnean Society, 10(4):385–432, 1978. https://doi.org/10.1111/j.1095-8312.1978.tb00023.x. URL https://onlinelibrary.wiley.com/doi/abs/10.1111/j.1095-8312.1978.tb00023.x.

S. M. Van Belleghem, P. A. Alicea Roman, H. Carbia Gutierrez, B. A. Counterman, and R. Papa. Perfect mimicry between ¡i¿heliconius¡/i¿ butterflies is constrained by genetics and development. Proceedings of the Royal Society B: Biological Sciences, 287(1931):20201267, 2020. 10.1098/rspb.2020.1267. URL https://royalsocietypublishing.org/doi/abs/10.1098/rspb.2020.1267.

W. van der Bijl, D. Zeuss, N. Chazot, K. Tunström, N. Wahlberg, C. Wiklund, J. L. Fitzpatrick, and C. W. Wheat. Butterfly dichromatism primarily evolved via darwin’s, not wallace’s, model. Evolution Letters, 4(6):545–555, 2020.

A. R. Wallace. On the phenomena of variation and geographical distribution as illustrated by the Papilionidae of the Malayan region. Read March 17, 1864. London,, 1865. URL https://www.biodiversitylibrary.org/item/38597. https://www.biodiversitylibrary.org/bibliography/9531.

C. Wiklund, A. Kaitala, V. Lindfors, and J. Abenius. Polyandry and its effect on female reproduction in the green-veined white butterfly (pieris napi l.). Behavioral Ecology and Sociobiology, 33(1):25–33, 1993. 10.1007/BF00164343. URL https://doi.org/10.1007/BF00164343.

M. K. Wourms and F. E. Wasserman. Bird predation on lepidoptera and the reliability of beak-marks in determining predation pressure. Journal of the Lepidopterists’ Society, 39(4):239–261, 1985. URL https://images.peabody.yale.edu/lepsoc/jls/1980s/1985/1985-39(4)239-Wourms.pdf.

R. Yamaguchi and Y. Iwasa. Reproductive character displacement by the evolution of female mate choice. Evolutionary Ecology Research, 15(1):25–41, 7 2013. ISSN 1522-0613.

A. Zahavi. Mate selection—a selection for a handicap. Journal of Theoretical Biology, 53(1):205–214, 1975. ISSN 0022-5193. https://doi.org/10.1016/0022-5193(75)90111-3. URL https://www.sciencedirect.com/science/article/pii/0022519375901113.

